# Integration of in situ hybridization and scRNA-seq data provides a 2D topographical map of the developing retina across species

**DOI:** 10.64898/2026.01.04.697548

**Authors:** Heer N. V. Joisher, ChangHee Lee, Chaitra Prabhakara, Isabella van der Weide, Yichen Si, Nicholas Lonfat, Constance Cepko

**Affiliations:** Department of Genetics, Blavatnik Institute, Harvard Medical School, United States; Department of Ophthalmology, Harvard Medical School, United States; Howard Hughes Medical Institute, United States; Somite Therapeutics, Boston, Massachusetts, United States; Broad Institute of MIT and Harvard, Cambridge, Massachusetts, United State; City Therapeutics, Boston, Massachusetts, United States

## Abstract

Precise regional patterning is fundamental to tissue organization, yet the spatial logic that governs it remains poorly defined for many tissues. In the vertebrate retina, molecular domains along the dorsoventral and nasotemporal axes provide positional cues for regional specializations such as the high-acuity area (HAA). We combined multiplexed in situ hybridization data with single-cell transcriptomic data to create quantitative two-dimensional maps of developing retinal cells. In the developing chicken retina, this approach resolved sharp expression boundaries of genes involved in patterning, and revealed novel candidates enriched in the anlagen of the HAA. Comparative analysis of chicken, mouse, and human data demonstrated conserved axis-based programs, but distinct fine-scale organization consistent with presence/absence of an HAA. Here, we show that spatial reconstruction from scRNA-seq data, anchored by experimental benchmarks, enables comparative 2D topographic mapping of gene expression across species and provides a generalizable strategy to investigate the spatial logic of molecular organization in developing tissues.

**GRAPHICAL ABSTRACT:** 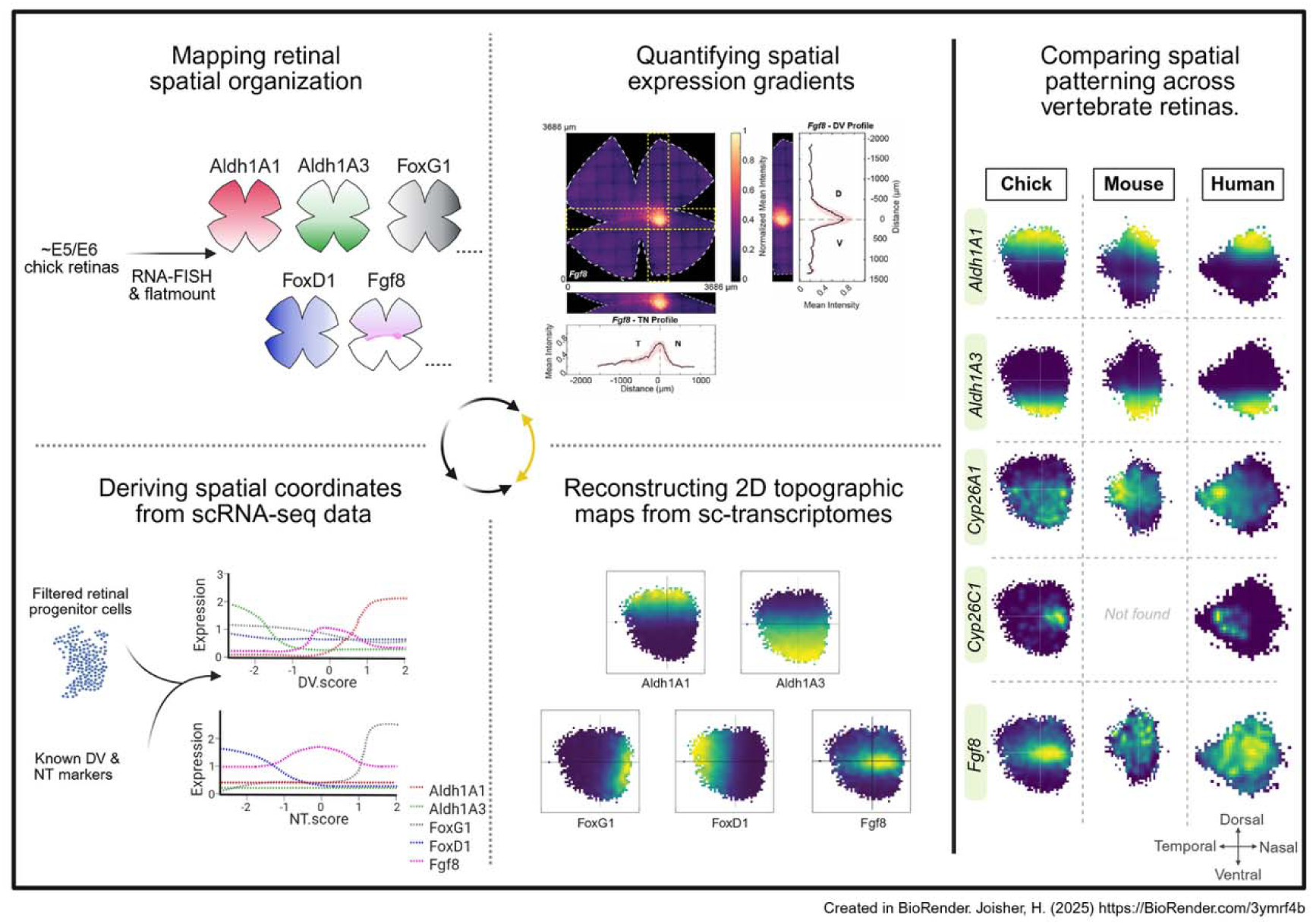

**HIGHLIGHTS:**

- Quantification of multiplexed RNA-FISH signals generated reproducible spatial expression maps in the developing chicken retina.
- Resolution of relative boundaries for early patterning genes revealed nested expression domains marking the boundaries of the developing HAA.
- RNA-FISH experimental data using genes with axis-specific expression allowed the generation of DV and NT scores which enabled the localization of each cell in a scRNA-seq dataset onto a 2D topographical map of the developing retina.
- 2D topographic maps derived from scRNA-seq data revealed novel HAA-enriched genes in the developing chicken retina.
- Cross-species comparisons showed conserved DV/NT programs but divergent fine-scale organization in retinal development.
- *Cyp26c1* expression pattern was validated in human retina correlating with the location of the future HAA.

## INTRODUCTION

Precise spatial patterning is fundamental to the development of complex tissues, ensuring that specific functional domains are properly localized and specified. During embryogenesis, localized cues orchestrate regional identity across tissue types and species (57,36,11,16,17,35,19,28,29). Tissues establish regional identity by interpreting positional information encoded in gradients, boundaries, and lineage constraints. The vertebrate eye is no exception to the principle of regional patterning. The retina is highly organized, with precise lamination and 2D topographic specializations that directly shape its function. The most well-studied aspect of retinal pattern is the 2D topographic map formed by axonal projections of retinal ganglion cells (RGCs), the output neurons of the retina (37). RGC axons project into multiple target locations in the brain in a pattern that maintains the topography of the the visual field (18,37). Another important aspect of retinal patterning is the arrangement of photoreceptors, the light-sensing cells of the retina (31). There are two main types of photoreceptors: rods, which mediate vision in dim light, and cones, which mediate vision in bright light and contribute to high acuity. Cones are further distinguished into subtypes based on the opsin protein that they express. Each opsin is optimized to detect specific wavelengths of light, enabling color vision. Both rod and cone density vary regionally across the retina (12,5).

Among the most striking examples of retinal specialization is the high-acuity area (HAA). In humans, this central HAA, referred to as the fovea, is responsible for high-acuity daylight vision (26). The fovea is characterized by a strikingly different photoreceptor composition compared to the rest of the retina. While ∼95% of photoreceptors outside the fovea are rods, the central portion of the fovea, the foveola, is completely devoid of rods and instead contains an exceptionally dense array of thin, elongated cones (12,31,27). In addition to its specialized photoreceptor mosaic, the fovea contains a uniquely high density of distinct ganglion cells, referred to as “midget” RGCs owing to their small size (14). These specializations minimize convergence of the light signal. At the fovea, one to three cones connect to a single RGC, compared to hundreds in the periphery - enabling exceptional spatial and chromatic acuity (30,47). Although the fovea occupies less than 1% of the retinal area, approximately 50% of the cells in the primary visual cortex receive input that originates from this small region (62, 63). Despite its importance, the formation of the HAA remains poorly understood, as typical mammalian models, e.g. mice and rats, lack a comparable structure. Additionally, foveal development has not been reported in organoid cultures (39,42,10). The fovea is highly susceptible to age-related macular degeneration and developmental abnormalities, such as foveal hypoplasia, making it critical to understand how it develops (20,34).

Similar to humans, certain birds and lizards possess HAAs that enhance sharp daylight vision; some species, such as raptors and brown anoles, even possess two foveae (45,51,48,59). We discovered that chickens have an area which shares many cellular features with the human fovea, including the absence of rods and presence of a high density of cones and RGCs, suggesting that it too enables high-acuity vision (13,5,9). However, unlike the primate fovea, this specialized area of the chicken, has not been directly tested for visual performance. Nonetheless, on the basis of shared anatomical and molecular features, we will refer to this area as the HAA, while noting that the chicken structure may more precisely represent a ‘higher-acuity’ region relative to the peripheral retina. Using the chicken as a model system, we have begun to explore the expression and function of early patterning genes potentially involved in HAA development, focusing on HAA localization within the nasal-central retina and the generation of its characteristic cellular composition.

The developing chicken HAA can be distinctly identified as a nasal-central region with high *Fgf8* expression (9,13). *Fgf8* is expressed most highly within a nasal-central spot which extends temporally into a stripe along the dorso-ventral (DV) border, referred to as the “equator”. Previous studies have demonstrated a correlation between the presence of *Fgf8* expression and the absence of retinoic acid (RA) signaling around the equator (13). Our previous studies indicated that absence of RA is necessary for *Fgf8* expression, and *Fgf8* expression is essential for HAA development (13). A similar expression pattern for RA pathway genes in early human and primate retinas suggests a conserved role across species (13,33). Moreover, we proposed that the retinal progenitor cells (RPCs) in the HAA with high *Fgf8* expression are distinct from other RPCs and possess the properties required to produce the specific cellular composition of the HAA (9,13).

Patterned expression of transcription factors and signaling molecules in the retina have been reported in several species, some showing early patterns that likely endow the eye anlage with coordinates along the naso-temporal (NT) and DV axes. (67,53,49,50,43,56,60,22). Some of these patterns are first established in the neural plate, with some persisting into the developing eye cup, while becoming increasingly refined. These early patterns converge to form asymmetric DV and NT expression domains crucial for establishing and maintaining retinal regional specialization (53,43,60). However, several key gaps remain in understanding the molecular mechanisms of retinal patterning. First, reliance on qualitative imaging limits the ability to rigorously assess subtle overlaps, gradients, and dynamic shifts in gene expression. Second, integration of DV and NT patterning axes with RA/*Fgf8* signaling has not been quantitatively or functionally resolved, leaving unclear whether these pathways might converge to specify the HAA. Third, although large-scale single-cell RNA-sequencing (scRNA-seq) has expanded the understanding of retinal cell states, such datasets are typically analyzed without spatial context, forfeiting topographic patterning information. Finally, evolutionary comparisons of spatial patterning remain sparse, limiting understanding of which molecular strategies are conserved versus species-specific.

In this study, we addressed these gaps by quantifying multiplexed fluorescent in situ hybridization (FISH) signals. These measurements focused on genes with known asymmetric early expression patterns across the DV and NT axes. A custom image analysis pipeline further generated quantitative spatial maps, refining boundaries and overlaps between RA metabolism genes, *Fgf8*, and these other early patterning genes. A large scRNA-seq dataset of chicken RPCs was assembled, from which a framework of DV and NT axes was generated. This approach recapitulated spatial patterns seen by RNA-FISH, identified novel HAA-enriched genes, and revealed patterns of signaling pathway genes. Extending this analysis to human and mouse scRNA-seq data uncovered conserved axis-based programs but divergent fine-scale organization. Genes identified to have expression patterns correlated with the location of the developing HAA in human were further validated by RNA-FISH.

Taken together, this work establishes a generalizable framework for quantitative spatial reconstruction of the retinal patterning landscape, defining positional gene expression domains, enabling cross-species comparisons, and revealing the molecular logic of local specializations such as the retinal HAA. Beyond the retina, it provides a scalable and versatile approach to map spatial organization in developing tissues using scRNA-seq. As well, by leveraging the large and growing body of scRNA-seq datasets across a wide diversity of species, it provides an inexpensive avenue to evolutionary comparisons of patterning.

## RESULTS

### Quantitative mapping of expression domains of *Fgf8* and RA pathway genes

Given the significance of *Fgf8* and RA signaling in HAA development, we aimed to map gene expression patterns that might establish or depend upon these restricted domains of expression. While single and double ISHs detect expression of one or two genes at a time, limiting analysis to a small number of patterns, multiplexed RNA-FISH allows simultaneous detection of expression of multiple genes, offering a more comprehensive view for pattern comparisons (8). To quantify subtle expression patterns, we developed a MATLAB-based workflow for retinal flat mounts (see Methods), converting raw fluorescence images into standardized, spatially registered intensity maps [**Figure S1**]. Alignment across samples enabled comparison of profiles, integration of multiplexed datasets, and visualization of multiple gene patterns within a unified spatial framework. These maps allowed for a rigorous assessment of domain overlaps, asymmetry, and boundary definition, providing a robust tool for dissecting retinal spatial organization [**Figure 1,Figure S2**].

**Figure 1.**
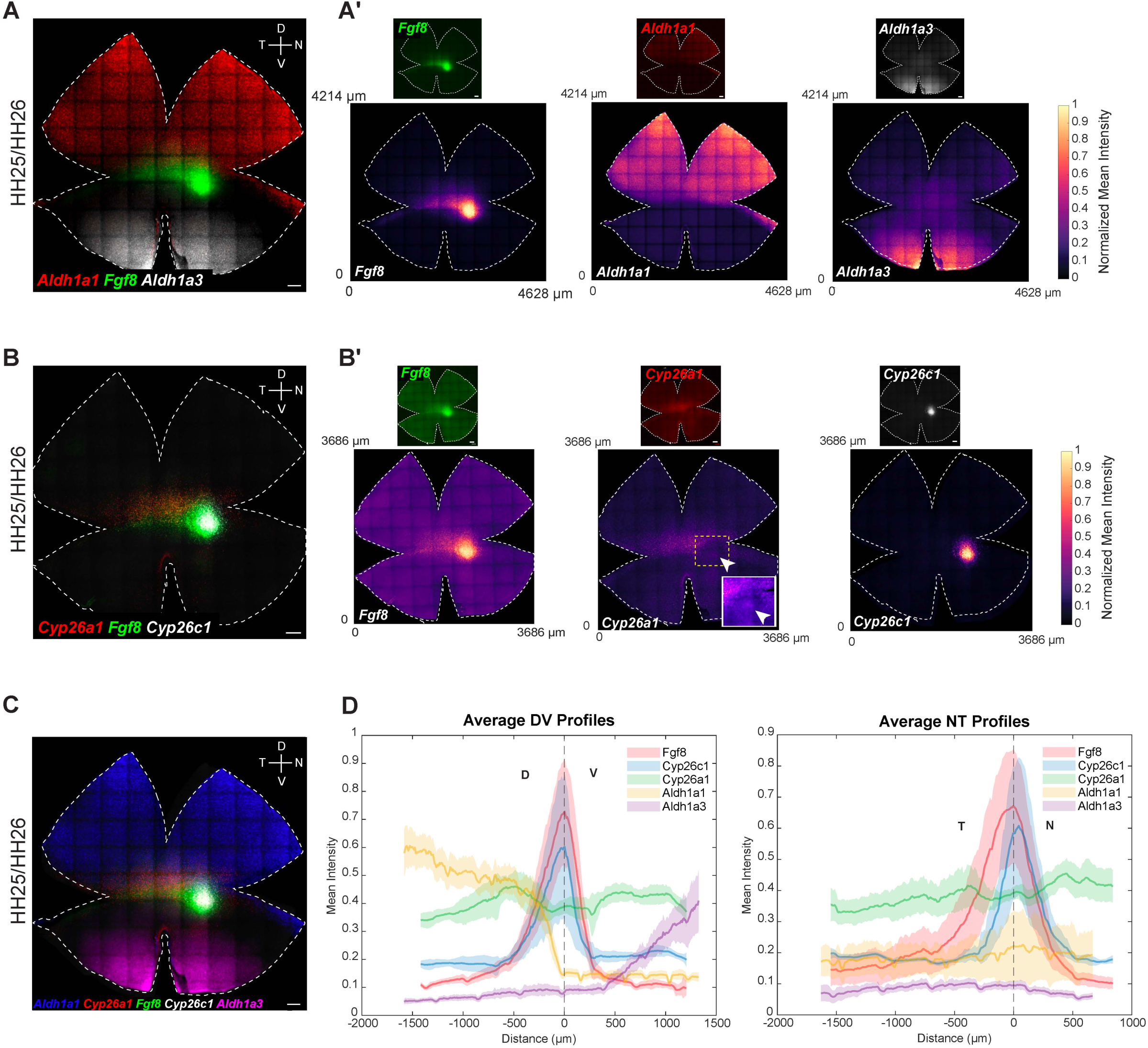
Quantitative mapping of expression domains of Fgf8 and RA pathway genes. **(A-C)** Two sequential rounds of multiplexed RNA-FISH were performed on a single HH25/HH26 retinal whole mount. (A) First round: probes for *Fgf8*, *Aldh1a1*, and *Aldh1a3*. (B) Second round: probes for *Fgf8*, *Cyp26a1*, and *Cyp26c1*. (C) Overlay of both rounds. (A’, B’) Individual channel images with corresponding quantified intensity heatmaps generated using the MATLAB pipeline. Inset in (B’) highlights the transient “bull’s-eye” spot of *Cyp26a1* expression, as marked by the white arrowhead. **(D)** Spatial expression profiles of *Fgf8* and RA pathway genes across the developing retina. (Left) Average DV expression intensity profiles of *Fgf8*, *Cyp26c1*, *Cyp26a1*, *Aldh1A1*, and *Aldh1A3*. (Right) Average NT profiles for the same genes. Dark lines indicate mean normalized fluorescence intensity, and shaded regions represent the standard error of the mean (SEM). Profiles were calculated from N = 3 biological replicates. Scale bars, 200 μm. HH, Hamburger and Hamilton; D, Dorsal; V, Ventral; N, Nasal; T, Temporal. All patterns were observed in N ≥ 3 retinas, apart from the *Cyp26a1* bull’s-eye pattern, which is transient and has been previously described (13).

To benchmark this quantification method, and to precisely map localized gene expression patterns, the expression patterns of *Fgf8* and RA pathway genes were examined in detail through multiplexed RNA-FISH on retinal flat mounts from Hamburger-Hamilton (HH) stage 25/26 embryos (23). At this stage, neurogenesis is active in the central retina and the retina is large enough for flat mount RNA-FISH (46,3,9). The formation of the ganglion cell layer (GCL) is also first apparent at this time. Precise expression patterns were observed for *Fgf8* and genes encoding RA-synthesizing (*Aldh1a1* and *Aldh1a3)* and RA-degrading (*Cyp26a1* and *Cyp26c1)* enzymes, consistent with our previous reports (54,13,9). *Aldh1a1* was expressed in a dorsal domain extending to the equator, whereas *Aldh1a3* was restricted to the region ventral to the equator [**Figure 1A, A’, D**]. While *Aldh1a3* did not share a boundary with *Fgf8*, the ventral border of *Aldh1a1* abutted the dorsal edge of the equatorial *Fgf8* stripe on the retinal flat mount [**Figure 1A, A’, D**]. In a second round of multiplexed RNA-FISH, expression patterns of RA-degrading enzymes, *Cyp26a1* and *Cyp26c1*, were assayed relative to the *Fgf8* pattern on the same tissue. Consistent with earlier observations (13), *Cyp26a1* was expressed in a dorsal-central domain. Its expression partially overlapped with *Fgf8* along the equatorial stripe, while being notably absent from the nasal *Fgf8* spot, except for a transient bull’s-eye pattern [**Figure 1B, B’, D**]. *Cyp26c1* expression was highest in a small spot partially overlapping the nasal *Fgf8* spot [**Figure 1B, B’, D**]. Merging the signals for all these genes from the same tissue further offered an enhanced view of the positioning of ventral borders of *Aldh1a1* and *Cyp26a1* expression and the dorsal boundary of *Fgf8* expression on the same retinal flat mount [**Figure 1C**]. In these experiments, two sequential rounds of RNA-FISH were required, and tissue had to be carefully realigned between rounds – a process that was time-consuming and imposed practical limitations. Our MATLAB-based quantitative workflow helped circumvent this process and improve upon the assessment of pattern reproducibility across samples. It produced results similar to those assessed by eye on flat mounts. This analysis further allowed for integration of multiple expression patterns across samples, thus serving as a reliable alternative to multi-round RNA-FISH on the same tissue. [**Figure 1D; Figure S1**, **Figure S2**].

DV cross-sections spanning the *Fgf8* spot showed similar overlapping domains for RA-synthesizing and RA-degrading enzymes, with signal observed in the outer neuroblastic layer of the retina, where progenitor cells reside [**Figure S3**] (13,9). Consistent with the flat mount data, *Fgf8* expression exhibited a sharp ventral border, contrasting with its diffuse dorsal border. Notably, a thicker area of the section, indicating the location of the developing HAA (9), was evident in the region of the cross-section characterized by the presence of *Fgf8* and *Cyp26c1* expression and complete absence of *Aldh1a1* and *Aldh1a3*.

### Expression patterns of early DV patterning genes relative to *Fgf8*

The neural retina is divided into multiple domains of gene expression along the DV axis, established early in development (22,5,3,49,43,44,56). To explore the relationship between the expression domains of DV-patterned genes and *Fgf8*, whole mount multiplexed RNA-FISH was performed. Early dorsal markers include *Tbx2, Tbx3,* and *Tbx5*, members of the *Tbx2* subfamily of T-box transcription factor genes (22). Early ventral markers include *cVax* (chicken ortholog of human *Vax1)*, a ventralizing homeobox transcription factor, and *Ventroptin*, an antagonist of BMP signaling, presumably *Bmp2/4,* which are expressed in the early dorsal retina (53,49,50). Expression patterns were assayed and quantified in retinas from two early developmental stages – HH25/26 and HH28/29 (8,23) [**Figure 2, Figure S4, Figure S5, Figure S6**].

**Figure 2.**
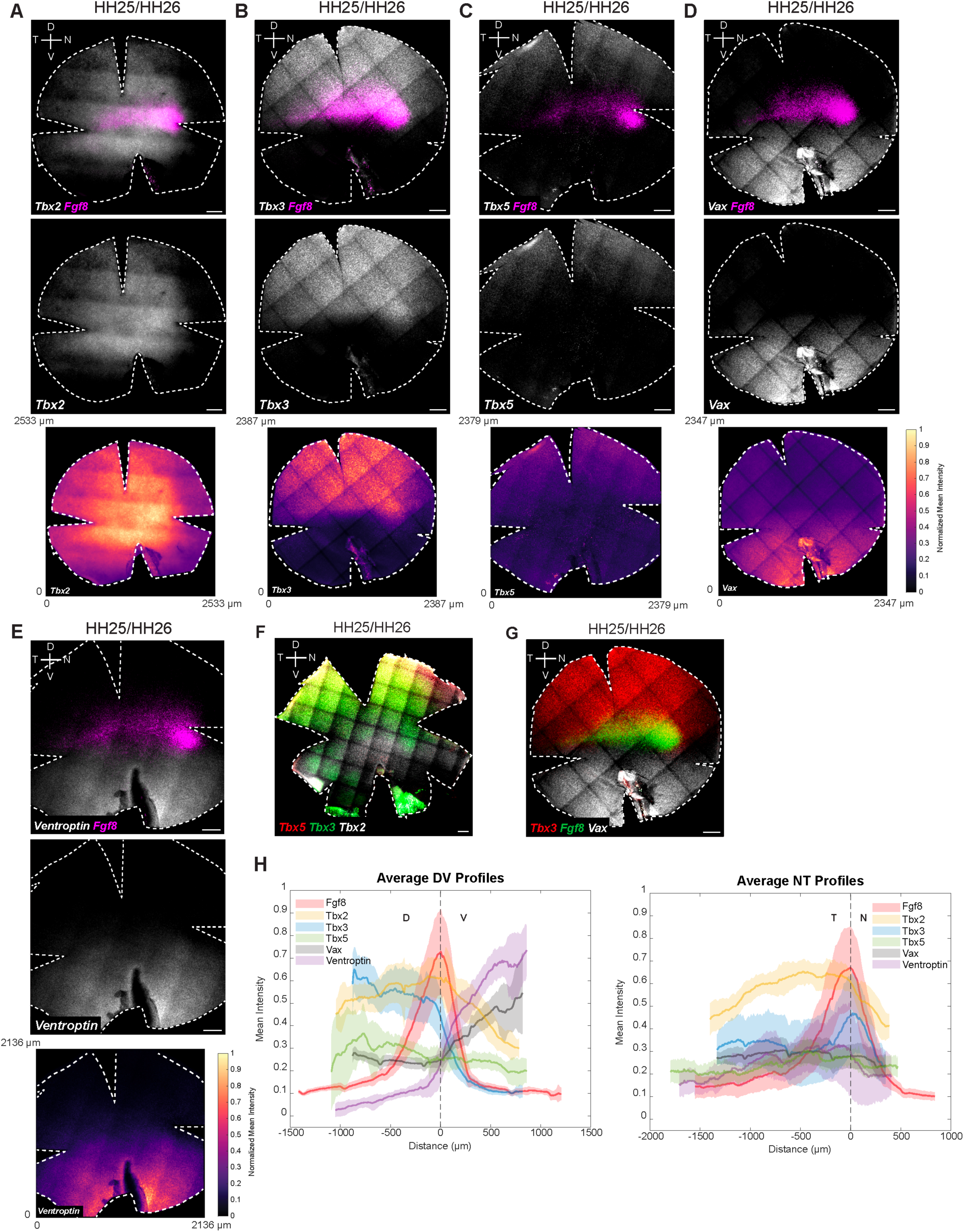
Quantitative mapping of expression domains of early DV genes relative to *Fgf8*. Multiplexed RNA-FISH was performed on HH25/HH26 **(A-G)** retinal whole mounts. (A) *Fgf8* and *Tbx2*; (B) *Fgf8* and *Tbx3*; (C) *Fgf8* and *Tbx5*; (D) *Fgf8* and *Vax*; (E) *Fgf8* and *Ventroptin*. Each panel shows merged images, single-channel expression images, and corresponding quantified intensity heatmaps. Additional combinations for multiplexed RNA-FISH included (F) *Tbx2, Tbx3*, and *Tbx5*; (G) *Fgf8*, *Tbx3*, and *Vax*. **(H)** Spatial expression profiles of early DV patterning genes across the developing retina. (Left) Average DV expression intensity profiles of *Fgf8*, dorsally patterned genes *(Tbx2, Tbx3, and Tbx5)*, and ventrally patterned genes *(Vax and Ventroptin)*. (Right) Average NT profiles for the same genes. Dark lines indicate mean normalized fluorescence intensity, and shaded regions represent the standard error of the mean (SEM). Profiles were calculated from N = 3 biological replicates for *Tbx2, Tbx3, Vax, Fgf8* and from N = 2 biological replicates for *Tbx5, Ventroptin*. Scale bars, 200 μm. HH, Hamburger and Hamilton; D, Dorsal; V, Ventral; N, Nasal; T, Temporal. All patterns were observed in N ≥ 3 retinas. The *Tbx2* image shown here (A) is a widefield image, whereas all other panels are maximum projections of confocal z-stacks.

*Tbx* genes were expressed in nested dorsal domains, as was previously reported (22, 70) [**Figure 2**]. At both stages, *Tbx2* extended further ventrally than *Fgf8*, while the *Tbx3* ventral border aligned with *Fgf8*. Notably, a domain of higher *Tbx3* expression was observed overlapping the *Fgf8* spot [**Figure 2B and H, Figure S4**]. *Tbx5* was confined to the dorsal-most region, with its ventral border just dorsal to the *Fgf8* stripe. Quantification of the RNA-FISH signal further revealed that *Tbx2*, *Tbx3*, and *Tbx5* were largely absent from the far peripheral retina [**Figure 2H, Figure S4**]. *Tbx2* displayed the most pronounced center-to-periphery expression gradient, with peak *Tbx2* expression observed dorsal-temporal to the *Fgf8* spot [**Figure 2A, Figure S4**]. Ventrally, *cVax* precisely abutted the *Fgf8* stripe and spot [**Figure 2D and H, Figure S4**]. When co-visualized, the dorsal border of *cVax* aligned with the ventral border of *Tbx3* and *Fgf8*. *Ventroptin* exhibited an oblique pattern, with expression largely restricted to the ventral retina. Similar to *cVax,* its dorsal border abutted the ventral edge of *Fgf8*. Interestingly, for both *cVax* and *Ventroptin*, we observed a sharp boundary ventral to the *Fgf8* spot [**Figure 2D and H, Figure S4**].

Ephrins and related genes also have been reported to show domain-specific expression patterns in the developing retina. They play key roles in guiding RGC axons to establish retinotopic maps (38). Consistent with previous reports, we observed that ligands *EphrinB1* and *EphrinB2* (also known as *EfnB1* and *EfnB2*) are dorsally restricted, with the ventral domain of *EphrinB1* more dorsal than that of *Ephrin B2* (43). The receptor, *EphB2*, is confined ventrally, consistent with previous reports. Interestingly, *EphrinB1* showed a *Tbx3*-like pattern, with potential enrichment within the *Fgf8* spot [**Figure S7**]. Together, these results contribute to a spatial map demarcating gene expression domains along the DV axis and highlight enrichments, exclusions, and shared borders associated with the developing HAA.

### Expression patterns of early NT patterning genes relative to *Fgf8*

The NT axis of the retina is also divided into distinct domains of gene expression early in development (67,56). The winged-helix transcription factors, *FoxG1* and *FoxD1,* display complementary regional expression, with *FoxG1* expressed nasally and *FoxD1* temporally (50,60). *FoxG1* has been shown to confer nasal identity, while *FoxD1* confers temporal identity in the developing chicken retina (50,60). *SOHo1*, a homeodomain transcription factor, has also been shown to be expressed nasally (15). Expression of these genes was examined in HH25/26 and HH28/29 retina alongside *Fgf8* to determine whether their expression domains correspond to the *Fgf8* spot [**Figure 3, Figure S8, Figure S9, Figure S10**].

**Figure 3.**
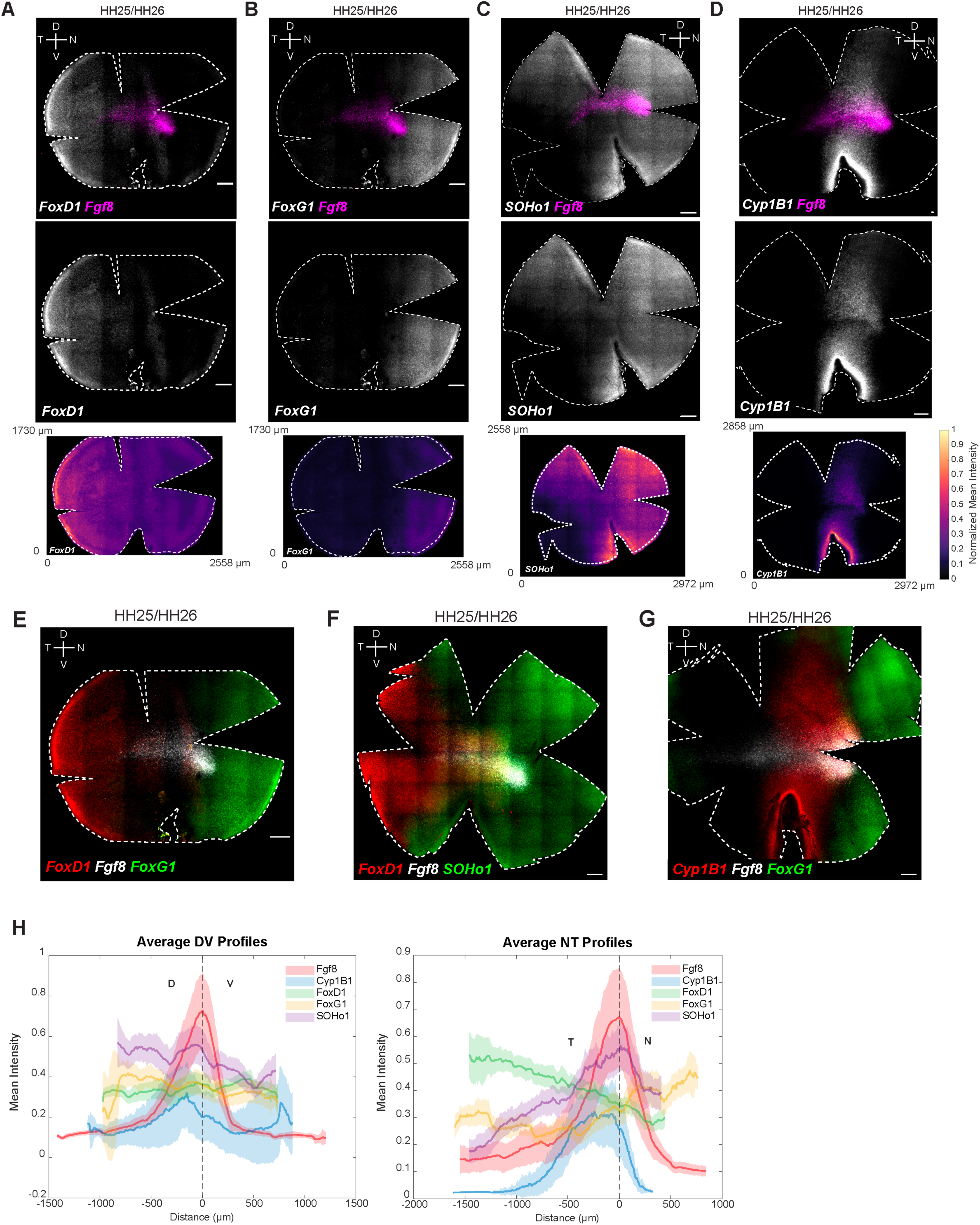
Quantitative mapping of expression domains of early NT genes relative to *Fgf8*. Multiplexed RNA-FISH was performed on HH25/HH26 **(A-G)** retinal whole mounts. (A) *Fgf8* and *FoxD1*; (B) *Fgf8* and *FoxG1* (C) *Fgf8* and *SOHo1*; (D) *Fgf8* and *Cyp1B1*. Each panel shows merged images, single-channel expression images, and corresponding quantified intensity heatmaps. Additional combinations for multiplexed RNA-FISH included (E) *Fgf8, FoxD1, and FoxG1;* (F) *Fgf8, FoxD1, and SOHo1;* (G) *Fgf8, Cyp1B1, and FoxG1.* **(H)** Spatial expression profiles of early NT genes across the developing retina. (Left) Average DV expression intensity profiles of *Fgf8*, nasally patterned genes (*FoxG1,* and *SOHo1*) and temporally patterned genes (*FoxD1)*. (Right) Average NT profiles for the same genes. Dark lines indicate mean normalized fluorescence intensity, and shaded regions represent the standard error of the mean (SEM). Profiles were calculated from N = 3 biological replicates. Scale bars, 200 μm. HH, Hamburger and Hamilton; D, Dorsal; V, Ventral; N, Nasal; T, Temporal. All patterns were observed in N ≥ 3 retinas.

*FoxD1* expression was confined to the temporal-most retina at both stages [**Figure 3A and H**], whereas *FoxG1* was localized nasally, with its temporal edge aligning with the nasal border of the *Fgf8* spot [**Figure 3B and H**]. This expression pattern is consistent with previous reports suggesting that *FoxG1* and *FoxD1* specify nasal and temporal retinal identities, respectively, and are mutually repressive (50,60). Within the nasal domain, *FoxG1* showed higher ventral expression, indicating additional DV asymmetry [**Figure S8**]. *SOHo1* expression extended from the nasal region, filling in the gap between the expression areas of *FoxG1* and *FoxD1* [**Figure 3C and H**], with an enrichment around the *Fgf8* spot [**Figure S8**]. Additionally, we discovered an unusual NT expression pattern of *Cyp1B1*, a gene encoding a protein capable of RA synthesis (7,64). In contrast to the DV expression patterns of other RA synthesizing enzymes (*Aldh1a1* and *Aldh1a3), Cyp1B1* exhibited a distinct NT-restricted pattern, forming an oblique stripe with strong signal near the optic fissure [**Figure 3D and H**]. Strikingly, there was a stripe in the equatorial area adjacent to the *Fgf8* spot where there was no detectable *Cyp1B1* expression, displaying a central notch immediately temporal to it [**Figure S8**]. Together, these data delineate molecular territories along the NT axis, highlighting distinct boundaries and localized enrichments at the *Fgf8* spot.

### Expression patterns of genes with profiles similar or complementary to *Fgf8*

To further elucidate the factors contributing to HAA development, we surveyed the literature to identify genes with expression patterns similar to *Fgf8*. *Bmp2* has been reported to be distinctly expressed along the equator, with higher expression in the nasal-central region (4). We observed a similar pattern for *Bmp2*, and co-hybridizing for *Fgf8* revealed nasally enriched expression of *Bmp2* in a small spot nested within the *Fgf8* spot. Additionally, the ventral border of *Bmp2* expression aligned perfectly with the ventral border of *Fgf8* expression [**Figure 4A, Figure S11A and A’**].

**Figure 4.**
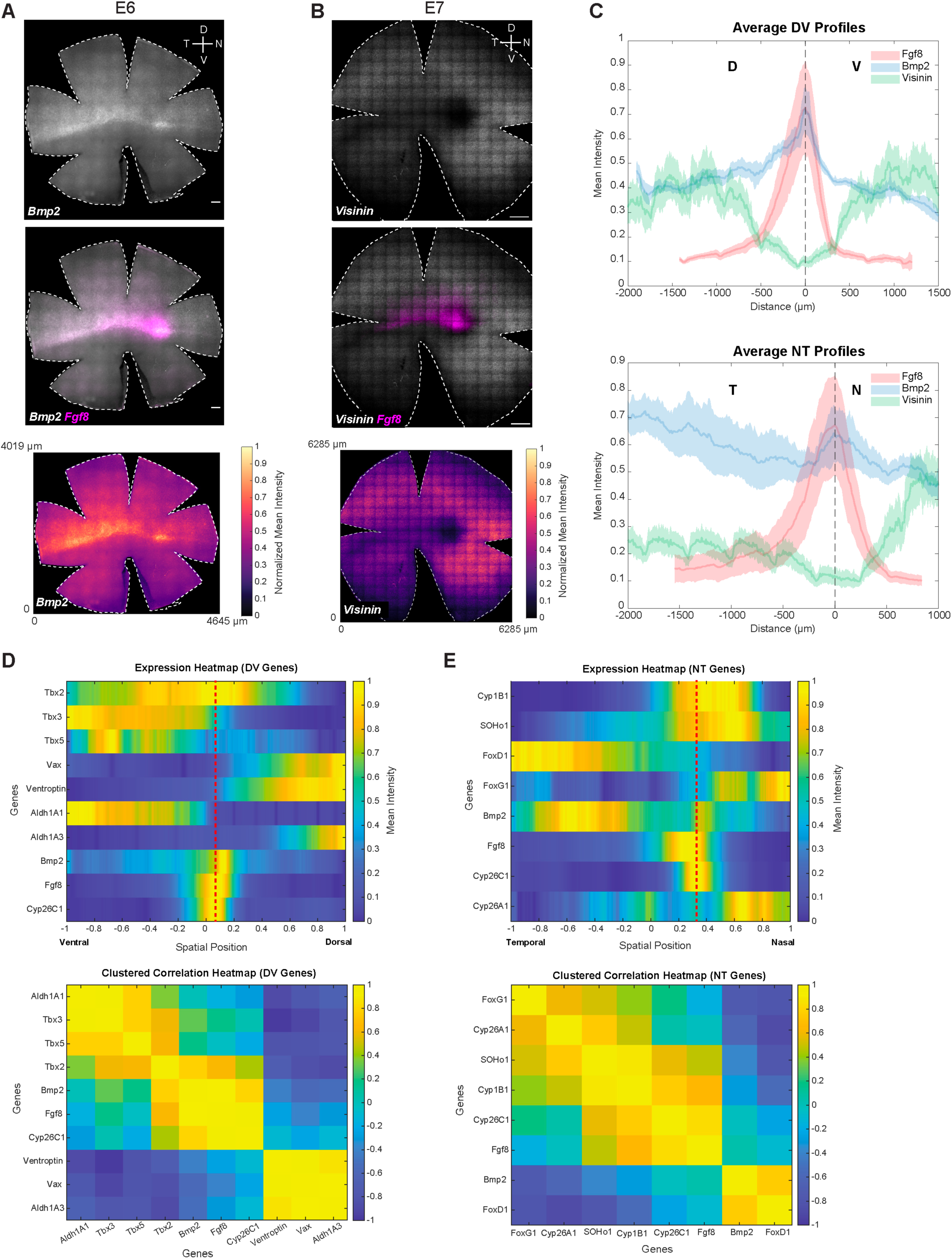
Quantitative mapping of expression domains of early developmental genes relative to *Fgf8*. **(A-B)** Multiplexed RNA-FISH was performed on (A) E6 retinal whole mounts for *Fgf8* and *Bmp2*; and (B) E7 retinal whole mounts for *Fgf8* and *Visinin*. Each panel shows merged images, single-channel expression images, and corresponding quantified intensity heatmaps. **(C)** Spatial expression profiles of *Fgf8*, *Bmp2* and *Visinin* across the developing retina. (Top) Average DV expression intensity profiles of *Fgf8*, *Bmp2 and Visinin*. (Bottom) Average NT profiles for the same genes. Dark lines indicate mean normalized fluorescence intensity, and shaded regions represent the standard error of the mean (SEM) for *Fgf8* and standard deviation (SD) for *Bmp2* and *Visinin*. Profiles were calculated from N = 3 biological replicates for *Fgf8*, and N =1 for *Bmp2* and *Visinin*. Scale bars, 200 μm. Bmp2 pattern was observed in N ≥ 3 retinas. *Visinin*-free spot was observed transiently in N = 1 retina and has been previously described (5,55). **(D, E)** Spatial correlation analysis of selected genes from RNA-FISH data along the (D) DV and (E) NT axis. Top, heatmap showing normalized expression of selected genes along the DV and NT axis. Right, clustered correlation heatmap for the same genes based on pairwise Pearson correlation of their spatial expression profiles, revealing (D) dorsal, central, ventral and (E) nasal, central, temporal gene modules. Genes are ordered according to correlation-based hierarchical clustering. The red dashed line indicates the approximate position of the developing HAA, marked at the center of the *Fgf8* and *Cyp26C1* expression domains. E, Embryonic day; D, Dorsal; V, Ventral; N, Nasal; T, Temporal.

In addition to patterns specifically localized to the developing HAA, the expression patterns that were selectively excluded from the HAA were also assessed. *Visinin*, which encodes a calcium-binding protein and is an early photoreceptor marker (25,65), has been reported to transiently lack expression specifically in the nasal-central retina (5,55). The transient *Visinin*-free region has been proposed to mark the future rod-free and rod-sparse retinal regions. Consistent with these reports, a distinct *Visinin*-free spot in the nasal-central retina and a central *Visinin-free* stripe extending temporally was observed [**Figure 4A, Figure S11B**]. Co-staining with *Fgf8* demonstrated complementary patterns, with *Visinin* excluded from the *Fgf8* spot and stripe - territories that later give rise to rod-free and rod-sparse domains (9). In addition to this nasal-central pattern, *Visinin* expression exhibited further asymmetry along the DV and NT axes. In the nasal retina, *Visinin* expression was higher ventrally than dorsally, whereas in the temporal retina it was enriched dorsally compared to ventrally [**Figure S11B and B’**]. The expression patterns of *Bmp2* and *Visinin* highlight both enriched and excluded molecular signatures in the developing HAA, providing further resolution of its distinct positional identity within the early chicken retina.

### Spatial correlation analysis of quantitative RNA-FISH data

Given the availability of quantitative RNA-FISH data with defined spatial coordinates, we performed spatial correlation analysis to systematically quantify relationships between gene expression patterns along the DV and NT axes. This approach enabled the identification of spatially co-expressed gene modules based on similarity of their position-dependent expression profiles [**Figure 4D and E**]. Analysis of RNA-FISH expression profiles along the DV axis revealed three spatially ordered gene modules **[Figure 4D]**. *Tbx2, Tbx3, Tbx5*, and *Aldh1A1* formed a dorsal domain with peak signal at the dorsal end and a graded decline toward more ventral positions. *Fgf8, Bmp2*, and *Cyp26C1* were concentrated in a narrow central band at the HAA, whereas *Vax1, Ventroptin*, and *Aldh1A3* marked the ventral compartment. Pairwise correlation-based clustering along the spatial axis mirrored this organization: dorsal and ventral genes each formed internally coherent clusters that were strongly anticorrelated with one another, while the *Fgf8*, *Bmp2,* and *Cyp26C1* occupied an intermediate central cluster. Dorsal markers showed stronger positive correlations with this central module than did ventral markers, indicating asymmetric coupling between dorsal and central domains. The same analytical framework along the NT axis revealed an analogous three-way organization **[Figure 4E]**. *FoxD1* and *Bmp2* were biased toward temporal positions, *FoxG1* and *Cyp26A1* toward the nasal end, and *Fgf8* and *Cyp26C1* were enriched centrally at the HAA. In addition, *SOHo-1* and *Cyp1B1* showed intermediate profiles, with expression shared between central and nasal regions. Clustering divided these genes into temporal (*FoxD1/Bmp2*), central (*Fgf8/Cyp26C1*), and nasal (*FoxG1/Cyp26A1*) modules, with strong positive correlations within each group and clear anticorrelation between temporal and nasal programs, emphasizing the transcriptional distinctness of the central signaling hub that separates opposing NT identities.

### Reconstruction of 2D topographic maps from chicken scRNA-seq datasets

The RNA-FISH results revealed the relative patterning of several genes with spatially restricted expression patterns. We next considered how this information might be leveraged in combination with scRNA-seq to investigate a broader set of gene expression patterns. Although spatial information is typically lost in scRNA-seq, the relative expression of known spatially restricted genes might enable inference of each cell’s approximate position [**Figure 5A**]. To this end, a dataset comprising in-house and public scRNASeq data from early chicken retina was assembled (E5–E6) [**Figure S12**].

**Figure 5.**
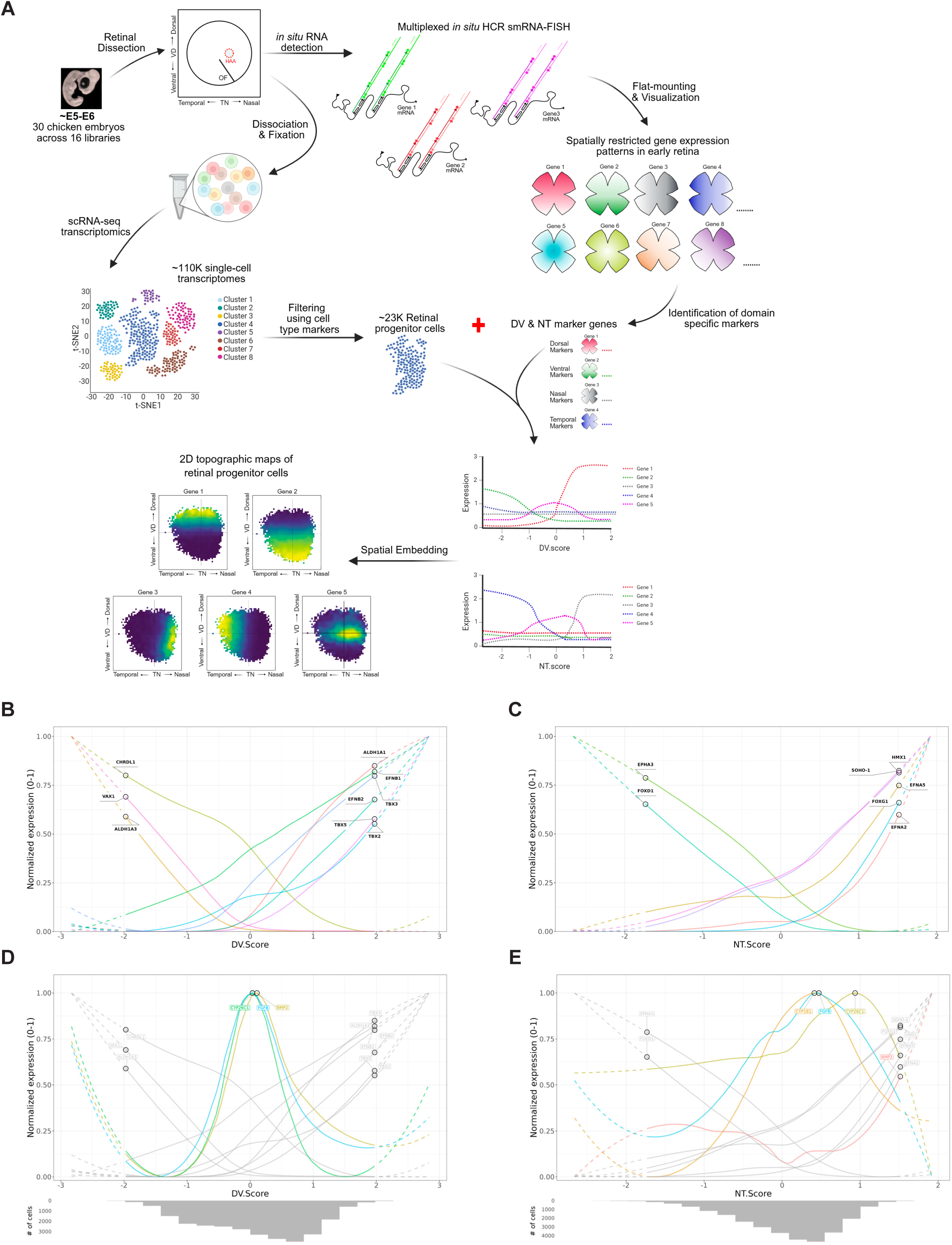
Generation of DV and NT scores using single-cell transcriptomes from the developing chicken retina. **(A)** Schematic workflow for reconstruction of spatial 2D topographic maps from chicken scRNA-seq datasets. **(B)** Spatial expression pattern of marker genes used to calculate the DV.Score. From the scRNA-seq values, D, V scores were computed using the relative expression of the following genes as a scoring set: {*Tbx5, Tbx2, Tbx3, Aldh1a1, EfnB2, EfnB1*} for D, {*Vax1, Chrdl1, Aldh1a3*} for V. **(C)** Spatial expression pattern of marker genes used to calculate the NT.score. (D) From the scRNA-seq values, N, T scores were computed using the relative expression of the following genes as a scoring set: {*FoxG1, SOHo-1, Hmx1, EfnA5, EfnA2*} for N, and {*FoxD1, EphA3*} for T. **(D)** Test of the spatial reconstruction pipeline using independent genes with well-defined axial expression patterns that were not included in the D, V scoring set; *Fgf8, Cyp26c1 and Bmp2* for a test of the mapping along the DV axis. **(E)** Test of the spatial reconstruction pipeline using independent genes with well-defined axial expression patterns that were not included in the N, T scoring set; *Fgf8, Cyp1B1, and Cyp26c1* for a test of the mapping along the NT axis. The histograms on the bottom represent coverage by aggregating all cells that contributed to the bins (total bins=50). The dashed lines represent the NT axis bins that do not have more than 50 cells covered, suggesting less confidence for proper DV/NT.Score. HAA, High(er) Acuity Area; OF, Optic Fissure; E, Embryonic day; D, Dorsal; V, Ventral; N, Nasal; T, Temporal; DV.score, Dorsal-Ventral score; NT.score, Nasal-Temporal score. Panel (A) Created in BioRender; Joisher, H. (2025) https://BioRender.com/vhj5dus

Starting from an initial ∼110K single-cell transcriptomes, a total of 23,706 RPCs from 30 embryos across 16 libraries were identified [**Figure S12**]. DV and NT position scores (DV.Score and NT.Score) were assigned to each cell based on the relative expression of 9 DV and 7 NT biased genes, respectively (see Methods) [**Figure S13, Figure S14**]. The validity of this scoring method was tested through analysis of the 2D topographic gene expression pattern of genes with well-defined axial expression patterns, genes that were not among the 16 genes used by the scoring algorithm – in this case *Fgf8* and *Cyp1B1*. The predicted 2D topographical expression pattern of these test genes closely recapitulated the nasal-central enrichment of *Fgf8* and the oblique stripe of *Cyp1B1* observed by RNA-FISH. These data demonstrated that the limited marker set was sufficient to capture nuanced spatial patterns in the developing retina [**Figure 5B-E**].

This analysis was then extended to the genes mapped in physical space in Figures 1-4. **[Figure S13E, Figure S14E]**. Spatial heatmaps of a subset of DV genes, ordered by inferred DV position revealed a dorsal *TBX/Aldh1A1* module (*Tbx2, Tbx3, Tbx5, Aldh1A1*), a central *Fgf8/Bmp2/Cyp26C1* module, and a ventral *Vax1/Ventroptin/Aldh1A3* module. Clustered correlation heatmaps mirrored these patterns, revealing three internally coherent clusters: dorsal genes were strongly positively correlated with one another, central genes were likewise positively correlated, and ventral markers formed a third, tightly correlated group, with dorsal and ventral compartments anticorrelated **[Figure S13E]**. Consistent with the RNA-FISH results, the central genes correlated more strongly with the dorsal than with the ventral module **[Figure 4D]**. Applying the same analysis along the NT axis also revealed a tripartite organization, with module separation that, while slightly subtler than along the DV axis, was still readily discernible **[Figure S14E]**. Reconstructed NT expression profiles showed *Bmp2* and *FoxD1* enriched toward temporal positions, *FoxG1* and *SOHo-1* biased nasally, and *Fgf8* and *Cyp26C1* peaking centrally, with *Cyp26A1* and *Cyp1B1* spanning central–nasal regions. Correlation-based clustering grouped these genes into temporal (*FoxD1/Bmp2*), central (*Fgf8/Cyp26C1/Cyp26A1/Cyp1B1*), and nasal (*FoxG1/SOHo-1*) modules, with strong within cluster correlations and pronounced anticorrelation between temporal and nasal modules.

By generating inferred spatial coordinates using the DV and NT scores, the single-cell transcriptomes were placed in two-dimensional spatial bins to visualize the 2D topographical expression of genes [**Figure 6**]. These reconstructed 2D topographic maps closely recapitulated spatial patterns as observed in the RNA-FISH experiments. Each transcriptome could also be traced back to its embryo of origin, enabling analysis of embryo-to-embryo variability when comparing different retinal regions. This further allowed differential gene expression analysis to identify spatially concordant and discordant genes for specific regions in the developing retina [**Figure 6H**]. For example, using the *Cyp26c1* expression profile to define the developing HAA, additional genes enriched in this region were identified. To systematically assess retinal spatial organization, up to twenty genes with the strongest spatial autocorrelation were identified (“anchor” genes). These “anchor” genes were then used to rank other genes by spatial correlation, and clusters were defined where expression patterns were mutually correlated. In total, this approach identified 17 clusters with distinct spatial expression patterns, including the well-characterized axis-biased patterns (DV/NT) [**Figure S16A**]. Notably, some established spatial markers subdivided into separate clusters, reflecting subtle differences in their expression domains (e.g., *EphB2, Aldh1a3*), while other genes associated with early differentiation (e.g., *Onecut1*) showed localized enrichment likely corresponding to the center-to-periphery differentiation axis.

**Figure 6.**
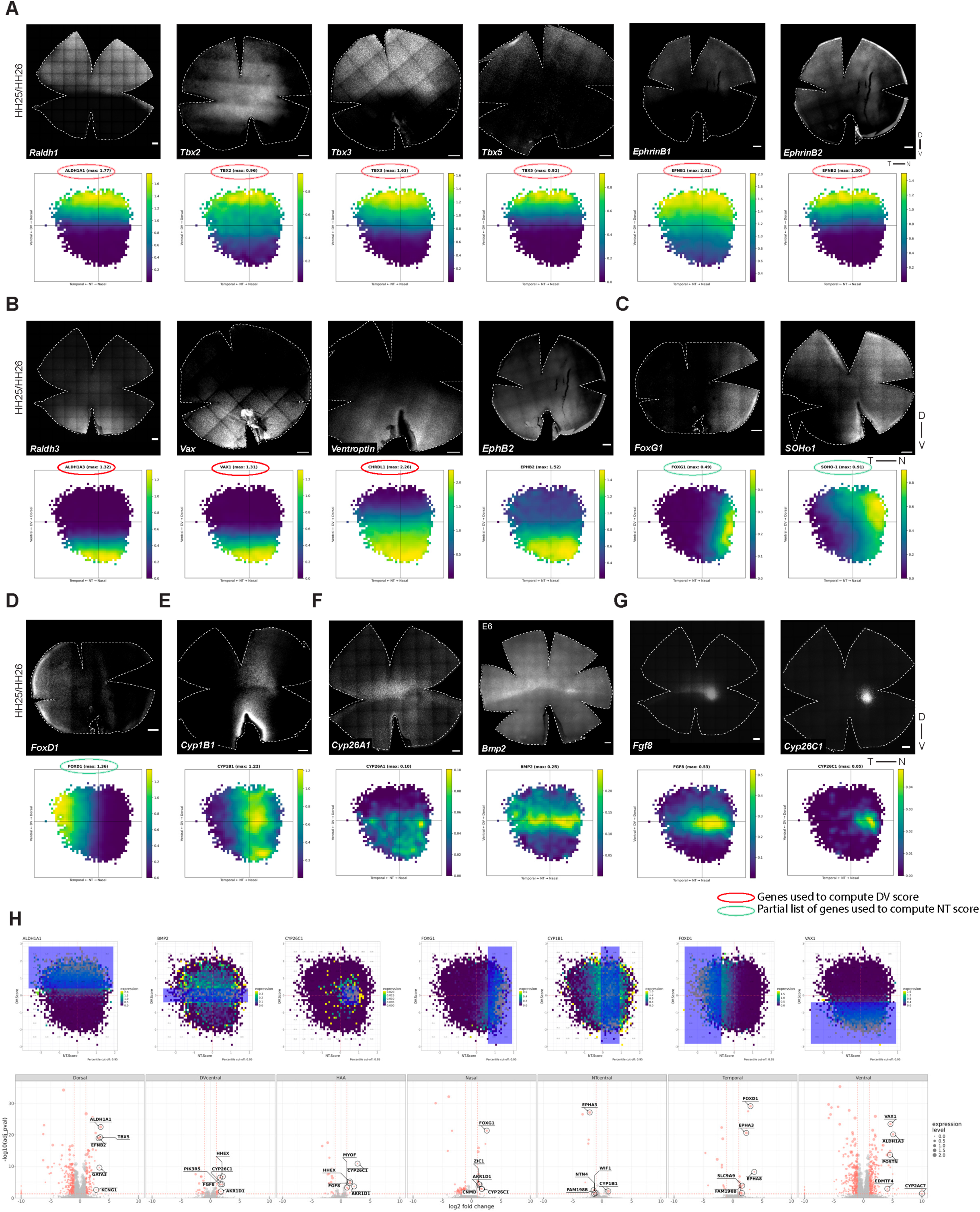
Reconstruction of 2D topographic maps from chicken scRNA-seq datasets. **(A-G)** 2D topographic maps of retinal gene expression compared with RNA-FISH patterns. Each panel shows the 2D topographic expression map reconstructed from scRNA-seq alongside the corresponding multiplexed RNA-FISH image for the same gene. (A) Dorsally restricted genes: *Aldh1a1, Tbx2, Tbx3, Tbx5, EphrinB1 (EFNB1), and EphrinB2 (EFNB2)* (B) Ventrally restricted genes: *Aldh1a3, Vax, Ventroptin (CHRDL1), and EphB2* (C) Nasally restricted genes: *FoxG1, and SOHo1* (D) Temporally restricted genes: *FoxD1* (E) Centrally restricted genes along the NT axis: *Cyp1b1* (F) Centrally restricted genes along the DV axis: *Cyp26a1* and *Bmp2* (G) HAA-specific genes: *Fgf8* and *Cyp26c1.* Red circle marks the genes used to calculate the DV score; green circle marks the partial list of genes used to calculate the NT score **(H)** Visual representation of virtual selection of the specific regions in the 2D topographic maps to assess differentially enriched or de-enriched genes across these regions. The 2D topographic gene expression of “region-specific marker gene” was overlayed with area bins, and the selected region was overlayed with blue. Volcano plots were generated using pseudobulked gene expression samples that are enriched or de-enriched in the selected regions. Points are scaled by average gene expression level, and genes above 2-fold expression difference between in-group and out-group and adjusted p-values <0.05 are shown in red. The top 5 enriched genes without the “LOC” prefix are highlighted using enlarged, black-outlined points accompanied by gene labels. See also Supplemental Table 1. Region-specific marker genes: (D) *Aldh1a1*, (Central domain across DV axis) *Bmp2*, (HAA) *Cyp26c1,* (N) *FoxG1*, (Central domain across NT axis) *Cyp1B1*, (T) *FoxD1,* (V) *Vax1.* Scale bars, 200 μm. “Max” refers to the gene expression value used to normalize the upper limit of the viridis color scale. Rather than using the absolute maximum, which can be skewed by a few outlier bins, normalization is based on a specified percentile cutoff (p). All values above this threshold are displayed using the brightest color, ensuring that extreme expression levels do not compress the dynamic range of the visualization and that spatial differences in the majority of the data remain visible. HAA, High(er) Acuity Area; HH, Hamburger and Hamilton; D, Dorsal; V, Ventral; N, Nasal; T, Temporal; DV.score, Dorsal-Ventral score; NT.score, Nasal-Temporal score.

Having established a robust framework using anchor-based clustering, the analysis was extended to the identification of additional genes with spatially restricted expression patterns [**Figure S16B-D**]. The analysis revealed novel candidates with strong enrichment in the *Fgf8* spot, including *Npy, Myof, Hhex,* and *Alr1d1*. These genes consistently ranked among the top correlations for more than one of the reference genes (*Fgf8, Cyp26c1* and *Bmp2*), which are shown to be enriched in the developing HAA [**Figure 1**, **Figure 4**]. Although their specific roles in retinal development remain unclear, their restricted expression suggests contributions to specific developmental events in this region. Along the NT axis, top-correlation queries for *Cyp1B1* did not reveal any similarly patterned genes, possibly due to the relatively less organized structure of NT gene domains at this stage [**Figure S17D**].

Given the critical role of *Fgf8* in HAA development, the spatial expression profiles of key components of the *Fgf* signaling pathway were examined [**Figure S17A and B**]. Several ligands and receptors showed dorsal (*Fgf7, Fgf12, Fgf14)* or ventral (*FgfR1, FgfRL1)* enrichment, but none exhibited the precise nasal-central localization characteristic of *Fgf8*. In contrast, some downstream effectors of *Fgf* signaling, including *Dusp6, Spry1*, and *Spry2* were enriched within the *Fgf8* spot. In addition to RA and *Fgf8* signaling, *Bmp* signaling also appeared to be regionally specialized in the developing retina (41,2). Among BMP pathway components, *Bmp2* showed the strongest and most restricted enrichment within the *Fgf8* spot, consistent with RNA-FISH data, suggesting a role in HAA specification. Other ligands and downstream factors displayed complementary patterning: *Bmp7*, *Tbx5*, *ID2/3*, and *Bamb*i were dorsal, whereas *Chrdl1* or *Ventroptin*, *Bmpr1A/B*, *ID1*, *Smad1*, and *Smad5* were ventral [**Figure S17C**]. Together, these patterns reveal asymmetric BMP signaling across the DV axis, defining a locally specialized signaling environment around the Fgf8 spot.

### Analysis of gene expression patterns across chicken, mouse and human retinas

Having reconstructed the spatial patterning of the chicken retina from scRNA-seq data, we applied this framework to additional species to explore conserved and divergent features of retinal patterning. To this end, publicly available scRNA-seq datasets of human and mouse RPCs were analyzed (see Methods). The developmental stages selected (PCW7-10 for human and E12-E14 for mouse) correspond to early retinal development, when RPC populations predominate and regional patterning programs are already established, or are in the process of being established/refined (13, 41).

To investigate patterning in human tissue, whole-retina scRNA-seq datasets were assembled from three studies covering PCW7-10, yielding 21,793 RPCs (see Methods). DV.Score and NT.Score were computed using the same axis-biased gene sets applied to chicken, and cells were mapped along DV and NT axes using these spatial coordinates [**Figure S18C and D**]. The resulting 2D topographic reconstructions revealed both concordant and divergent spatial domains compared to chicken [**Figure 7**]. For example, *Cyp26c1,* which is nasally enriched in chicken, displayed a more temporally shifted expression in human, whereas *Bmp2,* present as a central stripe with HAA-specific enrichment in chicken, showed ventral enrichment in human. These reconstructed maps were consistent with previously reported ISH data, which showed temporal expression of *Cyp26a1*, dorsal expression of *Aldh1a1*, and ventral expression of *Aldh1a3* in the developing human retina [**Figure 7**] (13). Moreover, the spatial autocorrelation analysis identified ten distinct spatial expression clusters, several of which extended beyond the primary DV and NT patterns to reveal finer regional specializations within the developing human retina. [**Figure S19B**].

**Figure 7.**
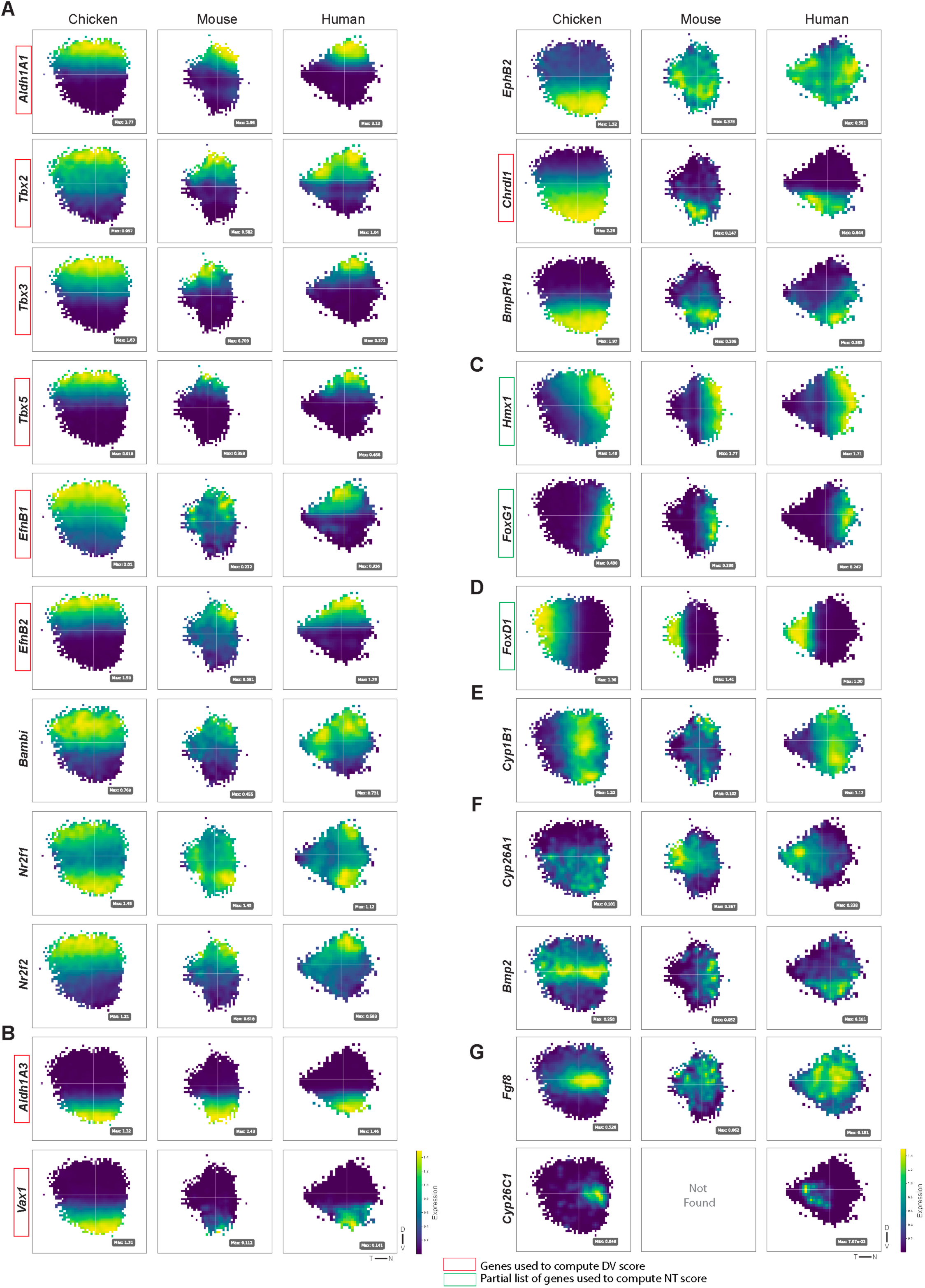
2D topographic maps for chicken, mouse and human retinas generated from scRNA-seq data. **(A-G)** 2D topographic maps of gene expression from human, mouse and chicken retinas. Each row shows the constructed 2D topographic expression map from scRNA-seq from the three species. (A) Dorsally restricted genes: *Aldh1a1, Tbx2, Tbx3, Tbx5, EphrinB1 (EfnB1), and EphrinB2 (EfnB2)* (B) Ventrally restricted genes: *Aldh1a3, Vax, Ventroptin (Chrdl1), and EphB2* (C) Nasally restricted genes: *FoxG1, and SOHo1* (D) Temporally restricted genes: *FoxD1* (E) Centrally restricted gene along the NT axis: *Cyp1b1* (F) Centrally restricted gene along the DV axis: *Cyp26a1* and *Bmp2* (G) HAA-specific genes: *Fgf8* and *Cyp26c1.* “Max” refers to the gene expression value used to normalize the upper limit of the viridis color scale. Red rectangle marks the genes used to calculate the DV score; green rectangle marks the partial list of genes used to calculate the NT score. D, Dorsal; V, Ventral; N, Nasal; T, Temporal

To investigate patterning in mouse tissue, 25,202 RPCs spanning E12-E14 were assembled from two independent datasets and processed identically to chicken and human samples (see Methods) [**Figure S18A and B**]. Spatial autocorrelation analysis detected only six significant clusters, suggesting a reduction in fine spatial organization relative to chicken and human [**Figure S19A**]. Expression of canonical HAA-associated markers, such as *Cyp26c1* and *Fgf8,* did not show a topographically enriched pattern among mouse RPCs, consistent with the absence of a HAA in the mouse retina [**Figure 7**]. Overall, the distinct clusters that emerged primarily reflected broad DV and NT axes, with little evidence of additional fine-scale patterning beyond these domains [**Figure S19A**].

Given the critical role of *Fgf8* in HAA development, the spatial expression profiles of Fgf signaling components and downstream effectors were examined in mouse and human retinas [**Figure S20, Figure S21**]. In both species, no ligand, receptor, or downstream effector displayed the nasal-central restricted pattern characteristic of *Fgf8*. In the human retina, *Fgf13*, *Fgf14*, and *Fgf19* (corresponds to *Fgf15* in mouse) were enriched nasally, while *Fgfrl1* showed enrichment in the temporal region. However, in the mouse retina, expression patterns were largely diffuse, with some genes showing temporal-central enrichment. Interestingly several downstream effectors of *Fgf8* exhibited enrichment in the temporal human retina. We also investigated spatial expression profiles of Bmp pathway components in mouse and human retinas [**Figure S22**]. Notably, neither species displayed a chicken *Bmp2*-like pattern with central localization and enrichment within the HAA. However, extracellular BMP antagonists showed enrichment in the temporal region of the human retina, suggesting localized inhibition or fine-tuning of BMP activity in the area.

### Identifying novel markers of the developing human HAA

We previously reported that *Cyp26a1* showed localized temporal expression in the developing human retina, correlating with the location of the future HAA (13). Consistent with this, the 2D topographic maps recapitulated robust *Cyp26a1* enrichment in the temporal human retina [**Figure 7**]. Additionally, *Cyp26c1* expression, though low, was specifically restricted to the *Cyp26a1*-enriched temporal domain [**Figure 7**]. To test whether *Cyp26c1* marks the developing human HAA, similar to the pattern observed in the chicken, its expression was assayed in nasal-temporal sections from Carnegie Stage 20 human retinas. Consistent with the 2D topographic maps, *Cyp26c1* expression was observed in a nested subdomain within the *Cyp26a1* territory in the temporal retina [**Figure 8A**]. The limited domain of expression for *Cyp26c1* is consistent with its low transcriptomic abundance and underscores the ability of these 2D topographic maps to capture subtle and highly localized expression patterns. Differential expression analysis was carried out to identify spatially concordant and discordant genes across retinal regions in the human retina [**Figure S23**]. Using *Cyp26c1* and *Cyp26a1* as potential HAA markers, novel genes with closely correlated expression profiles were identified, including *Nkd2, NPW, Tmem37, Dact3,* and *Gdpd2* [**Figure 8B**]. Together, these correlations point to a broader transcriptional program associated with temporal retinal specialization and the development of the HAA.

**Figure 8.**
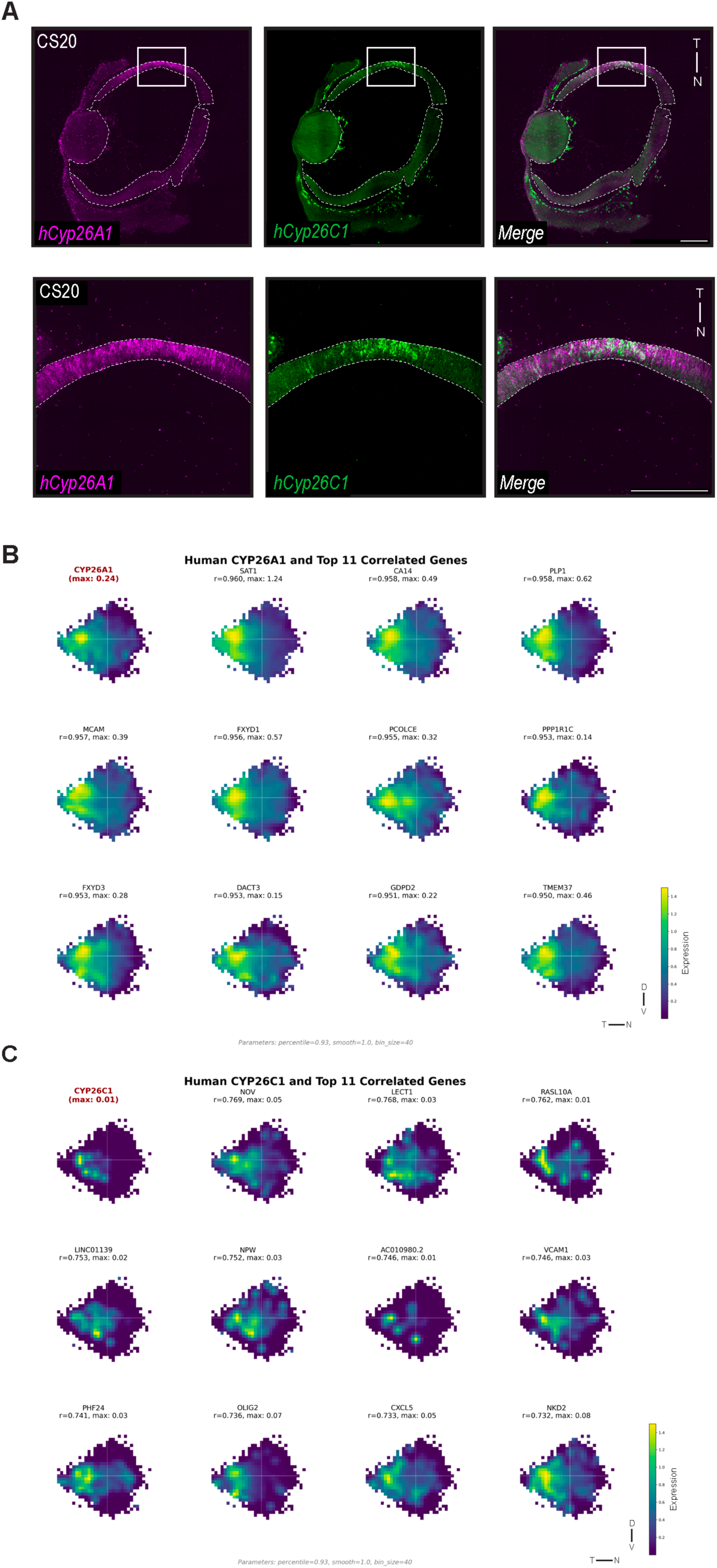
Expression patterns of genes marking the developing HAA in the human retina. **(A)** Multiplexed RNA-FISH on NT cross-sections of a CS20 human retina, showing expression of *Cyp26c1* and *Cyp26a1*. A magnified view of the temporal region, corresponding to the presumptive macula, as marked by *Cyp26a1*, is also shown. **(B-C)** 2D topographic maps of genes most strongly correlated with (B) *Cyp26a1*, and (C) *Cyp26c1* spatial patterns in the human retina. “Max” refers to the gene expression value used to normalize the upper limit of the viridis color scale. r = Pearson correlation coefficient: perfect positive correlation (r = 1), no correlation (r = 0), perfect negative correlation (r = −1). Scale bars, 200 μm. D, Dorsal; V, Ventral; N, Nasal; T, Temporal; CS, Carnegie Stage.

## DISCUSSION

The vertebrate retina provides a powerful system to study how localized territories emerge during development, with the HAA standing out as a particularly striking example. In this study, we examined the patterning landscape of the developing retina relative to the position of the HAA, leveraging the chicken as a tractable model to establish ground-truth data for cross-species reconstruction of retinal patterning. By integrating spatial and transcriptomic analyses, this work establishes a framework for understanding the developmental mechanisms that underlie HAA development. The analysis also revealed many additional 2D topographic domains, likely used by the retina to establish other spatially patterned functions, e.g. 2D topographic maps for projections to different brain regions. These methods should be applicable to any tissue to define gene expression patterns of relevance to its development and function.

### Molecular mechanisms underlying HAA specialization

Overlapping expression patterns likely act in networks that confer positional identity and refine the location of specialized areas, such as the HAA, during development. Several genes expressed very early in development establish broad dorsal, ventral, nasal, and temporal axes. While these overlapping gradients may set the foundation for further patterning, local cross-regulation and feedback are likely necessary to precisely position the HAA. Genes with early specialized expression in the developing HAA include, but might not be limited to, *Cyp26a1, Cyp26c1, Tbx3, EphrinB1, Soho1, Fgf8*, and *Bmp2*. Additional genes such as *Tbx2, Cyp1b1*, and *FoxG1* are expressed in patterns that abut or overlap the borders of the developing HAA. In addition to marking axis identities, these genes may refine the boundaries of the developing HAA and reinforce its spatial position. Yet these overlaps and boundaries alone do not fully explain how the HAA is precisely positioned - overlapping gradients may set initial conditions, but local cross-regulation and feedback are likely necessary to sharpen boundaries and define the HAA with precision.

Among these early patterning genes, the Tbx family of transcription factors have been shown to play a pivotal role in DV patterning. *Tbx2*, *Tbx3*, and *Tbx5* are expressed in overlapping dorsal patterns, but each have distinct ventral boundaries, consistent with a potential role in refining positional domains (67). These transcription factors are also known to interplay with Fgf and RA signaling. Studies in *Xenopus laevis* have shown that *Tbx2* can act as a negative regulator of *Fgf8* by suppressing *Flrt3*, a known enhancer of Fgf signaling (68). Moreover, *Tbx5* has been linked to the modulation of *Fgf8* indirectly through RA pathways. In mouse cardiopulmonary development, *Tbx5* promotes the expression of the RA-synthesizing enzyme *Aldh1a2* (also known as *Raldh2*), which in turn represses *Fgf8* activity (69). Perturbation of Tbx genes in the early chicken retina provides direct evidence for the roles of these Tbx genes in retinal patterning (70). We showed that the Tbxs regulate each other, as well as RA pathway genes and *Fgf8*. Examination of late-stage chicken embryos revealed that these perturbations affect the patterning of photoreceptors, consistent with earlier work showing the RA and *Fgf8* are required for HAA development (13).

The BMP pathway is likely another key component of retinal patterning. First *Bmp4,* then *Bmp2,* are expressed dorsally, complementary to the ventral BMP antagonist *Ventroptin* (49). Evidence from cardiac development indicates that BMP signaling and *Tbx2* act cooperatively to regulate proliferation and restrict gene expression in a chamber-specific manner in the developing heart (71,72). In the retina, the presence of dorsally enriched BMP ligands together with ventrally localized receptors, such as *Bmpr1b*, highlights an unresolved question of how BMP signaling is coordinated across the DV axis (73,74). Wnt signaling is another fundamental pathway in early patterning. It has been shown to intersect with Tbx, BMP, and RA signaling across retinal development and to play a role in photoreceptor patterning (75,76). However, the role of Wnt signaling in establishing broad DV and NT patterning, or in defining the HAA, remains untested.

Despite these insights, the regulatory relationships among these key patterning genes remain poorly understood. Do *Vax* and *Ventroptin* function redundantly or in parallel to sharpen the ventral *Fgf8* border? Do Tbx genes act upstream to directly position *Fgf8*, and/downstream, as effectors of RA/FGF signaling? How is the central expression domain of *Cyp1B1* set up along the NT axis, with its characteristic lack of expression within the *Fgf8* stripe? Is the presence of *Fgf8* and absence of RA sufficient to set up an ectopic HAA? How are BMP and Wnt activities integrated with these transcriptional programs to set up the HAA? While resolving these questions will require systematic perturbation and integrative network modeling, our findings suggest a model in which several partially overlapping regulatory factors cooperate to generate a sharply defined HAA.

### Integrating spatial and scRNA-seq data to reconstruct retinal patterning

To resolve the spatial landscape of the developing retina, we integrated data from multiplexed RNA-FISH flat mounts with quantitative image analysis and scRNA-seq analysis. This approach bridges spatial imaging and transcriptomic profiling, moving beyond descriptive ISH observations to generate reproducible spatial heatmaps, aligned intensity profiles, and axis-specific positional scores. A critical aspect of our approach is establishing biologically-grounded coordinates to embed single-cell transcriptomes onto reconstructed spatial axes. Rather than relying on unsupervised dimensionality reduction methods that may capture technical variation or axes irrelevant to the biological question, we constructed DV and NT scores using a limited set of marker genes with well-characterized spatial expression patterns identified by RNA-FISH experiments. This allowed us to reconstruct maps that reflect anatomical axes relevant to retinal patterning. The resolution of these spatial reconstructions scaled with dataset size. By assembling a large collection of RPC transcriptomes from each species at the relevant time points – 23,706 RPCs from chicken, 25,202 from mouse, and 21,793 from human – we achieved sufficient sampling density for resolution of fine-scale spatial domains, such as the HAA, that would not be identified in smaller datasets. Our spatial embedding technique, combined with adequate cell numbers, enabled clustering of genes by distinct spatial patterns, allowing identification of new potentially relevant genes by their predicted expression patterns. Such an approach complements spatial transcriptomic methods that face limitations in resolution, gene coverage, or cell segmentation. In addition, and importantly for many labs, this analytic method can be applied to publicly available scRNA-seq data, reducing the cost to near zero.

Spatial correlation analysis using both RNA-FISH as well as the scRNA-seq derived 2D topographic patterns revealed tripartite DV and NT architectures and their asymmetric relationships to the central FGF/BMP/RA-free hub. The close agreement of gene–gene correlation patterns obtained from these two orthogonal approaches supports the validity of the reconstruction framework and demonstrates that spatial expression relationships defined by RNA-FISH are preserved at the transcriptome level. This concordance enables extension of the analysis to many additional genes not experimentally assayed by RNA-FISH, whose relative expression profiles along the DV and NT axes can now be inferred from spatially reconstructed transcriptomes. Moreover, by applying this framework across developmental stages and integrating existing retinal scRNA-seq datasets, spatial expression dynamics can be reconstructed at high resolution. Together, these results establish a generalizable strategy for leveraging limited, multiplexed spatial measurements to transform conventional scRNA-seq data into a high-content, spatially resolved atlas, greatly expanding the scope of genes for which 2D topographic expression can be interrogated.

### Cross-species conservation and divergence of retinal patterning

This study places the chicken HAA within a broader comparative context and identifies both conserved and divergent aspects of retinal patterning across vertebrates. Several molecular features are remarkably conserved across the chicken, mouse and human retina. In both chicken and human retina, RA pathway genes mark the developing HAA, underscoring a potentially shared strategy for establishing this specialized region (13,9,33). Similarly, transcriptional programs that identify cardinal axes, such as Tbx, *FoxG1/FoxD1*, and *Vax* have specific aspects to their expression patterns that may enable refinements of central, specialized territories in chick and possibly in human. Yet, our comparative reconstructions highlight key species-specific distinctions: human RPCs express *Cyp26c1* and *Cyp26a1* in the temporal region, while the chicken expresses these genes nasally. These distinct expression patterns correlate with the location of the HAA - temporal in the human retina and nasal in the chicken - consistent with a conserved role for at least some of the patterning machinery under study here. In contrast, while mice share foundational DV and NT axis-defining genes, patterned gene expression in mouse RPCs tends toward relatively broad domains, with little evidence of additional specialization, consistent with the absence of a HAA. This contrast emphasizes that while the molecular toolkit for early axis determination is conserved, its deployment can vary among species, producing distinct morphological and functional outcomes. However, one aspect of deployment that may be conserved is the way in which the topography of the visual field is transmitted to target locations in the brain. The patterning of Ephrins and Ephs appears to be conserved among all 3 species, with their expression patterns likely set by the commonalities of early axis formation (79). Together, our findings support a model in which conserved molecular modules establish the foundation for retinal polarity and specializations, with species-specific elaborations sculpting the final architecture.

Future work should extend these comparisons beyond chicken, mouse, and human to include other species with distinct retinal specializations. Raptors and anoles, for instance, possess two foveae formed through differential tissue expansion during development (45, 51, 48).These species also provide an opportunity to test whether morphogenetic features of the HAA, including asymmetric retinal growth and foveal pit, arise from conserved signaling frameworks or from distinct patterning programs that produce species-specific morphologies. Moreover, species with two foveae offer a unique context in which to investigate whether distinct molecular or mechanical mechanisms underlie the formation of central versus temporal foveae, or whether these arise through modulation of a shared developmental program. Ultimately, comparative studies across these diverse visual systems will be essential to reveal how evolution reuses, modifies, or replaces conserved molecular programs to generate the remarkable diversity of retinal architectures.

## MATERIALS & METHODS

### Animals

Embryonic retinal tissues were obtained from fertilized White Leghorn eggs (Charles River). Fertilized eggs were incubated at 38 °C with humidity around 40% and embryos were staged according to Hamburger and Hamilton staging (Hamburger and Hamilton 1951).

### Tissue preparation

Right retinas were dissected in 1X PBS and fixed overnight in 4% paraformaldehyde (v/v) at 4 °C. After fixation, retinas were washed three times for 5 minutes each in 1X PBS at room temperature (RT) before proceeding with RNA-FISH as described below. To obtain retinal cross-sections through the HAA, DV rectangles were cut out from imaged flat mounted tissue after RNA-FISH for *Fgf8*. The retinal pieces were then incubated in 30% sucrose in PBS (w/v) overnight at 4 °C, and transferred to a 1:1 solution (v/v) of 30% sucrose in PBS:OCT (Tissue-Tek® O.C.T. Compound, Product code - 4583) for 3 hours. The samples were then frozen in the same 1:1 30% sucrose:OCT solution in cryomolds and stored at −80 °C. For each dorsal-HAA-ventral sample, 30 μm retina cryosections were collected and placed onto a cold superfrost plus microscope slide (Fisherbrand) within the cryostat, and stored at −80 °C.

### HCR *in-situ* hybridization on retinal whole-mounts

HCR in-situ hybridization was performed as described previously with some modifications (8,9). HCR v3.0 probes were manufactured by Molecular Instruments based on the sequences provided below. Fixed retinas were permeabilized in 70% ethanol/PBS for 2-4 hours at RT, incubated in wash buffer at RT for 10 minutes, incubated in hybridization buffer at 37°C for 30 minutes, and hybridized in 6nM probe solution overnight at 37 °C. The next day, retinas were washed in washing buffer twice at RT for 30 minutes each, washed in 5X SSCT with 0.1% Tween-20 twice at RT for 20 minutes each, and incubated in amplification buffer at RT for 10 minutes. Appropriate pairs of amplifier hairpins were heated at 95 °C for 90 seconds, incubated at RT for 30 minutes in the dark, and combined with amplification buffer at a 48nM concentration of each hairpin. To detect the probes, retinas were incubated in the resulting amplification solution overnight at RT. The following day, retinas were washed in 5X SSCT with 0.1% Tween-20 twice at RT for 30 minutes each. Subsequently, retinas were mounted and imaged. We also performed repeated rounds of HCR RNA-FISH to enable visualization of more probes on the same sample. This required removal of the prior set of probes using DNase. After the first round of imaging, coverslips were removed in PBS, retinas were demounted from the slide and treated with DNase (Roche, 4716728001) (1:50 dilution, for example, mix 30 mL of DNase (10 U/μL) with 150 mL of 10X buffer and 1,320 μL of distilled water) for 1 hour at 37 °C. After one hour, the existing DNase was removed, and retinas were incubated in new DNase O/N at 37 °C. The next day, retinas were washed twice with 2X SSC containing 65% formamide (Millipore Sigma, F9037) (for example, mix 1 mL of 20X SSC with 6.5 mL of formamide and 2.5 mL of distilled water) for 30 minutes each at 37 °C. Then, retinas were washed with 5X SSCT for 30 minutes at RT. The retinas then underwent a second round of HCR RNA-FISH, skipping the ethanol incubation step, instead starting with the first incubation in wash buffer, and continuing as described above.

| Species | Gene | NCBI Accession Number |
| --- | --- | --- |
| *Gallus gallus domesticus* | *Fgf8* | NM_001012767.2 |
|  | *Aldh1a1* | NM_204577.5 |
|  | *Aldh1a3* | NM_204669.3 |
|  | *Cyp26a1* | NM_001001129.2 |
|  | *Cyp26c1* | XM_040675448.2 |
|  | *Tbx2* | XM_001235320.7 |
|  | *Tbx3* | NM_001270878.2 |
|  | *Tbx5* | NM_204173.1 |
|  | *Vax* | NM_204799.3 |
|  | *Ventroptin* | NM_204171.2 |
|  | *FoxD1* | NM_205192.3 |
|  | *FoxG1* | NM_205193.1 |
|  | *SOHo-1* | NM_205386.2 |
|  | *Cyp1B1* | XM_015283751.4 |
|  | *Bmp2* | NM_001398170.1 |
|  | *Visinin* | NM_205514.1 |
|  | *EphrinB1 (EfnB1)* | NM_205035.3 |
|  | *EphrinB2 (EfnB2)* | NM_204824.2 |
|  | *EphB2* | NM_206951.4 |
| *Homo sapiens* | *Cyp26c1* | NM_183374.3 |
|  | *Cyp26a1* | NM_000783.4 |

### HCR RNA-FISH on retinal cross-sections

RNA-FISH was performed on retinal cross-sections using the HCR method as described previously with minor modifications for sectioned tissue (8,9). Sections were briefly rinsed in PBS, hydrated in PBS for 5 minutes, air-dried for 10 minutes, or until slides were visibly dry, and rehydrated in PBS for an additional 5 minutes. Samples were then incubated in 70% ethanol for 1 hour at room temperature to permeabilize tissue. All subsequent hybridization, amplification, and washing steps were otherwise identical to those described above for HCR RNA-FISH on retinal whole-mounts. For all steps, 250–300 µL of reagent was applied per slide. During overnight hybridization and amplification incubations, sections were placed in humidified chambers (Thomas Scientific MS-9020) at 37 °C and covered with hydrophobic covers (Grace Bio-Labs HybriSlip™ Hybridization Covers) to minimize evaporation.

### Microscopy

Samples were imaged with a Yokogawa CSU-W1 single disk (50mm pinhole size) spinning disk confocal unit attached to a fully motorized Nikon Ti2 inverted microscope equipped with a Nikon motorized stage and an Andor Zyla 4.2 plus sCMOS monochrome camera using a Plan Apo λ 20x/0.8 DIC I and Plan Fluor 40x/1.3 Oil DIC H/N2 objective lens. Nikon Elements AR 5.02 acquisition software was used to acquire the data. Z-stacks were acquired using Piezo Z-device, with the shutter closed during axial movement. Data were saved as ND2 files. Image files were analyzed in FIJI. For each sample, maximum projection images were generated using z-planes containing clear in-situ hybridization signal. Retinal whole-mount images were edited in FIJI, using first the “Remove Outliers” function (radius: 2 or 5 pixels, threshold: 50 raw intensity units) to eliminate bright spots, then a Gaussian Blur filter with a radius of 5 pixels was applied to improve visibility of punctate signal, and finally the mean grey value of the background was subtracted from all pixels in the image. Brightness and contrast of widefield and maximum projection images were edited to optimize visibility of signal, as assessed by eye. Schematic diagrams were created with BioRender (www.biorender.com).

### Quantification of RNA-FISH signal

To complement qualitative assessment of multiplexed RNA-FISH, we developed a custom MATLAB pipeline (Insert code name/link) to quantify transcript distribution across retinal flat-mounts while preserving spatial information.

#### Software and reproducibility

Analyses were performed in MATLAB (Version R2024B), using the Image Processing Toolbox (for blockproc) and custom colormap functions (magma) provided in the repository. The full code and required functions are available at [link to be added here].

#### Pre-processing

All retinal flat-mount images were oriented prior to analysis such that the nasal retina was consistently placed to the screen-right, the ventral retina in the bottom of the screen, and the optic fissure aligned at ∼90° to serve as a consistent anatomical landmark. This ensured that all samples were registered into a common orientation prior to quantification. Raw TIFF images were cropped (1 pixel = 0.3248 µm). Binary mask files were generated to restrict analysis to retinal tissue and exclude background regions. These masks were applied to all subsequent analyses.

#### Block-wise quantification

Each masked image was divided into square bins (25×25 pixels by default). This bin size was chosen empirically to balance spatial resolution with signal-to-noise. Within each bin, the mean fluorescence intensity was calculated. Fluorescence intensities across all bins were then normalized to a [0, 1] scale. Heatmaps were generated with a perceptually uniform colormaps (magma) and exported at 300 dpi resolution [**Figure S1].**

#### Axis-specific normalization

To highlight expression asymmetries across anatomical axes, intensity matrices were normalized column-wise and row-wise, producing maps that emphasized DV or NT gradients, respectively [**Figure S1].**

#### Origin alignment and profile extraction

A user-defined coordinate system was implemented by selecting a reference point (0,0), typically centered on the HAA. Using this origin, 500 µm-wide strips were extracted along both the x (NT) and y (DV) axes. Mean intensity profiles and standard deviations were computed across these x (NT) and y (DV) strips, yielding 1D profiles of transcript intensity distribution [**Figure S1].** Profiles were exported as Excel tables with distance (µm), normalized intensity values and standard deviation for downstream statistical analysis. When extracting intensity profiles, we were mindful of edge effects, as peripheral regions often contained few signal datapoints and a predominance of background noise. To address this, we included only those where at least 80% of the values represented signal rather than background. However, for six specific flat mounts (a sample each for *Tbx2, Tbx5, Ventroptin, FoxD1, FoxG1, Cyp1b1*), we relaxed this threshold due to the nature of the retinal cuts in those preparations.

#### Cross-sample alignment

Each retinal flat mount was reorganized, with the coordinate system adjusted according to the user-defined origin reference point, enabling heatmaps and one-dimensional intensity profiles to be aligned and compared across flat mounts. Data from individual flat mounts were then linearly interpolated onto a common spatial axis. By overlaying intensity profiles from independent samples within this unified coordinate system, expression boundaries could be directly compared and reproducibility across replicates. For each transcript, the combined mean and standard deviation intensity profiles could be calculated. Aligned profiles highlighted conserved boundary positions and permitted averaging across multiple flat mounts to generate representative maps of spatial gene expression patterns.

#### Validation and reproducibility

Quantified heatmaps were visually compared to raw RNA-FISH images to ensure fidelity of expression patterns and boundaries, and the two were highly concordant. Analyses were performed on at least three independent whole-mounts per gene (N = 3), yielding reproducible maps and profiles for *Fgf8, Aldh1a1, Aldh1a3, Cyp26a1, Cyp26c1, Tbx2, Tbx3, Vax, Cyp1b1, FoxD1, FoxG1, SOHo1* or two independent experiments (N = 2) for *Tbx5, Ventroptin*.

#### Output

For each image, the pipeline produced (i) heatmaps of normalized expression, (ii) column- and row-normalized maps, (iii) intensity profiles across chosen axes, and (iv) tabulated numerical values of distance versus intensity. This framework enabled reproducible quantification of spatial transcript distributions and direct comparison of expression boundaries across multiple HCR datasets.

### 2D topographic maps using chicken, human, and mouse single-cell RNA-seq datasets

To reconstruct spatial expression patterns from dissociated single-cell data, chicken, human, and mouse RPC cell transcriptomes were processed through a standardized pipeline. Quality-controlled datasets were scored for DV and NT identity, discretized into spatial grids, and smoothed to produce 2D topographic maps. This framework enabled direct comparison of spatial expression modules within and across species, and highlighted conserved as well as divergent gene expression patterns throughout retinal development.

#### Processing of single-cell RNA-seq datasets from chicken, human and mouse

##### Processing of chicken developing retina scRNA-seq datasets

Fifteen in-house libraries were generated from E5 - E6 chicken retinas using the 10X Genomics Chromium v2 and v3.1 single-cell protocols (single and dual index). Embryos were cultured ex ovo for 1-2 days prior to dissociation. Some libraries contained retroviral GFP barcode infections or CRE-GFP plasmid transfections for unrelated studies; reads corresponding to these manipulations were excluded. An additional publicly available data set (GSE142244) was included, reprocessed starting from FASTQ files to generate relevant count matrix (Cellranger v9.0.0) with the identical chicken reference genome [**Figure S12**].

From the generated count matrix, doublets were identified and removed using scDblFinder (21). The doublet-filtered dataset was subjected to SNP detection with cellsnp-lite (v1.2.3) and genotype-based demultiplexing with vireoSNP (v0.5.8) for further doublet removal, and identification of unique genotypes. For genotype-based demultiplexing, at least 100 different random seeds were applied to obtain the best fit. The identity of unique genotypes (used for pseudobulk later) were validated with the sex chromosome genes (W/Z). Simple intron and exon ratio was computed to identify droplet transcriptomes that had low intron presence to further filter putative debris (40). The amount of ambient RNA was assessed using SoupX (v1.6.2, 66), but found to be similar across libraries and at low level, so no further action was taken to correct for the ambient RNA effect.

Among initial 110,061 cells, 85,135 single-cell transcriptomes were subject to standard processing in Seurat v5 framework (24, 58, 6, 52, 77) for normalization, integration (Harmony - 32), and graph-based clustering (Leiden - 61). For clustering, cell cycle score (G2M/S Score, computed from each library separately), mitochondrial content, and W/Z gene content were regressed out. Leiden clustering (resolution = 0.1) showed good demarcation of cell clusters, including the RPC populations, and identification of cell transcriptomes with signatures of viral infection (due to retroviral barcode infection), which were filtered out.

The resulting RPC population amounted to 23,706 single-cell transcriptomes, derived from 30 individual embryos (based on inferred genotypes by vireo) (78). From the data, the DV.Score and NT.Score were computed using the AddModuleScore function in Seurat, with the combination of {*Tbx5, Tbx2, Tbx3, Aldh1a1, EfnB2, EfnB1*}, {*Vax1, Chrdl1, Aldh1a3*}, {*FoxG1, SOHo-1, Hmx1, EfnA5, EfnA2*}, and {*FoxD1, EphA3*} respectively [**Figure S13, Figure S14**]. Subsequently, the DV.score and NT.score were calculated by subtracting the Ventral score from the Dorsal score, and the Temporal score from the Nasal score. There was little difference of score distribution whether the score was computed in library or technology level (10X v2, v3.1) or all together, so the integrated dataset as a whole was used to compute the score.

##### Processing of human developing retina scRNA-seq datasets

Only whole retina samples between PCW7-11 were used for integration (GSE138002, GSE234963, GSE246169). The publicly available count matrices were merged, normalized, integrated, and clustered following the same pipeline as for chicken mentioned above. Because mammals exhibit dosage compensation of sex chromosome genes, W/Z regression was not required. After clustering, RPCs were identified based on the publicly available meta information overlap (RPCs and mitotic cell population, which overlap with RPC cluster after cell-cycle regression; 21,793 cells). For the RPC population, the same set of genes as used above for chicken analysis were used to construct the DV.Score and NT.Score [**Figure S18**].

##### Processing of mouse developing retina scRNA-seq datasets

Publicly available dataset from GSE139904 (E13; total 11,239 cells) and GSE118614 (E14; 26,905 cells, E16; 5,382 cells) were processed accordingly. The count matrices were merged after harmonization of gene names. Mitochondrial content and cell-cycle scores were regressed out during the normalization and scaling process. After clustering, RPCs (25,202 cells) were identified based on the publicly available meta information overlap and was used to construct the DV.Scores and NT.Scores with the same orthologous gene sets as chicken and human [**Figure S18**].

#### Spatial gene expression analysis for chicken, human and mouse

##### Spatial binning and smoothing

Spatial analyses were performed in Python (v3.11) using scanpy (v1.9.6), anndata (v0.10.3), and custom modules (https://github.com/chlee-tabin/retina-spatial-scrna-analysis). To reconstruct spatial gene expression patterns, the continuous DV and NT coordinates were discretized into regular 2D grids, where each grid cell represents an equally sized spatial region. Species-specific grid resolutions were optimized based on cell density and spatial precision requirements.

To evaluate how grid resolution affects reconstructed spatial patterns, coarse (20×20) and fine (100×100) grids were compared using *Cyp26c1,* which shows a highly specific spatial pattern but is expressed at low levels in our dataset, and *Fgf8*, which is both highly specific and robustly expressed [**Figure S15**]. The 100×100 grid preserved fine spatial detail but introduced considerable noise, whereas the 20×20 grid reduced noise but oversmoothed the expression patterns.

- Chicken: We selected an intermediate grid resolution of 51×51 cells (2,601 total), with minimum thresholds of ≥3 cells and ≥20 gene counts per grid cell.
- Human: We used a 40×40 grid (1,600 cells) with thresholds of ≥3 cells and ≥15 gene counts per grid cell.
- Mouse: Due to higher cell density, we used a 51×51 grid with thresholds of ≥5 cells and ≥30 gene counts per grid cell.

Gaussian smoothing (σ = 1.0) was applied to the binned expression values to reduce noise while preserving spatial patterns. Expression values exceeding the 93rd-95th percentile (species-dependent) were clipped to minimize outlier effects in visualization.

##### Spatial pattern detection

We identified spatially variable genes using a custom greedy algorithm that maximizes spatial autocorrelation [**Figure S17, Figure S19**]. For each gene, we calculated Moran’s I statistic to quantify spatial clustering. Genes with significant spatial variance (p < 0.01 after Benjamini-Hochberg correction) were retained for pattern analysis. Spatial similarity between genes was assessed using cosine similarity of their smoothed expression patterns. Genes with correlation coefficients > 0.9 were considered to have highly similar spatial distributions. Pre-defined spatial anchor genes were selected among the genes below with the highest expression per species to orient patterns along anatomical axes (Dorsal: *Aldh1a1, Bambi*; Ventral: *Aldh1a3, Vax1*; Temporal: *EphA8, EphA2, EphA3, FoxD1*; Nasal: *FoxG1, Hmx1* for human/chicken with appropriate orthologs for mouse). “Max” refers to the gene expression value used to normalize the upper limit of the viridis color scale. Rather than using the absolute maximum, which can be skewed by a few outlier bins, normalization is based on a specified percentile cutoff (p). The percentile cutoff defines the expression value at which the color scale reaches its maximum intensity. All values above this threshold are displayed using the brightest color, ensuring that extreme expression levels do not compress the dynamic range of the visualization and that spatial differences in the majority of the data remain visible. “r” refers to the Pearson correlation coefficient - linear similarity between two sets of spatial patterns; r = 1.0 → perfect positive correlation (patterns are nearly identical); r = 0 → no correlation.; r = –1.0 → perfect negative correlation (patterns are mirror-opposites).

##### Differentially expressed genes in regions

Regions of interest (e.g., HAA, nasal-most, temporal-most) were defined by bins enriched for anchor gene expression, as noted in the figure [**Figure 6H**]. The differentially expressed gene analysis used glmGamPoi (1) to first pseudobulk the single cell transcriptomes based on the defined region. Only pseudobulks containing more than 100 cells were used.

#### Cross-species comparative analysis of spatial gene expression patterns

##### Orthology mapping

To enable cross-species comparisons, we mapped genes between species using a dual-database approach. Primary orthology information was retrieved from MyGene.info API (accessed via mygene Python client v3.2.2), with Ensembl BioMart serving as a fallback source. Species were identified using NCBI taxonomy IDs: 9606 (human), 10090 (mouse), and 9031 (chicken).

#### Statistical analysis for 2D topographic maps generated from scRNA-seq datasets

##### Correlation and Similarity Metrics

Pearson correlation coefficients were calculated for all pairwise gene comparisons within spatial bins. Spatial patterns were considered significantly correlated at r > 0.9 (p < 0.001). Cosine similarity was used as an additional metric for high-dimensional expression pattern comparisons.

##### Multiple Testing Correction

All statistical tests involving multiple comparisons were adjusted using the Benjamini-Hochberg false discovery rate (FDR) method with a significance threshold of q < 0.05.

#### Software and Reproducibility

Complete analysis code is available at https://github.com/chlee-tabin/retina-spatial-scrna-analysis. Processed data files containing expression matrices, spatial coordinates, and cell annotations are deposited at GEO (GSEXXXXX).

#### Data Availability

Raw sequencing data and processed expression matrices are available through GSEXXXXX.

### Spatial correlation analysis of quantitative RNA-FISH and scRNA-seq data

To quantify pairwise relationships between genes and identify co-expressed modules, we first interpolated each gene’s spatial profile onto a common coordinate grid and then performed correlation-based hierarchical clustering. These interpolated profiles were assembled into a gene-by-position matrix. Pairwise Pearson correlation coefficients were then computed between all gene profiles, using only positions where both genes had non-NaN values; this yielded a symmetric correlation matrix that captures the similarity of spatial expression patterns.

To cluster genes, the correlation matrix was converted to a distance matrix (distance = 1 − correlation). This distance matrix was supplied to hierarchical clustering using average linkage. Optimal leaf ordering was applied to the resulting dendrogram to minimize within cluster distances along the heatmap axes. The clustered correlation matrix was visualized as a heatmap with genes ordered according to this dendrogram, revealing groups of genes with highly similar spatial profiles. This procedure was applied to a selected subset of genes (DV and NT genes) to generate the focused heatmaps and clustered correlation matrices shown in **Figure 4, Figure S13E, and Figure S14E.**

## ACKNOWLEDGEMENTS

All images were acquired using Microscopy Resources on the North Quad (MicRoN) core at Harvard Medical School. We thank Paula Montero Llopis, Praju Vikas Anekal and Adrienne Wells for their assistance with microscopy and helpful discussions. We thank Susana da Silva for providing human tissue and feedback.

## SUPPLEMENTARY FIGURES

**Figure S1. Quantification of RNA-FISH signal for Fgf8**

**(A)** Normalized mean intensity heatmap showing *Fgf8* expression. **(B)** Column- and row-normalized intensity maps for *Fgf8*. **(C)** Spatial intensity mapping and 1D expression profiling of *Fgf8*. Normalized mean intensity profiles of *Fgf8* along DV and NT axes are shown with the origin set at the center of the *Fgf8* spot. The x-axis shows distance (µm) from the center of the *Fgf8* expression spot. The y-axis shows normalized mean fluorescence intensity, calculated from RNA-FISH signal. To generate the DV and NT profiles (1D intensity profiles), a 500-µm-wide strip was extracted along the DV or NT axis, and fluorescence values were averaged across the width of the strip at each position along the axis. Black lines denote the mean normalized intensity; red lines represent the standard deviation (SD). **(D, E)** Spatial expression profiles of *Fgf8* across three HH25/26 retinas. DV and NT profiles for mean intensity profiles were generated for each sample (N = 3) after alignment to their respective origins. The averaged DV and NT profiles are shown across the three replicates. Averaged DV and NT profiles were plotted from these 3 samples. Black lines denote the mean normalized fluorescence intensity; red lines represent the standard error of the mean (SEM). Profiles were calculated from N = 3 biological replicates. HH, Hamburger and Hamilton; D, Dorsal; V, Ventral; N, Nasal; T, Temporal.

**Figure S2. Quantification of RNA-FISH signal for RA pathway genes relative to Fgf8**

Normalized mean intensity profiles of RA pathway genes along DV and NT axes with the origin set at the center of the *Fgf8* spot. From each origin, 500 µm wide strips were extracted along the x- and y-axes, and mean intensity values were computed across 1D cross-sections to create graphs of relative gene expression. Black lines denote the mean normalized fluorescence intensity; red/blue lines represent the standard deviation (SD). DV and NT mean intensity profiles for **(A)** *Aldh1A1* and *Fgf8* **(B)** *Aldh1A3* and *Fgf8* **(C)** *Cyp26a1* and *Fgf8* **(D)** *Cyp26c1* and *Fgf8.* White arrow highlights the transient “bull’s-eye” pattern of *Cyp26a1*, which aligns with the *Fgf8* peak along the DV and NT axis. HH, Hamburger and Hamilton; D, Dorsal; V, Ventral; N, Nasal; T, Temporal.

**Figure S3 Quantification of RNA-FISH signal for Fgf8 and RA pathway genes in retinal sections**

Multiplexed RNA-FISH hybridization on HH25/HH26 DV cross-sections through the HAA using probes for **(A)** *Fgf8, Aldh1a1*, and *Aldh1a3* or **(B)** *Fgf8, Cyp26a1*, and *Cyp26c1*. **(A’, B’)** Normalized mean intensity heatmaps generated using the MATLAB pipeline. Scale bars, 100µm. HH, Hamburger and Hamilton; D, dorsal; V, ventral.

**Figure S4. Quantification of RNA-FISH signal for DV genes relative to Fgf8**

Normalized mean intensity profiles of DV genes along DV and NT axes with the origin set at the center of the HAA (*Fgf8* spot). From each origin, 500 µm wide strips were extracted along the x- and y-axes, and mean intensity values were computed across 1D cross-sections to create graphs of relative gene expression. Black lines denote the mean normalized fluorescence intensity; red/blue lines represent the standard deviation (SD). DV and NT mean intensity profiles for **(A)** *Tbx2* and *Fgf8* (White arrow highlights region of peak *Tbx2* expression, which is placed right next to the *Fgf8* spot along the DV and NT axis) **(B)** *Tbx3* and *Fgf8* (White arrow highlights region of peak *Tbx3* expression, which aligns with the *Fgf8* peak along the DV and NT axis) **(C)** *Tbx5* and *Fgf8,* **(D)** *Vax* and *Fgf8, and* **(E)** *Ventroptin* and *Fgf8.* HH, Hamburger and Hamilton; D, Dorsal; V, Ventral; N, Nasal; T, Temporal.

**Figure S5. Expression domains of DV genes relative to Fgf8 at HH25/26**

Multiplexed RNA-FISH was performed on HH25/HH26 retinal whole mounts for (A) *Fgf8* and *Tbx2*; (B) *Fgf8* and *Tbx3*; (C) *Fgf8* and *Tbx5*; (D) *Fgf8* and *Vax*; (E) *Fgf8* and *Ventroptin*. Each panel shows merged images and corresponding single-channel expression images. Dorsally patterned genes (*Tbx2, Tbx3,* and *Tbx5*); Ventrally patterned genes (*Vax* and *Ventroptin*); Scale bars, 200 μm. HH, Hamburger and Hamilton; D, Dorsal; V, Ventral; N, Nasal; T, Temporal. All patterns were observed in N ≥ 3 retinas. The *Tbx2* image shown here (A) is a widefield image, whereas all other panels are maximum projections of confocal z-stacks.

**Figure S6. Expression domains of DV genes relative to Fgf8 at HH28/29**

Multiplexed RNA-FISH was performed on HH28/HH29 retinal whole mounts for (A) *Fgf8* and *Tbx2*; (B) *Fgf8* and *Tbx3*; (C) *Fgf8* and *Tbx5*; (D) *Fgf8* and *Vax*; (E) *Fgf8* and *Ventroptin*. Each panel shows merged images, and corresponding single-channel expression images. Dorsally patterned genes (*Tbx2, Tbx3,* and *Tbx5*); Ventrally patterned genes (*Vax* and *Ventroptin*); Scale bars, 200 μm. HH, Hamburger and Hamilton; D, Dorsal; V, Ventral; N, Nasal; T, Temporal. All patterns were observed in N ≥ 3 retinas. All the images are maximum projections of confocal z-stacks.

**Figure S7. Expression domains of Eph/Ephrin in the developing chicken retina**

Multiplexed RNA-FISH was performed on HH25/HH26 retinal whole mounts for (A) *EphrinB1* (B) *EphrinB2* (C) *EphB2*. Each panel shows single-channel expression images, and corresponding quantified intensity heatmaps. (D-E) Merged images with two channels for visualization of relative expression domains. Dorsally patterned genes (*EphrinB1* and *EphrinB2*); Ventrally patterned genes (*EphB2*); Scale bars, 200 μm. HH, Hamburger and Hamilton; D, Dorsal; V, Ventral; N, Nasal; T, Temporal. All patterns were observed in N ≥ 3 retinas. All the images are widefield.

**Figure S8. Quantification of RNA-FISH signal for nasal-temporal genes relative to Fgf8**

Normalized mean intensity profiles of early NT patterned genes along DV and NT axes with the origin set at the center of the HAA (*Fgf8* spot). From each origin, 500 µm wide strips were extracted along the x- and y-axes, and mean intensity values were computed across 1D cross-sections to create graphs of relative gene expression. Black lines denote the mean normalized fluorescence intensity; red/blue lines represent the standard deviation (SD). DV and NT mean intensity profiles for **(A)** *FoxD1* and *Fgf8* **(B)** *FoxG1* and *Fgf8* **(C)** *SOHo1* and *Fgf8* (White arrow highlights region of peak *SOHo1* expression, which aligns with the *Fgf8* peak along the NT axis) **(D)** *Cyp1B1* and *Fgf8.* HH, Hamburger and Hamilton; D, Dorsal; V, Ventral; N, Nasal; T, Temporal.

**Figure S9. Expression domains of nasal-temporal genes relative to Fgf8 at HH25/26**

Multiplexed RNA-FISH was performed on HH25/HH26 retinal whole mounts for (A) *Fgf8* and *FoxD1*; (B) *Fgf8* and *FoxG1*; (C) *Fgf8* and *SOHo1*; (D) *Fgf8* and *Cyp1B1* (White arrow highlights region of absence of *Cyp1B1* expression along the equator). Each panel shows merged images, and corresponding single-channel expression images. Scale bars, 200 μm. HH, Hamburger and Hamilton; D, Dorsal; V, Ventral; N, Nasal; T, Temporal. All patterns were observed in N ≥ 3 retinas. All the images are maximum projections of confocal z-stacks.

**Figure S10. Expression domains of nasal-temporal genes relative to Fgf8 at HH28/29**

Multiplexed RNA-FISH was performed on HH28/HH29 retinal whole mounts for (A) *Fgf8* and *FoxD1*; (B) *Fgf8* and *FoxG1*; (C) *Fgf8* and *SOHo1*; (D) *Fgf8* and *Cyp1B1* (White arrow highlights region of absence of *Cyp1B1* expression along the equator). Each panel shows merged images, and corresponding single-channel expression images. Scale bars, 200 μm. HH, Hamburger and Hamilton; D, Dorsal; V, Ventral; N, Nasal; T, Temporal. All patterns were observed in N ≥ 3 retinas. All the images are maximum projections of confocal z-stacks.

**Figure S11. Quantification of RNA-FISH signal for Bmp2 and Visinin relative to Fgf8**

Multiplexed RNA-FISH was performed on **(A)** E6 retinal whole mounts for *Fgf8* and *Bmp2*; and **(B)** E7 retinal whole mounts for *Fgf8* and *Visinin*. Each panel shows mean intensity profiles of these genes along DV and NT axes with the origin set at the center of the Fgf8 spot. From each origin, 500 µm wide strips were extracted along the x- and y-axes, and mean intensity values were computed across 1D cross-sections to create graphs of relative gene expression. Black lines denote the mean normalized fluorescence intensity; red/blue lines represent the standard deviation (SD). **(A’)** Column-normalized intensity maps for *Bmp2.* **(B’)** Row-normalized intensity maps for *Visinin.* E, Embryonic day; D, Dorsal; V, Ventral; N, Nasal; T, Temporal. *Bmp2* pattern was observed in N ≥ 3 retinas. *Visinin*-free spot was observed transiently and has been previously described (55).

**Figure S12. Processing of scRNA-seq datasets from developing chicken retina**

(A) UMAP plot of major cell clusters from chicken E5-6 retinal samples. (B) Violin plot of marker gene expression across the graph-based clusters (Leiden resolution=0.1). (C) UMAP plots color coded by cell cycle phase (top left), version of 10X Chromium technology (top right), mitochondrial content (bottom left), gene count per cell (bottom right). (D) Cell type composition of samples from each embryo. (E) Violin plot of W chromosome and Z chromosome gene fraction per cell for inferred individual embryos from all libraries. Note that avian transcriptomes do not display sex chromosome dosage compensation such that females (WZ) have half the Z gene expression than males (ZZ), and males do not have W gene expression.

**Figure S13. Generation of DV score using single-cell transcriptomes from the developing chicken retina**

**(A)** Distribution of Dorsal, Ventral score and the subtracted composite score DV.Score. **(B)** Distribution of Dorsal and Ventral scores. Each point represents an individual single-cell transcriptome. **(C)** Distribution of gene expression along the DV.Score axis. The grayed-out genes represent the genes used for the DV score construction. **(D)** Binned expression pattern along the DV.Score axis, with individual points representing pseudobulked expression from individual embryos. **(E)** Spatial correlation analysis of selected genes from scRNA-seq data along the DV axis. Left, heatmap showing normalized expression of selected DV genes ordered by inferred DV position. Right, clustered correlation heatmap for the same genes based on pairwise Pearson correlation of their spatial expression profiles, revealing dorsal, central, and ventral gene modules. Genes are ordered according to correlation-based hierarchical clustering. The red dashed line indicates the approximate position of the developing HAA, marked at the center of the *Fgf8* and *Cyp26C1* expression domains. D, Dorsal; V, Ventral; DV.score, Dorsal-Ventral score.

**Figure S14. Generation of NT score using single-cell transcriptomes from the developing chicken retina**

**(A)** Distribution of Nasal, Temporal score and the subtracted composite score NT.Score. **(B)** Distribution of Nasal, Temporal scores. Each point represents an individual single-cell transcriptome. **(C)** Distribution of gene expression along the NT.Score axis. The grayed-out genes represent the genes used for the NT score construction. **(D)** Binned expression pattern along the NT.Score axis, with individual points representing pseudobulked expression from individual embryos. **(E)** Spatial correlation analysis of selected genes from scRNA-seq data along the NT axis. Left, heatmap showing normalized expression of selected DV genes ordered by inferred NT position. Right, clustered correlation heatmap for the same genes based on pairwise Pearson correlation of their spatial expression profiles, revealing nasal, central, and temporal gene modules. Genes are ordered according to correlation-based hierarchical clustering. The red dashed line indicates the approximate position of the developing HAA, marked at the center of the *Fgf8* and *Cyp26C1* expression domains. N, Nasal; T, Temporal; NT.score, Nasal-Temporal score.

**Figure S15. Grid size sensitivity analysis for 2D topographic map reconstruction**

Panels show reconstructed expression maps computed using two grid resolutions: a fine grid (100 × 100 subdivisions) and a coarse grid (20 × 20 subdivisions). Each grid cell represents the mean expression value of all points falling within that spatial region. Maps are shown for *Cyp26c1*, which exhibits a highly specific spatial pattern but is expressed at low levels in the dataset, and *Fgf8*, which is both highly specific and robustly expressed. “Max” refers to the gene expression value used to normalize the upper limit of the viridis color scale. Max denotes the percentile-based upper limit of the color scale (93rd–95th percentile, species-dependent), not the absolute maximum expression. D, Dorsal; V, Ventral; N, Nasal; T, Temporal; DV.score, Dorsal-Ventral score; NT.score, Nasal-Temporal score.

**Figure S16. Identification of spatial expression clusters in the chicken retina**

**(A)** 20 distinct spatial expression patterns (anchors) that maximize the divergence of spatial expression pattern in chicken scRNA-seq datasets. Each anchor represents a characteristic spatial profile, and genes were assigned to anchors based on similarity of their reconstructed expression patterns. Anchors that shared mutual nearest neighbors were grouped, with the associated number of genes indicated. Clusters manually annotated based on known marker gene expression are labeled and highlighted in yellow. The bottom-right square within each panel (p) shows the percentile cutoff used for visualization, where bins at or above the selected percentile are assigned the maximum color value to minimize distortion from outliers. Red border = forms cluster, (n) = number of genes in cluster. **(B-D)** Genes most strongly correlated with (B) *Fgf8*, (C) *Cyp26c1*, and (D) *Bmp2* spatial patterns. “Max” refers to the gene expression value used to normalize the upper limit of the viridis color scale. r = Pearson correlation coefficient: perfect positive correlation (r = 1), no correlation (r = 0), perfect negative correlation (r = −1). D, Dorsal; V, Ventral; N, Nasal; T, Temporal; DV.score, Dorsal-Ventral score; NT.score, Nasal-Temporal score.

**Figure S17. Spatial expression patterns of genes associated with Fgf, Bmp and RA signaling pathways in the chicken retina**

2D topographic maps of retinal gene expression of **(A)** *Fgf* signaling ligands and receptors, **(B)** downstream targets of *Fgf8*, **(C)** *Bmp* signaling pathway genes, **(D)** Genes most strongly correlated with the *Cyp1B1* spatial pattern. “Max” refers to the gene expression value used to normalize the upper limit of the viridis color scale. r = Pearson correlation coefficient: perfect positive correlation (r = 1), no correlation (r = 0), perfect negative correlation (r = −1). D, Dorsal; V, Ventral; N, Nasal; T, Temporal; DV.score, Dorsal-Ventral score; NT.score, Nasal-Temporal score.

**Figure S18 Generation of DV and NT scores using single-cell transcriptomes from developing mouse and human retinas**

Spatial expression pattern of marker genes used to calculate the **(A, C)** DV.Score and **(B, D)** NT.score using scRNA-seq data from developing (A, B) mouse and (C, D) human retina. From the data, D, V, N and T scores were computed with the combination of {*Tbx5, Tbx2, Tbx3, Aldh1a1, EfnB2, EfnB1*}, {*Vax1, Chrdl1, Aldh1a3*}, {*FoxG1, SOHo-1, Hmx1, EfnA5, EfnA2*}, and {*FoxD1, EphA3*} respectively. Validation of the spatial reconstruction pipeline using independent genes with well-defined axial expression patterns that were not included in the scoring set; **(A’, C’)** *Fgf8*, *Cyp26c1* and *Bmp2* for validation along DV axis and **(B’, D’)** *Fgf8*, *Cyp1B1*, and *Cyp26c1* for validation along NT axis. The histogram on the bottom represent coverage by aggregating all cells that contributed to the bins (total bins=50). The dashed lines represent the DV or NT axis bins that do not have more than 50 cells covered, suggesting less confidence for proper DV/NT.Score.

**Figure S19. Identification of clusters for spatial expression patterns in mouse and human retinas**

Spatial patterns (anchors) that maximize the divergence of spatial expression pattern were detected using (A) mouse and (B) human retinal scRNA-seq datasets. Anchors that have mutual neighbors - that is, genes exhibiting reciprocal similarity in their spatial expression patterns - were highlighted with the number of genes associated with the cluster (set of genes that share similar expression patterns). 6 main clusters were detected in mouse and 10 in the human dataset. A yellow label indicated manual annotation of certain clusters. The bottom right presents the percentile clip. Red border = forms cluster, (n) = number of genes in cluster. “Max” refers to the gene expression value used to normalize the upper limit of the viridis color scale. D, Dorsal; V, Ventral; N, Nasal; T, Temporal; DV.score, Dorsal-Ventral score; NT.score, Nasal-Temporal score.

**Figure S20. 2D topographic maps of Fgf family members in mouse and human retinal RPCs**

2D topographic maps of retinal gene expression of *Fgf* signaling ligands and receptors in retinal scRNA-seq datasets from (A) mouse and (B) human. D, Dorsal; V, Ventral; N, Nasal; T, Temporal.

**Figure S21. 2D topographic maps of genes related to *Fgf8* signaling pathway in mouse and human retinal RPCs**

2D topographic maps of retinal gene expression of downstream targets of Fgf signaling in retinal scRNA-seq datasets from (A) mouse and (B) human. D, Dorsal; V, Ventral; N, Nasal; T, Temporal.

**Figure S22. 2D topographic maps of BMP signaling pathway components in developing mouse and human retinas**

2D topographic maps of retinal gene expression of Bmp signaling pathway genes in retinal scRNA-seq datasets from (A) mouse and (B) human. D, Dorsal; V, Ventral; N, Nasal; T, Temporal.

**Figure S23. Differential gene expression analysis across different regions in the human retina**

(A) Visual representation of virtual selection of the specific regions in the 2D topographic maps to assess differentially enriched or de-enriched genes across these regions. The 2D topographic gene expression of “region-specific marker gene” was overlayed with area bins, and the selected region was overlayed with blue. Volcano plots were generated using pseudobulked gene expression samples that are enriched or de-enriched in the selected regions. Points are scaled by average gene expression level, and genes above 2-fold expression difference between in-group and out-group and adjusted p-values less than 0.05 were colored. The top 5 enriched genes without the “LOC” prefix were highlighted. See also Supplemental Table 2. Region-specific marker genes: *Aldh1a1* (Dorsal), *Bmp2* (Central domain across Dorsal-Ventral axis), *Cyp26c1* (HAA), *FoxG1* (Nasal), *Cyp1B1* (Central domain across Nasal-Temporal axis), *FoxD1* (Temporal), *Vax1* (Ventral). Scale bars, 200 μm. HAA, High(er) Acuity Area; N, Nasal; T, Temporal.

## SUPPLEMENTARY TABLE

**Supplementary Table 1 & 2. Differentially expressed genes across spatial regions of chicken retina (Supplementary Table 1) and human (Supplementary Table 2).** This table lists genes that are enriched or de-enriched within spatial domains identified in (Supplementary Table 1) Figure 6 and (Supplementary Table 2) Figure S23. Differential expression was calculated using pseudobulked transcriptomes derived from region-specific selections in the 2D topographic maps. For each region, the table reports the gene name, average expression, log₂ fold-change between in-group and out-group cells, adjusted p-value, and significance. Region-specific markers used for defining the domains include (Dorsal) *Aldh1a1*, (DV-central) *Bmp2*, (HAA) *Cyp26c1*, (Nasal) *FoxG1*, (NT-central) *Cyp1b1*, (Temporal) *FoxD1*, and (Ventral) *Vax1*.

**Supplementary Table 3. Comprehensive analysis of expression patterns of FGF and BMP pathway genes across human and mouse datasets.** The table lists all genes surveyed, indicating which genes were detected, not detected, and which exceeded the expression threshold (>0.1 TPM). Genes are categorized by detection status and expression level to identify candidates potentially involved in early patterning and signaling.

