## Supplementary figures and images for "Integration of in situ hybridization and scRNA-seq data provides a 2D topographical map of the developing retina across species"

### Supplemental Figure 1

**Supplementary Figure 1. Quantification of RNA-FISH signal for *Fgf8***

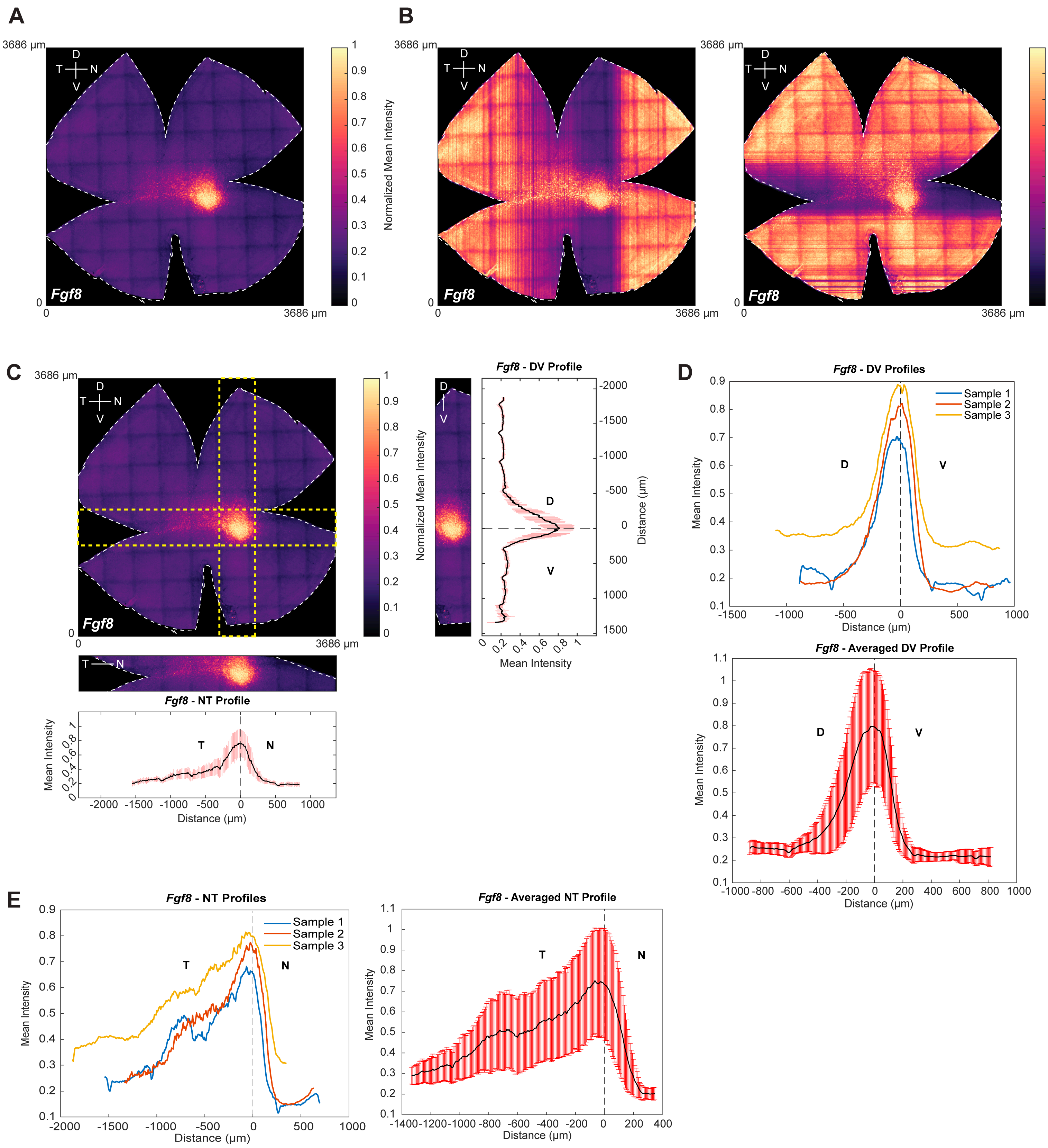

### Supplemental Figure 2

#### Supplementary Figure 2. Quantification of RNA-FISH signal for RA pathway genes

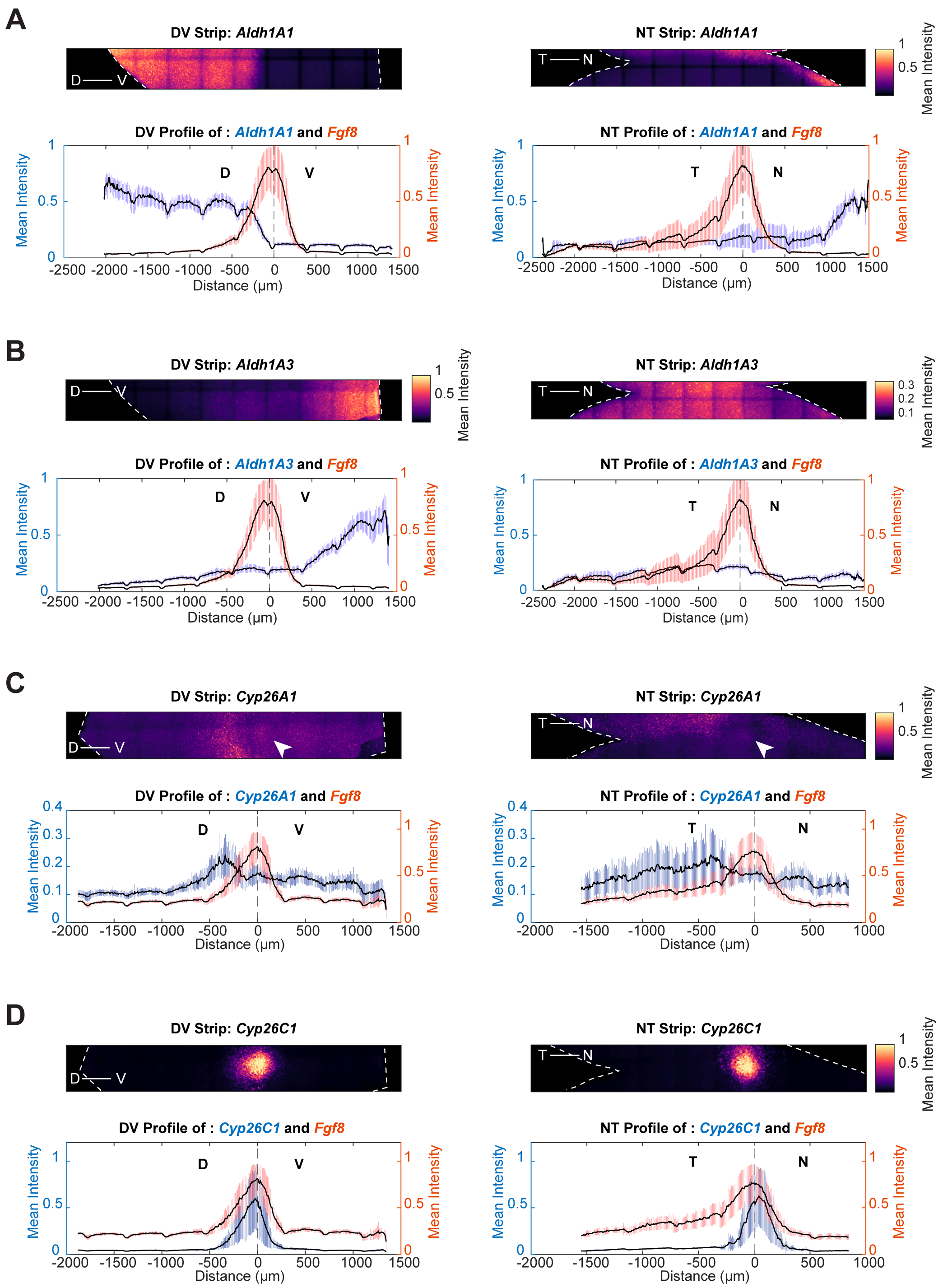

### Supplemental Figure 4

##### Supplementary Figure 4. Quantification of RNA-FISH signal for early DV patterning genes

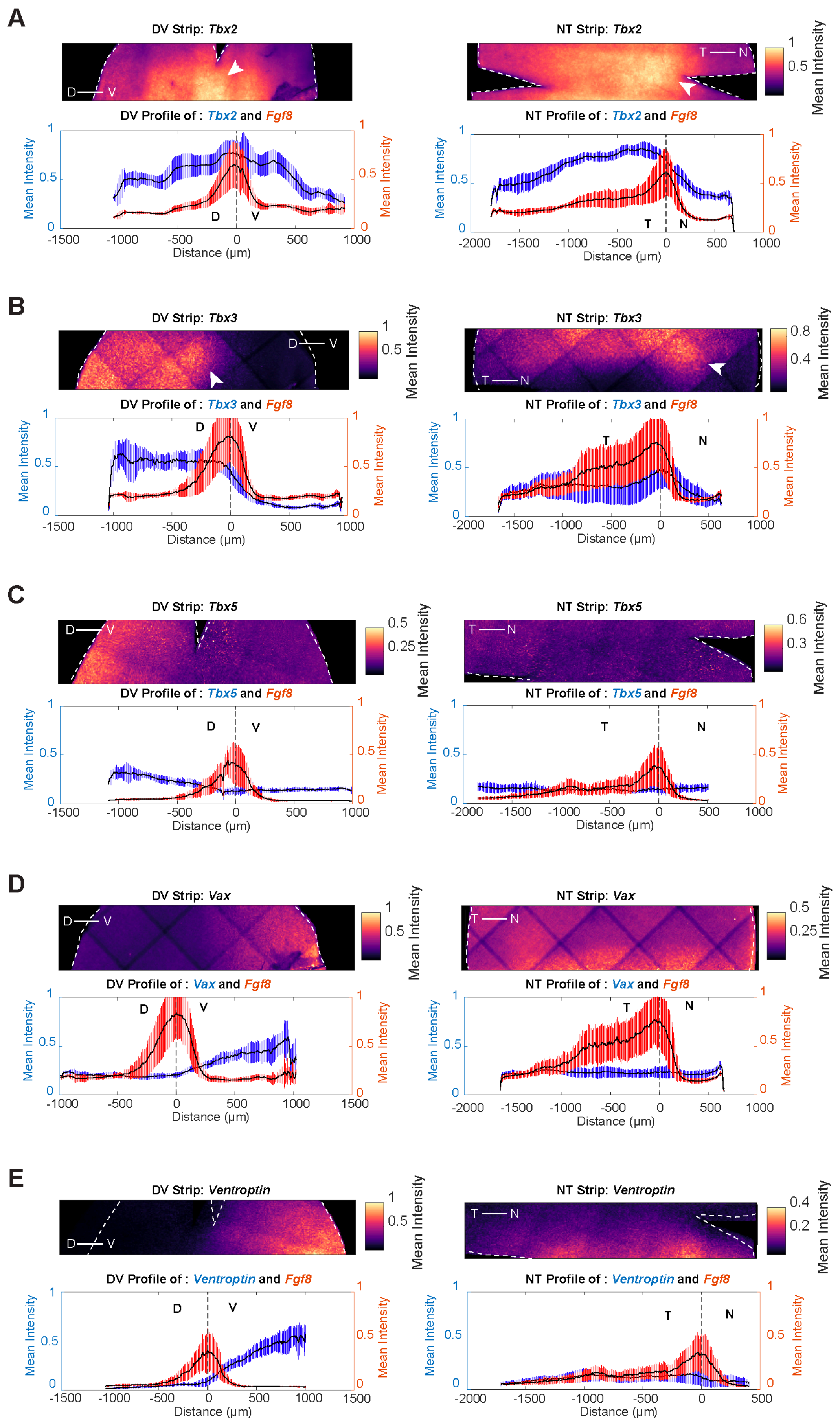

### Supplemental Figure 5

Supplementary Figure 5. Expression domains of early DV patterning genes relative to *Fgf8* at HH25/26

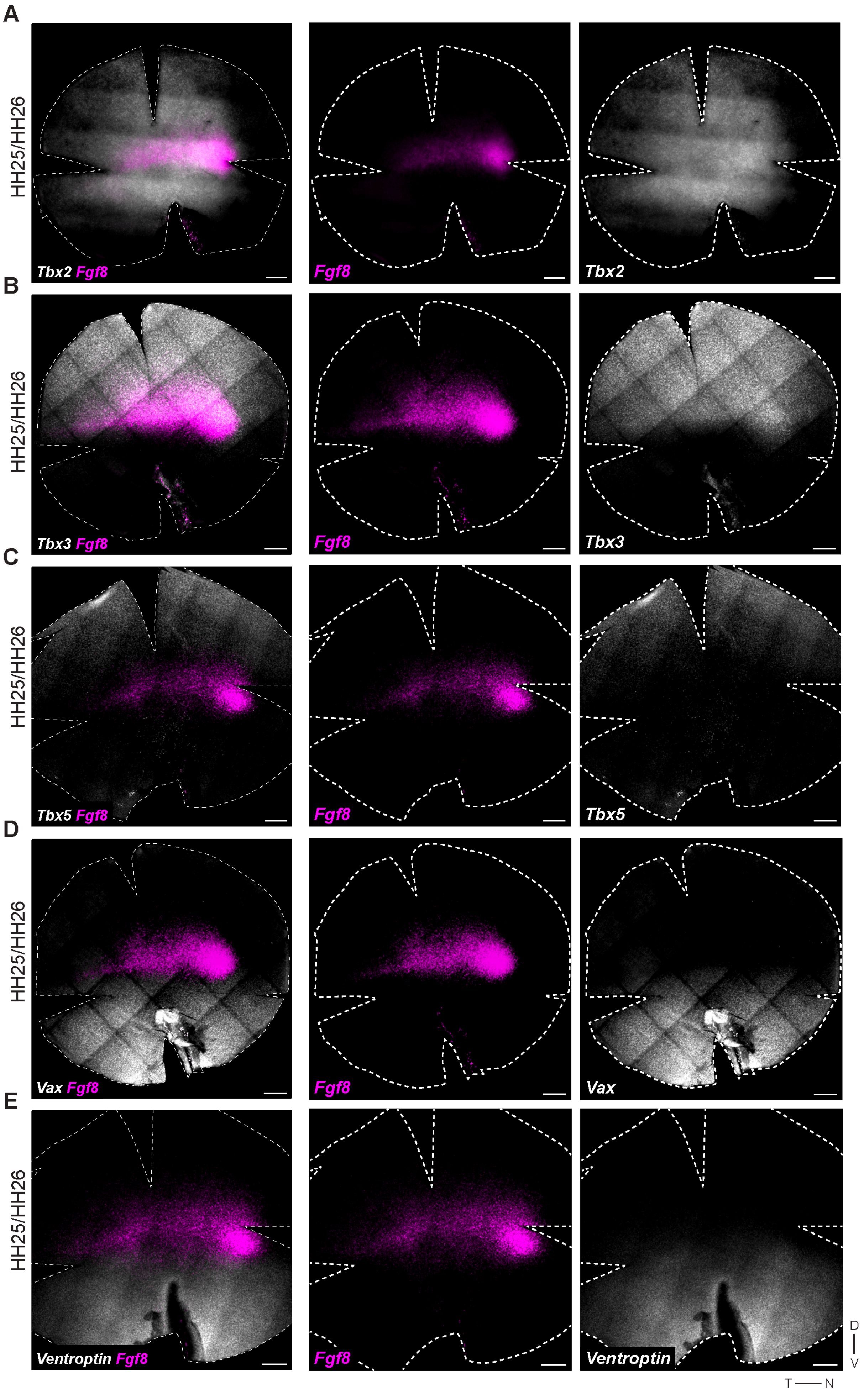

### Supplemental Figure 6

**Supplementary Figure 6. Expression domains of early DV patterning genes relative to Fgf8 at HH28/29**

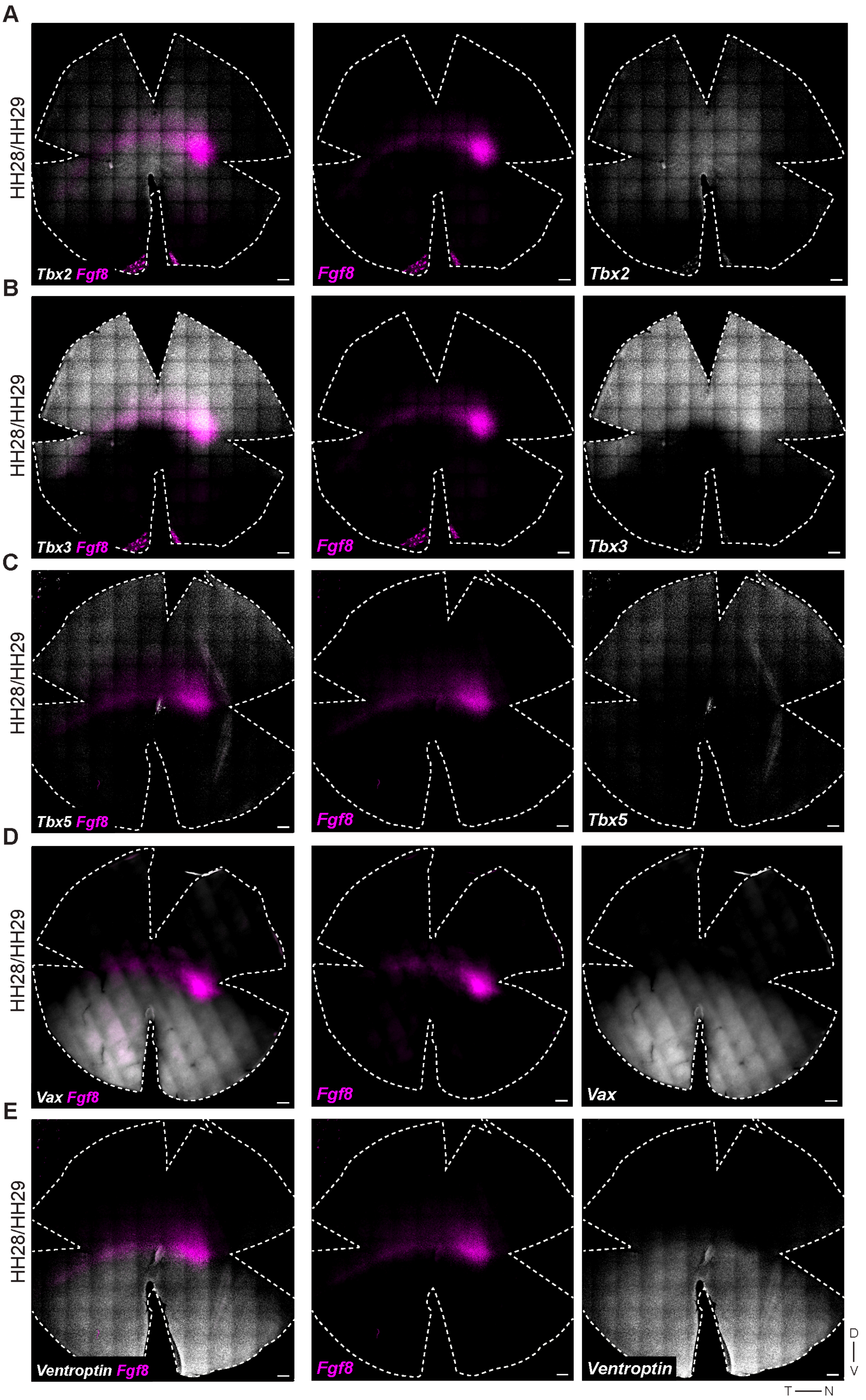

### Supplemental Figure 8

#### Supplementary Figure 8. Quantification of RNA-FISH signal for early NT patterning genes

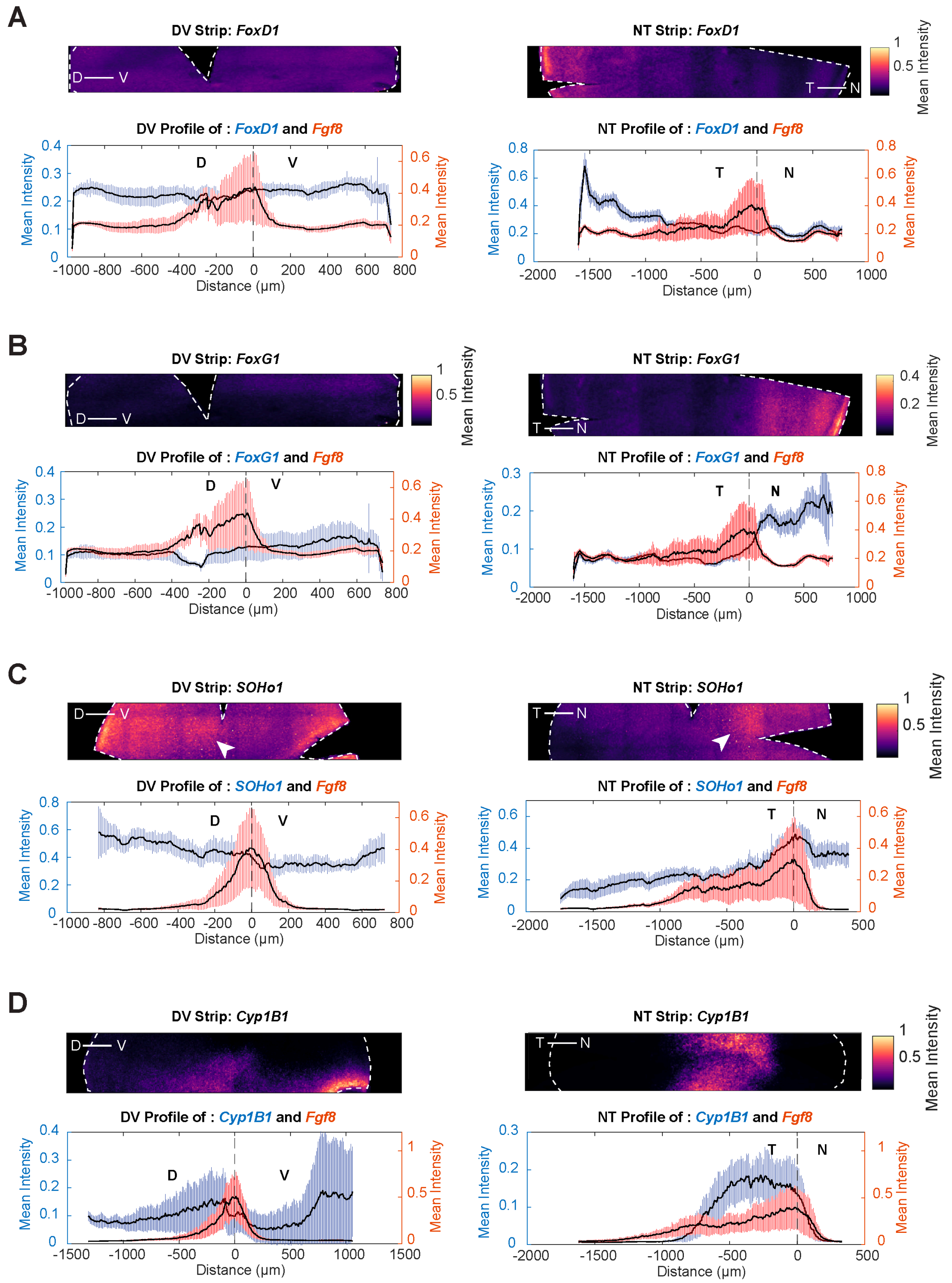

### Supplemental Figure 9

**Supplementary Figure 9. Expression domains of early NT patterning genes relative to *Fgf8* at HH25/26**

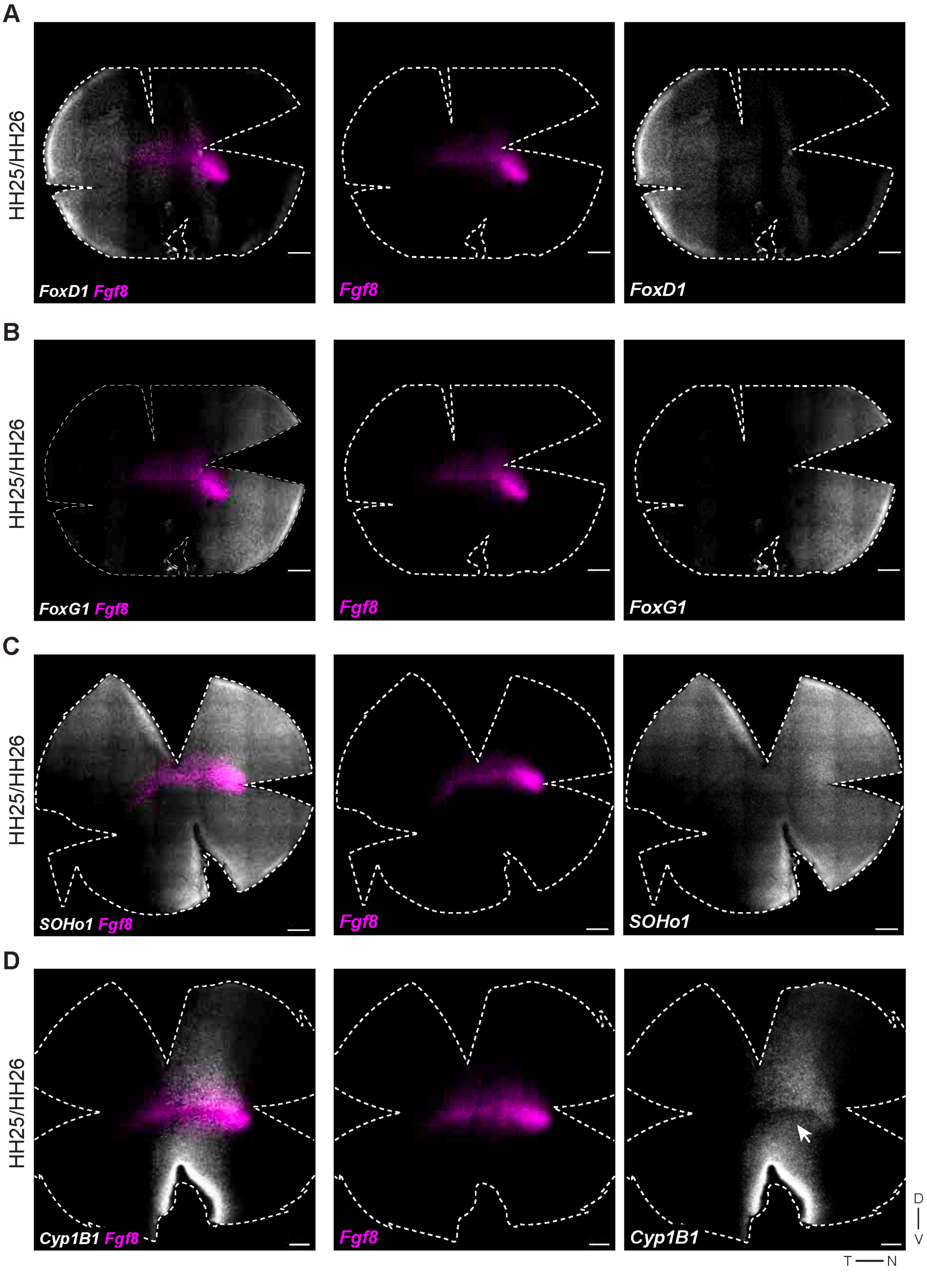

### Supplemental Figure 10

**Supplementary Figure 10. Expression domains of early NT patterning genes relative to *Fgf8* at HH28/29**

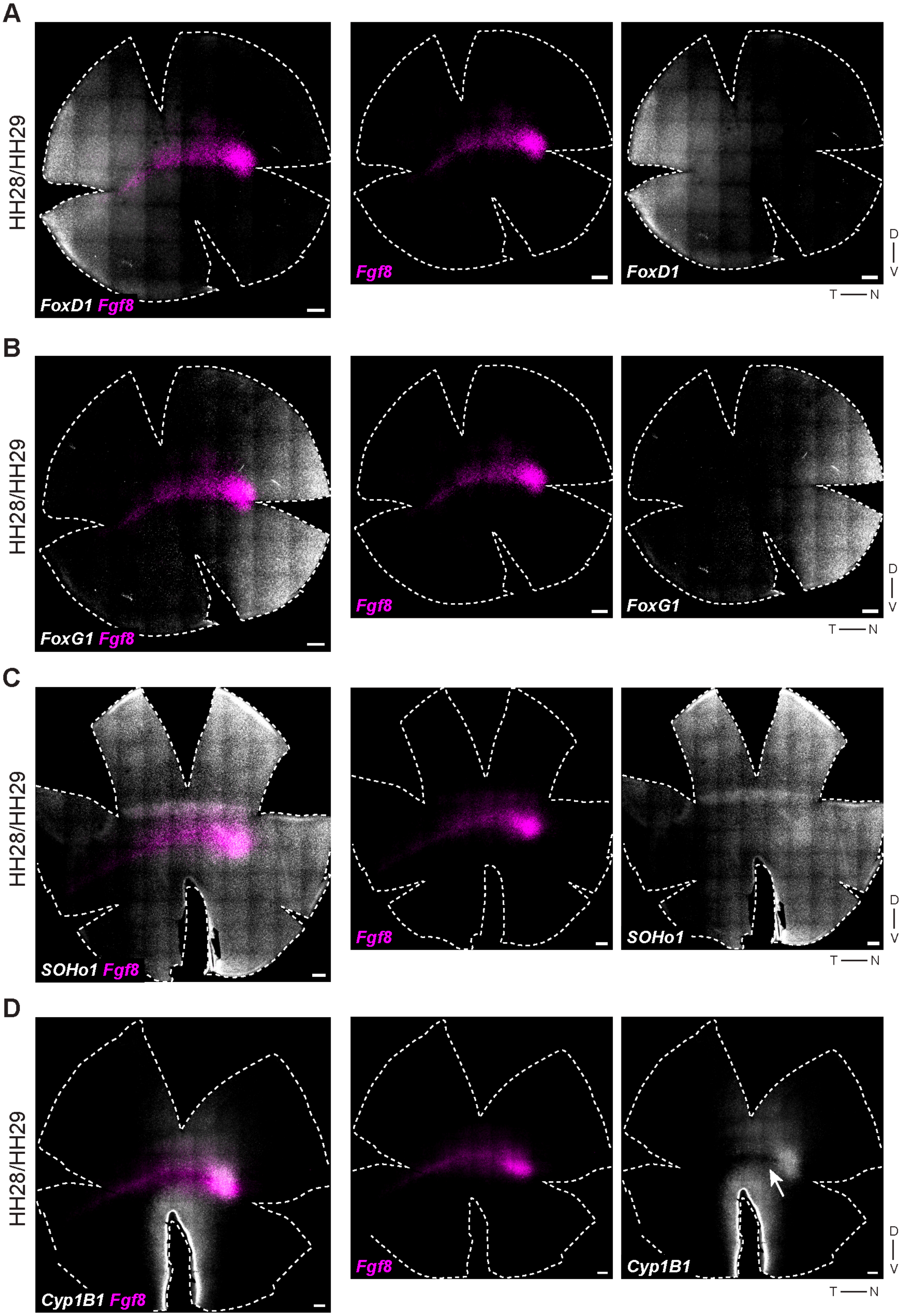

### Supplemental Figure 11

##### Supplementary Figure 11. Quantification of RNA-FISH signal for *Bmp2* and *Visinin*

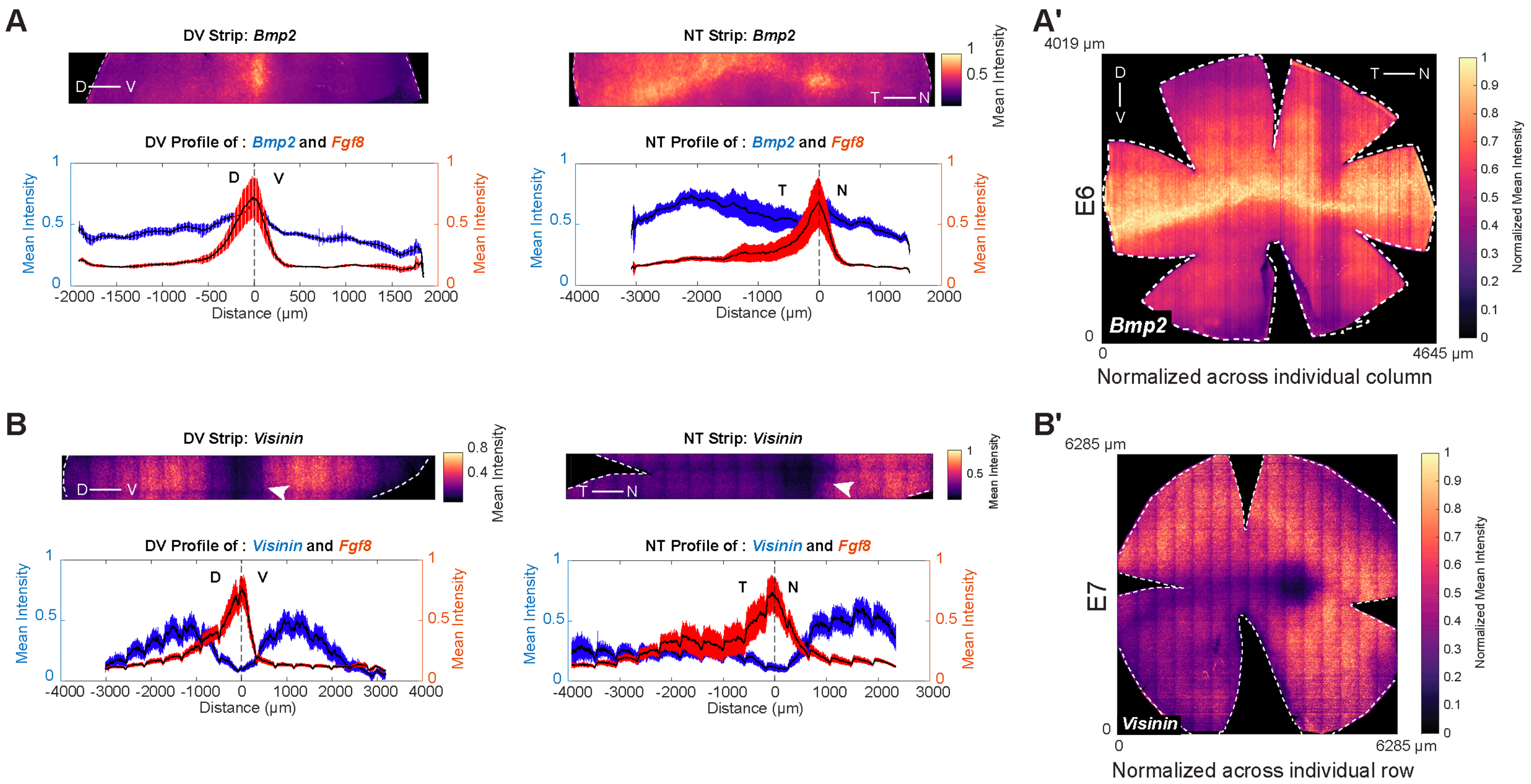

### Supplemental Figure 12

#### Supplementary Figure 12. Processing of scRNA-seq datasets from developing chick retina

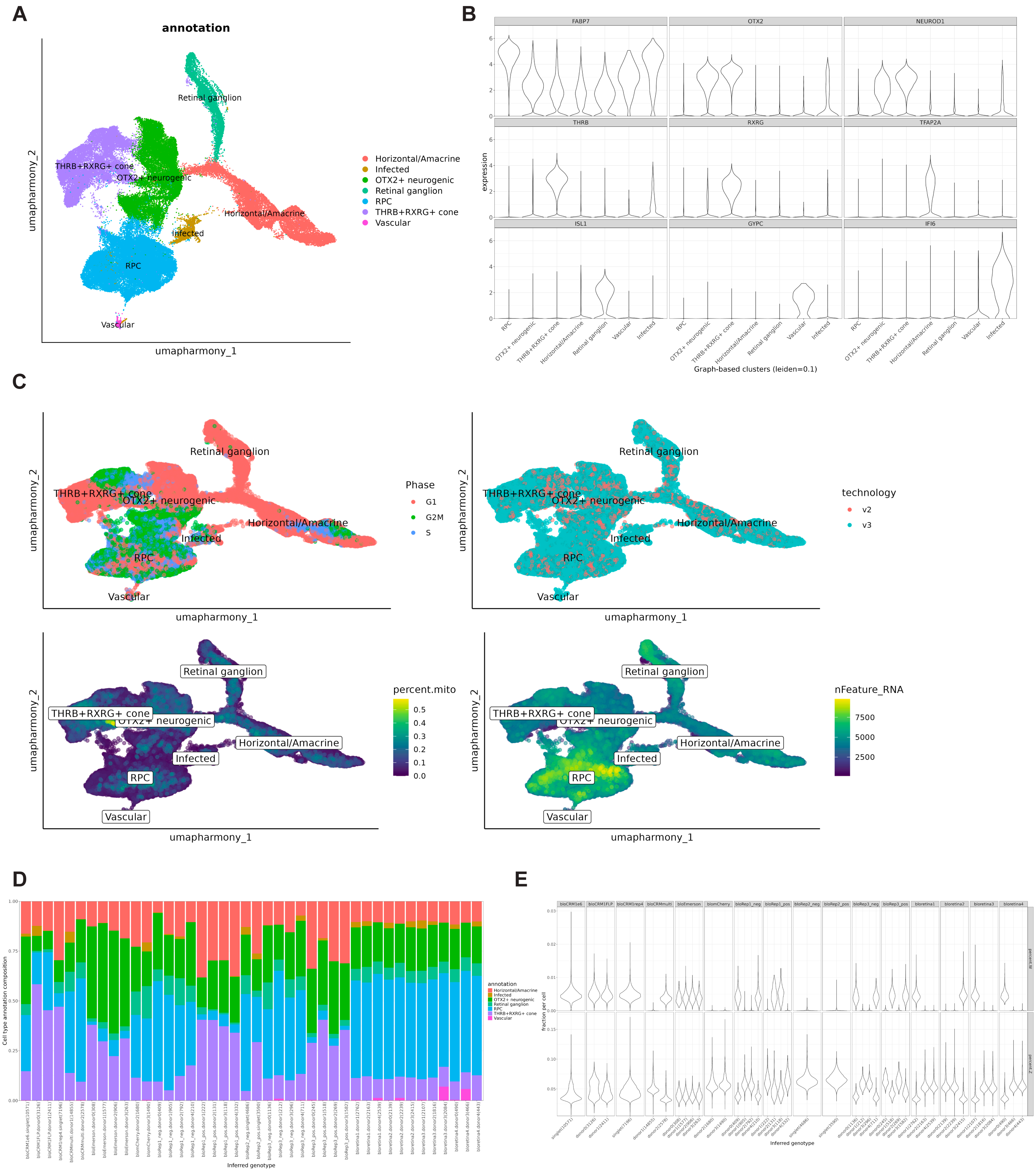

### Supplemental Figure 15

##### Supplementary Figure 15. Grid size sensitivity analysis for 2D topographic map reconstruction

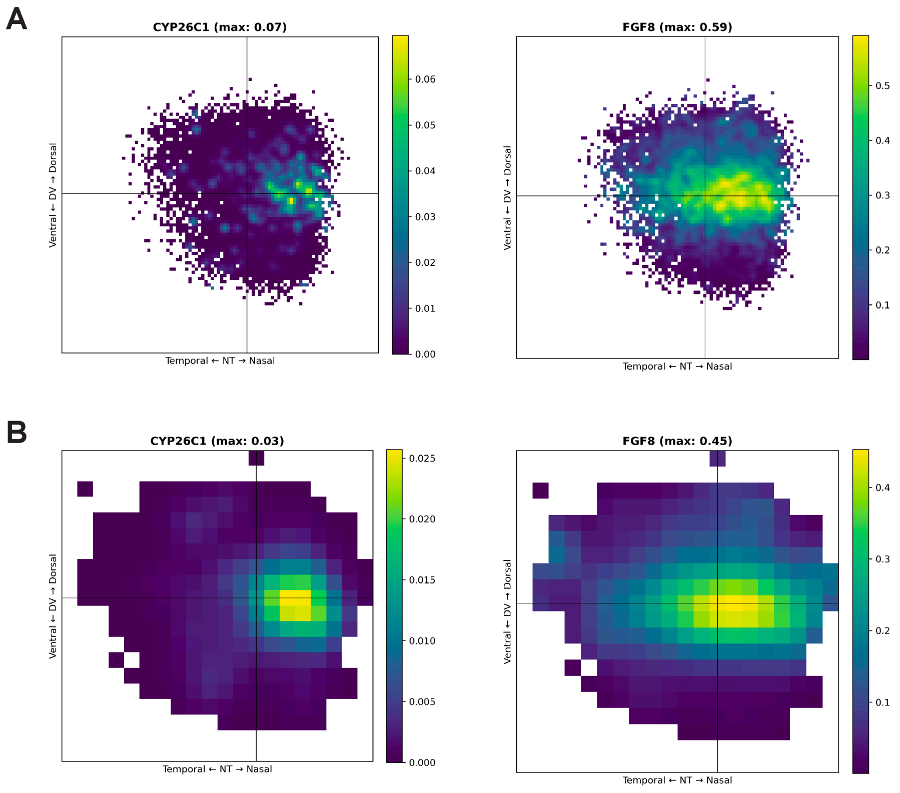

### Supplemental Figure 20

#### Supplementary Figure 20. 2D topographic maps of Fgf family members in mouse and human retinal RPCs

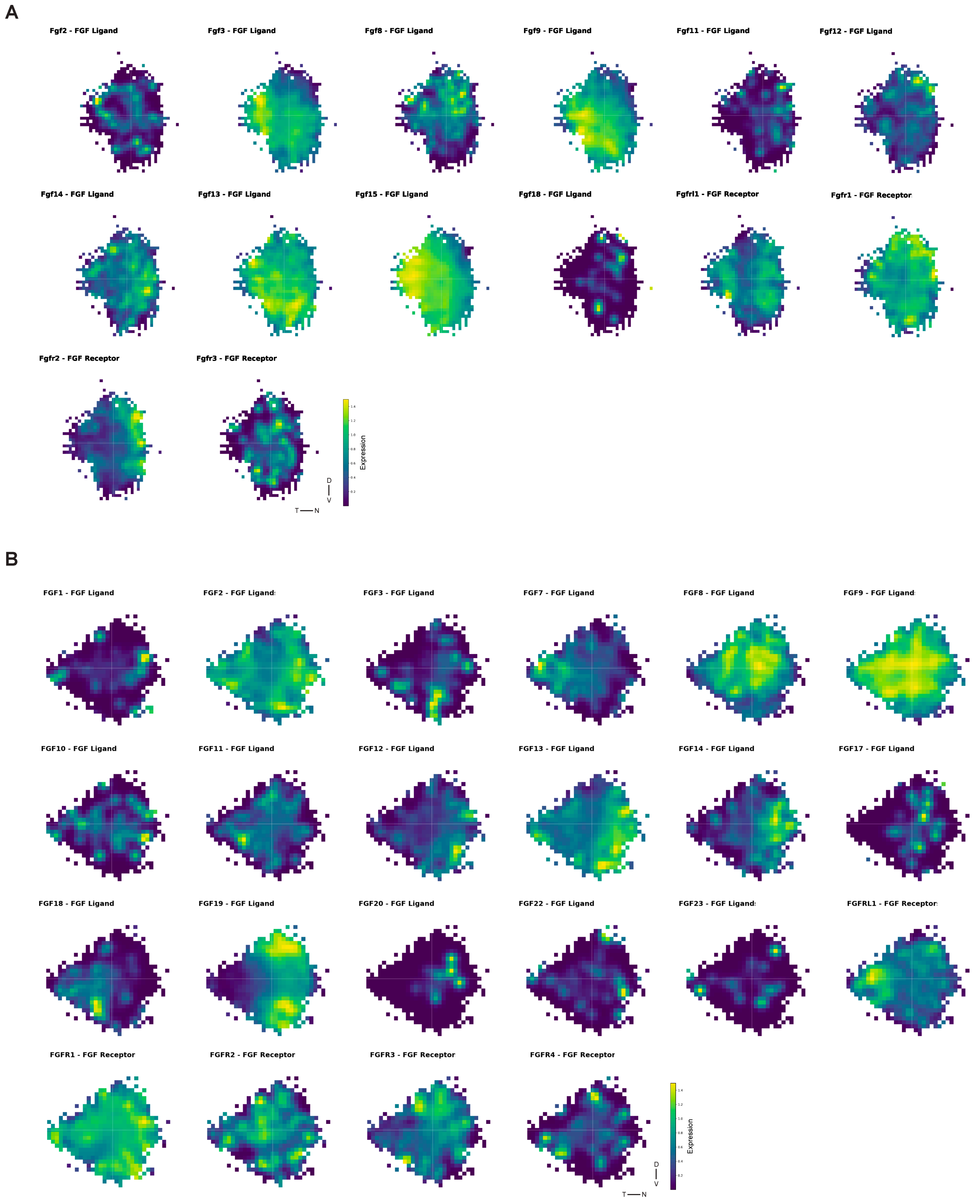
