## Supplemental Figure 3 for "Integration of in situ hybridization and scRNA-seq data provides a 2D topographical map of the developing retina across species"

Supplementary Figure 3. Quantification of RNA-FISH signal for *Fgf8* and RA pathway genes in retinal sections

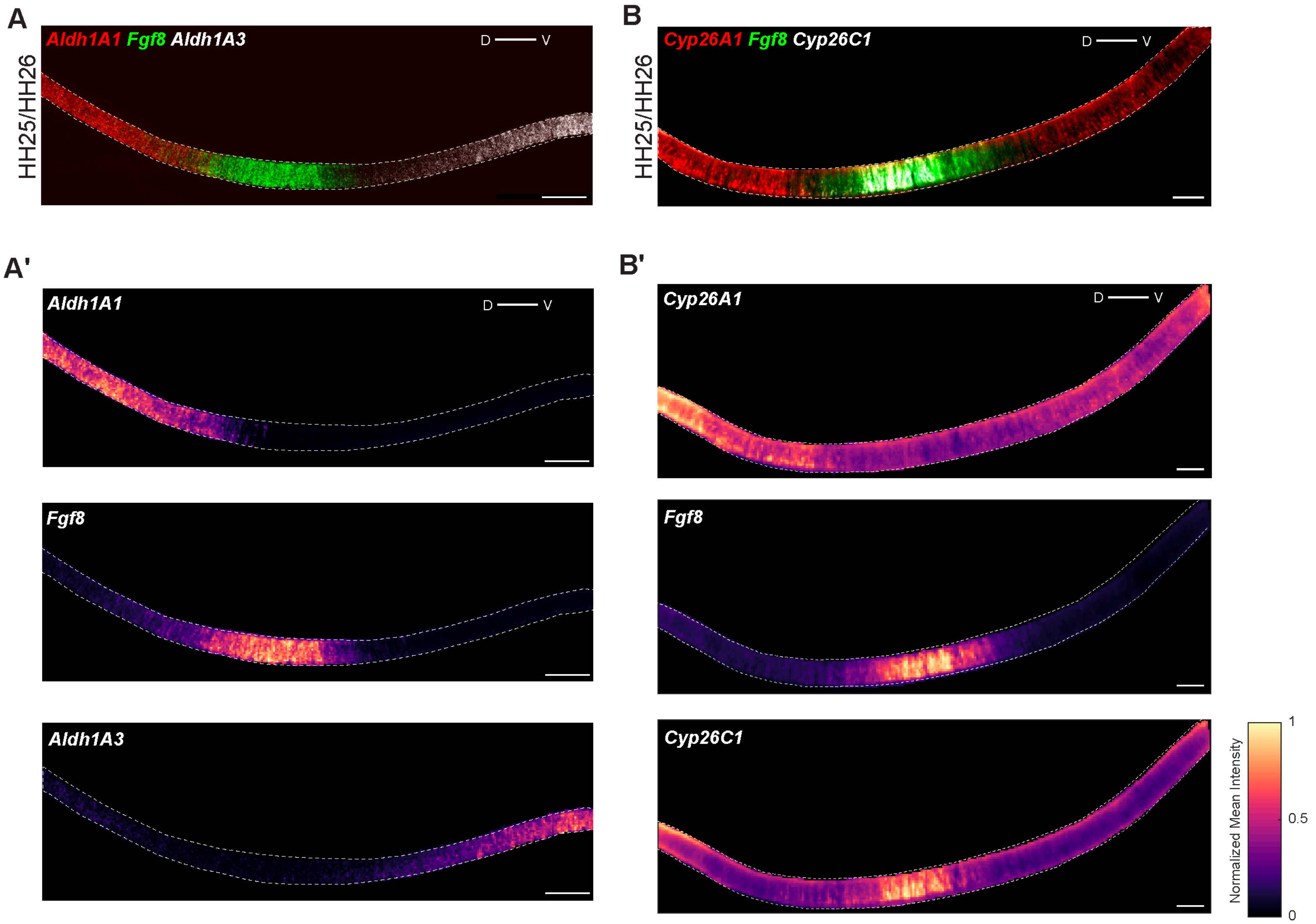
