## Supplemental Figure 7 for "Integration of in situ hybridization and scRNA-seq data provides a 2D topographical map of the developing retina across species"

**Supplementary Figure 7. Expression domains of Eph/Ephrin signaling genes in the developing chicken retina**

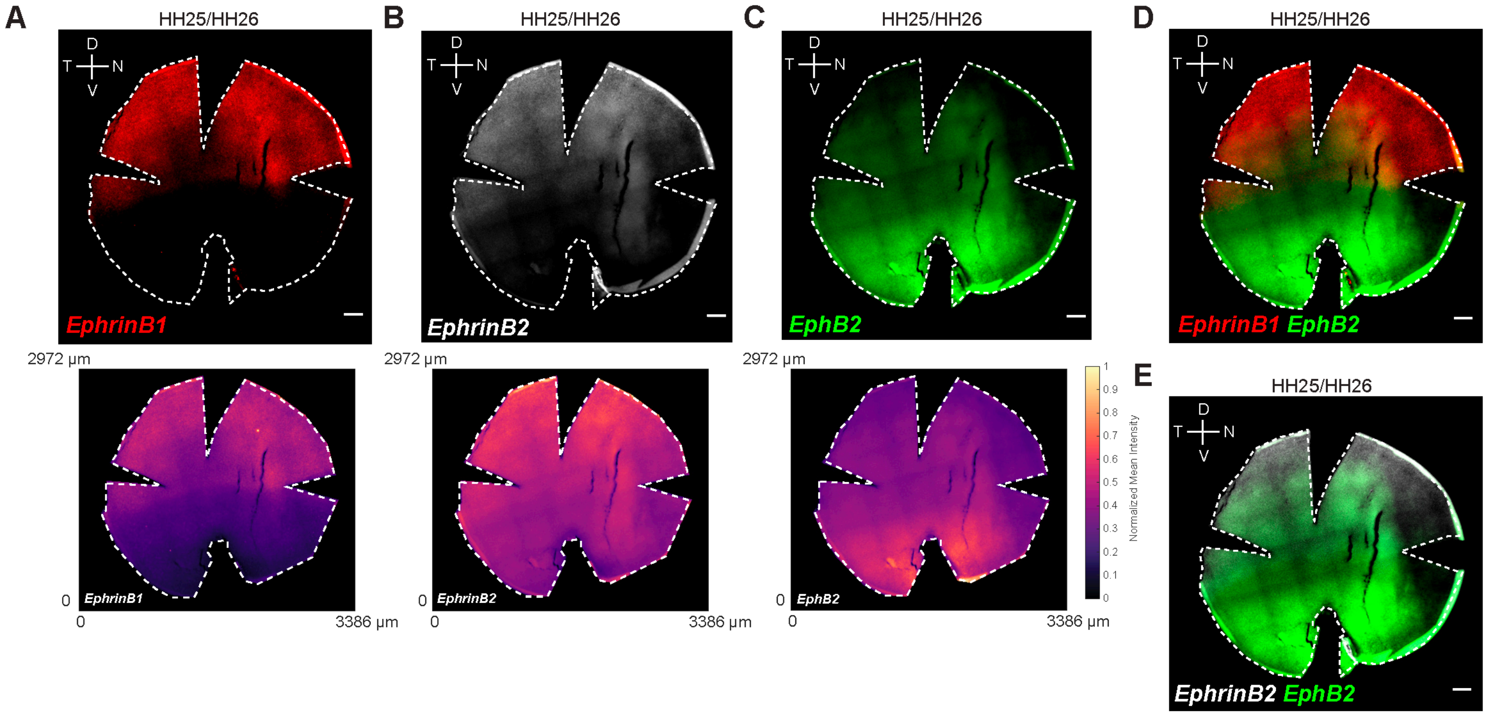
