## Supplemental Figure 13 for "Integration of in situ hybridization and scRNA-seq data provides a 2D topographical map of the developing retina across species"

### Supplementary Figure 13. Generation of DV score using single-cell transcriptomes from the developing chick retina

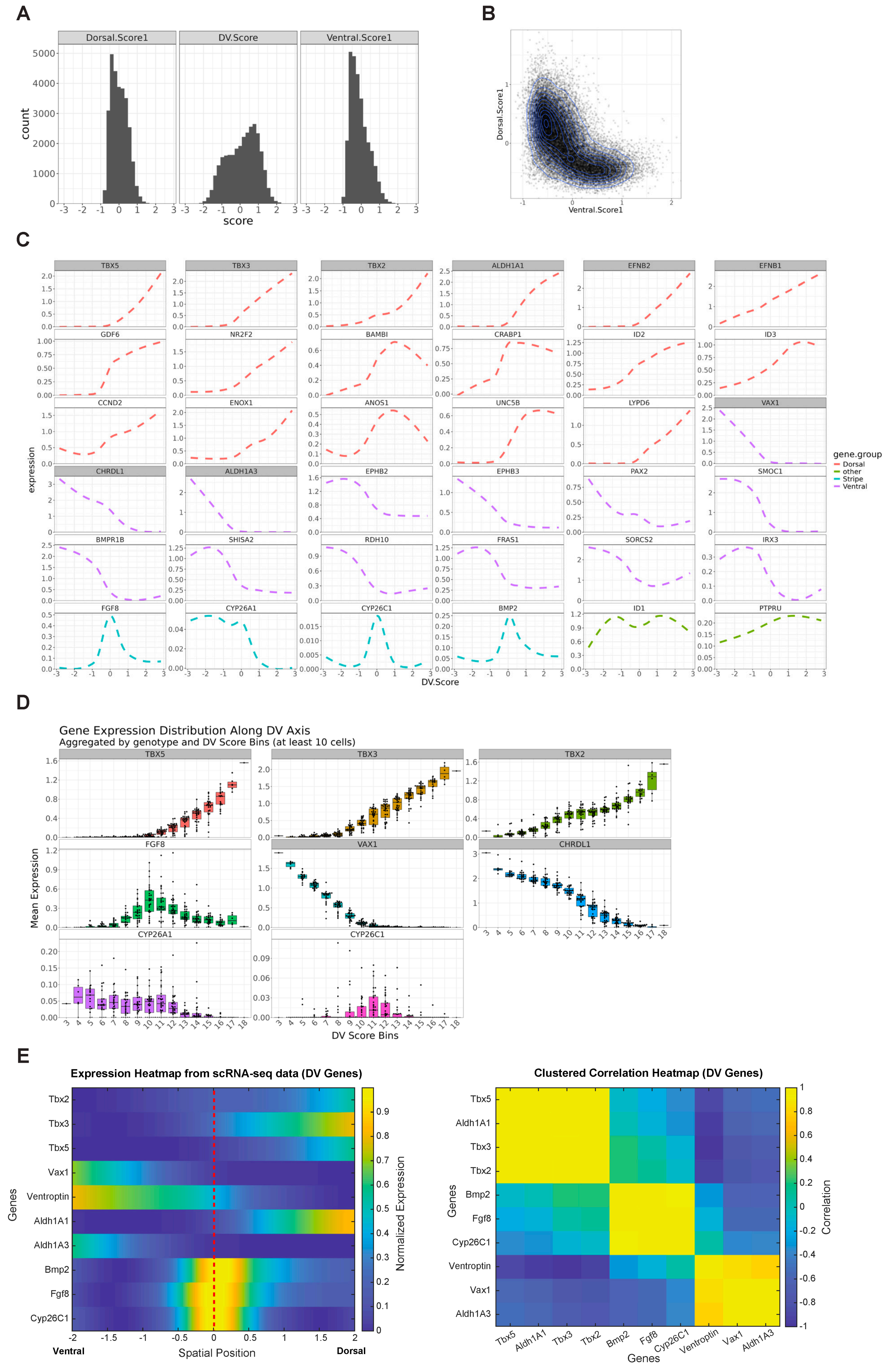
