## Supplemental Figure 16 for "Integration of in situ hybridization and scRNA-seq data provides a 2D topographical map of the developing retina across species"

### Supplementary Figure 16. Identification of genome-wide spatial expression patterns in the chick retina

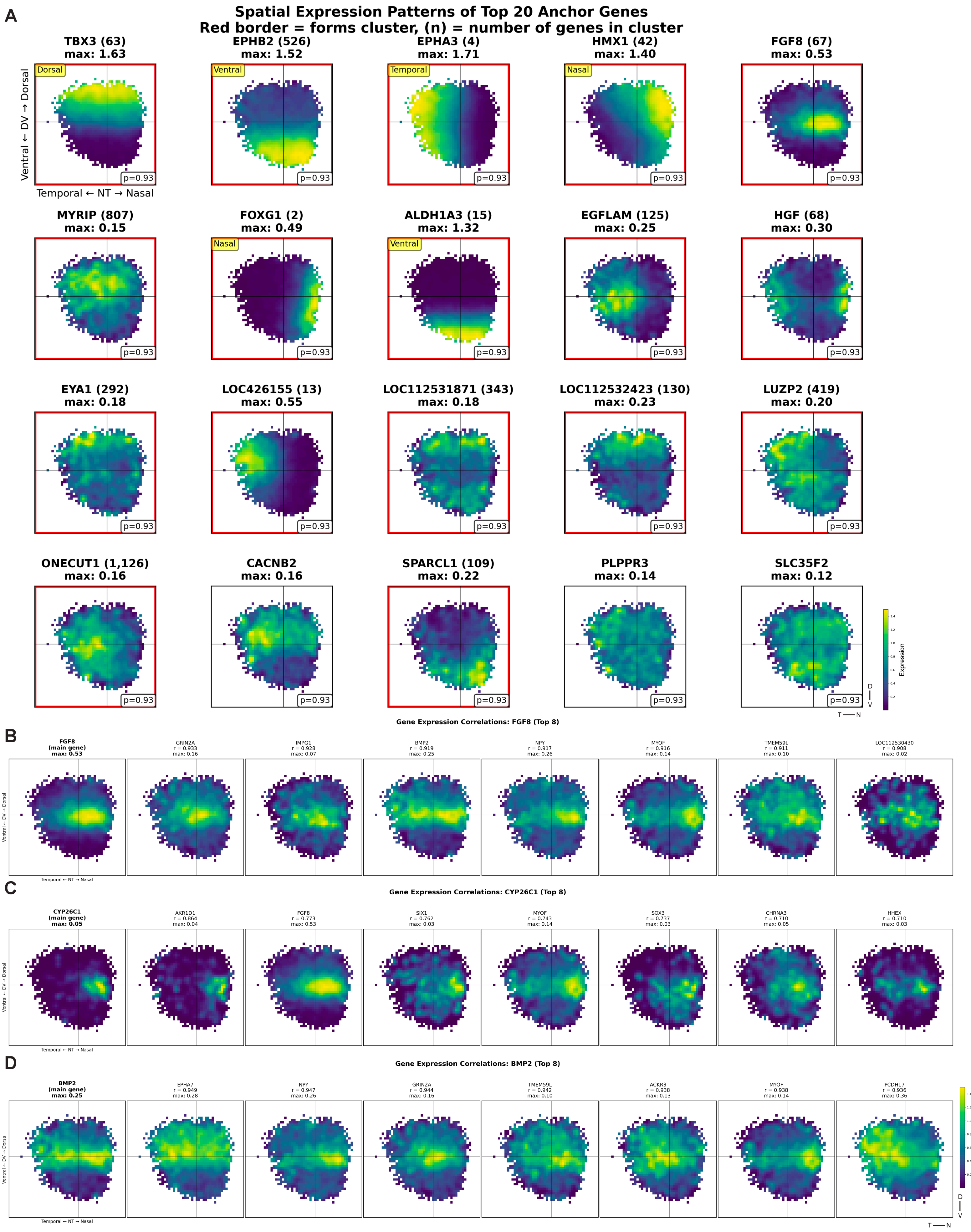
