## Supplemental Figure 17 for "Integration of in situ hybridization and scRNA-seq data provides a 2D topographical map of the developing retina across species"

### Supplementary Figure 17. Spatial expression patterns of genes associated with Fgf, Bmp and RA signaling pathways in the chick retina

A

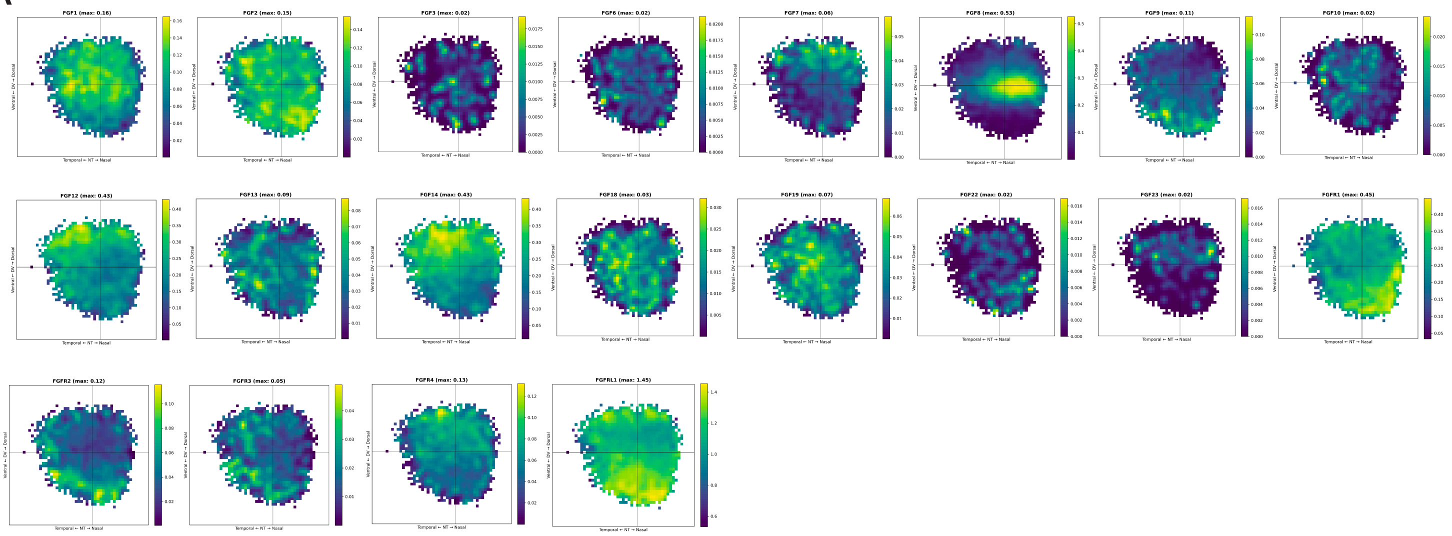


B

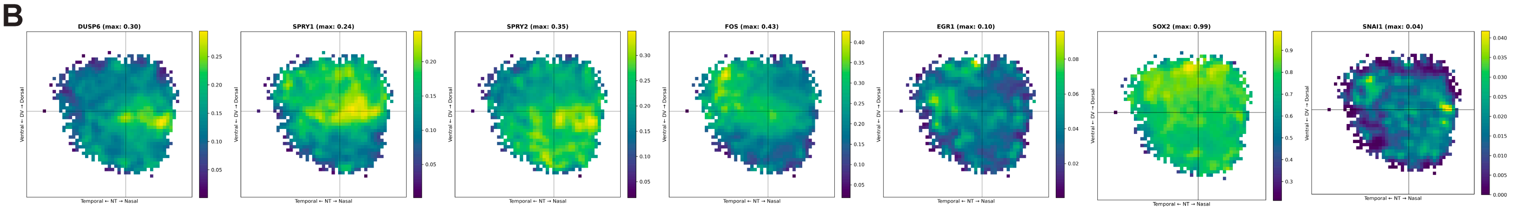


C

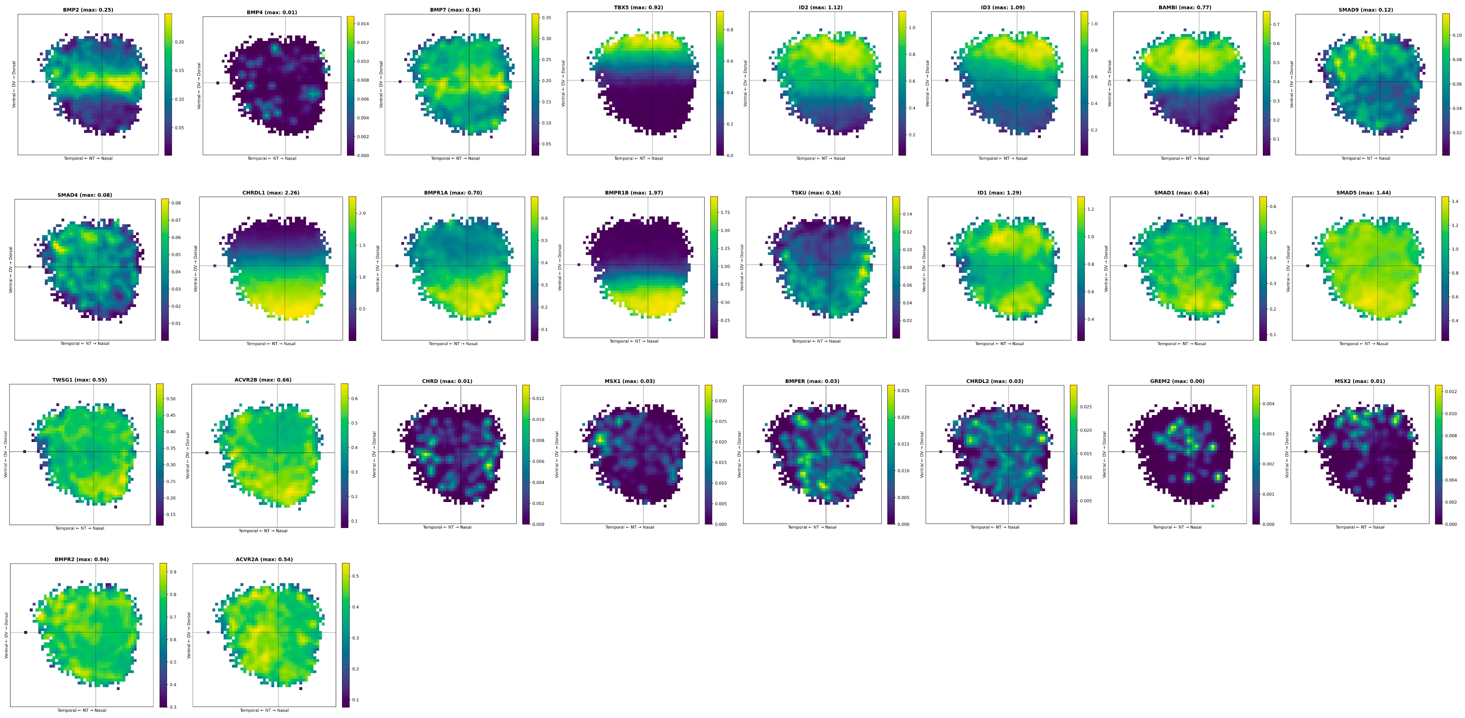


D

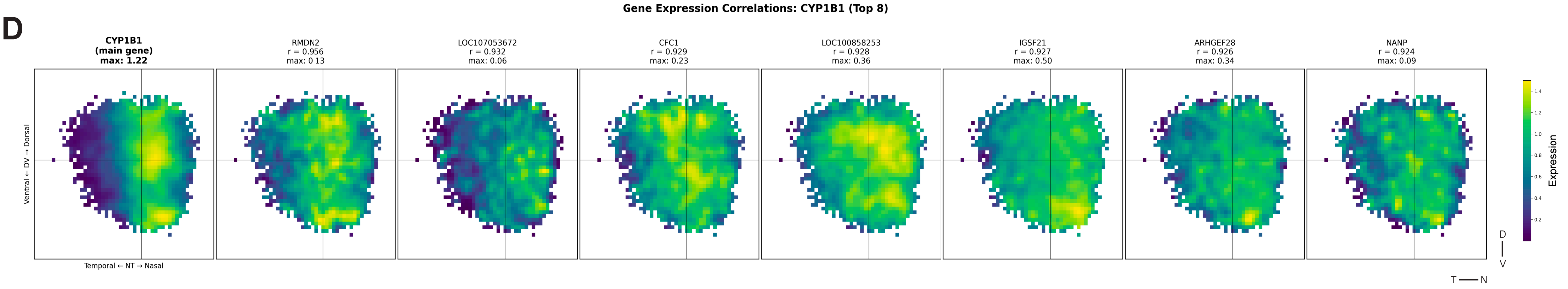
