## Supplemental Figure 18 for "Integration of in situ hybridization and scRNA-seq data provides a 2D topographical map of the developing retina across species"

### Supplementary Figure 18. Generation of DV and NT scores using single-cell transcriptomes from developing mouse and human retinas

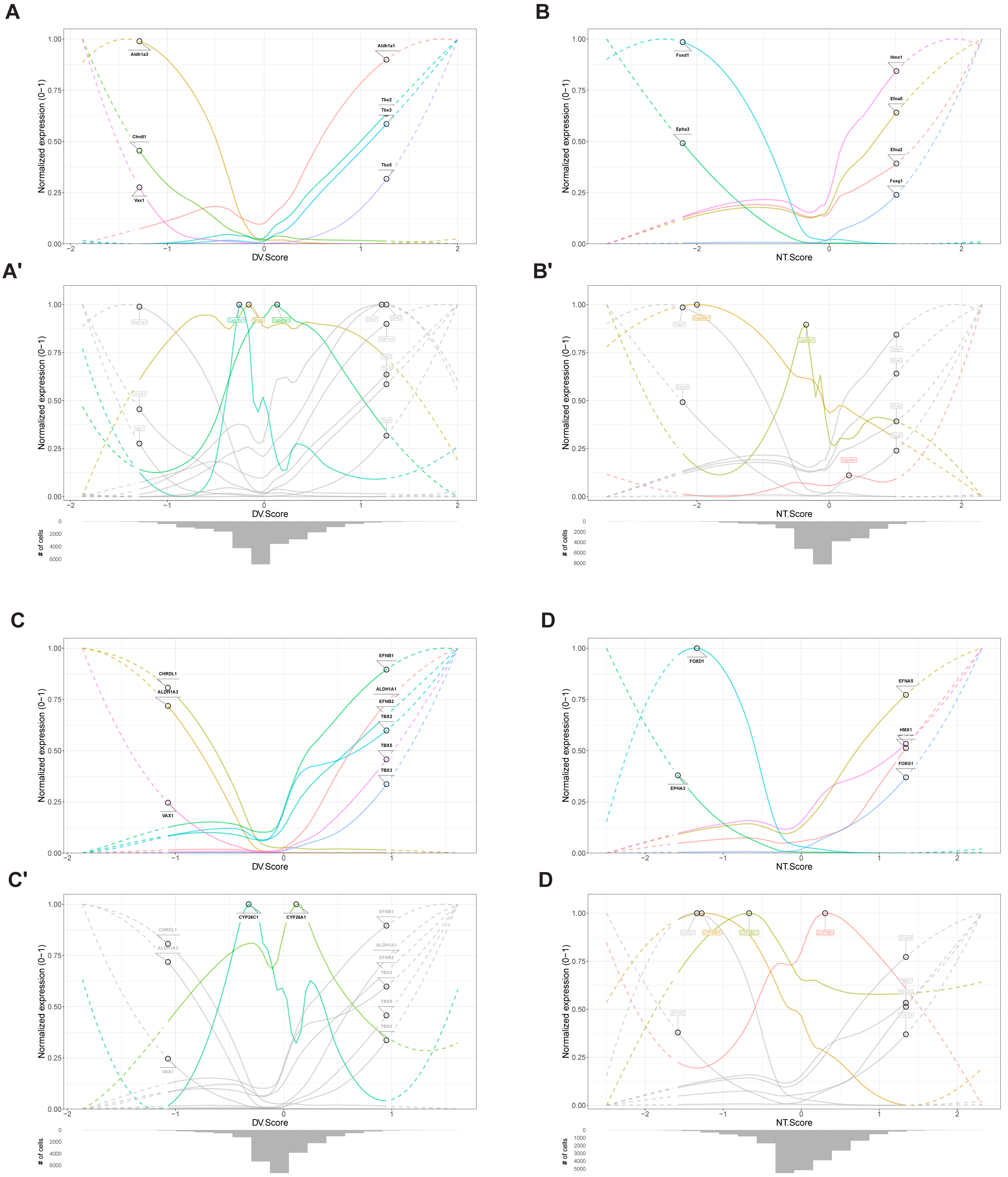
