## Supplemental Figure 19 for "Integration of in situ hybridization and scRNA-seq data provides a 2D topographical map of the developing retina across species"

**Supplementary Figure 19. Identification of clusters with spatial expression patterns in mouse and human retinas**

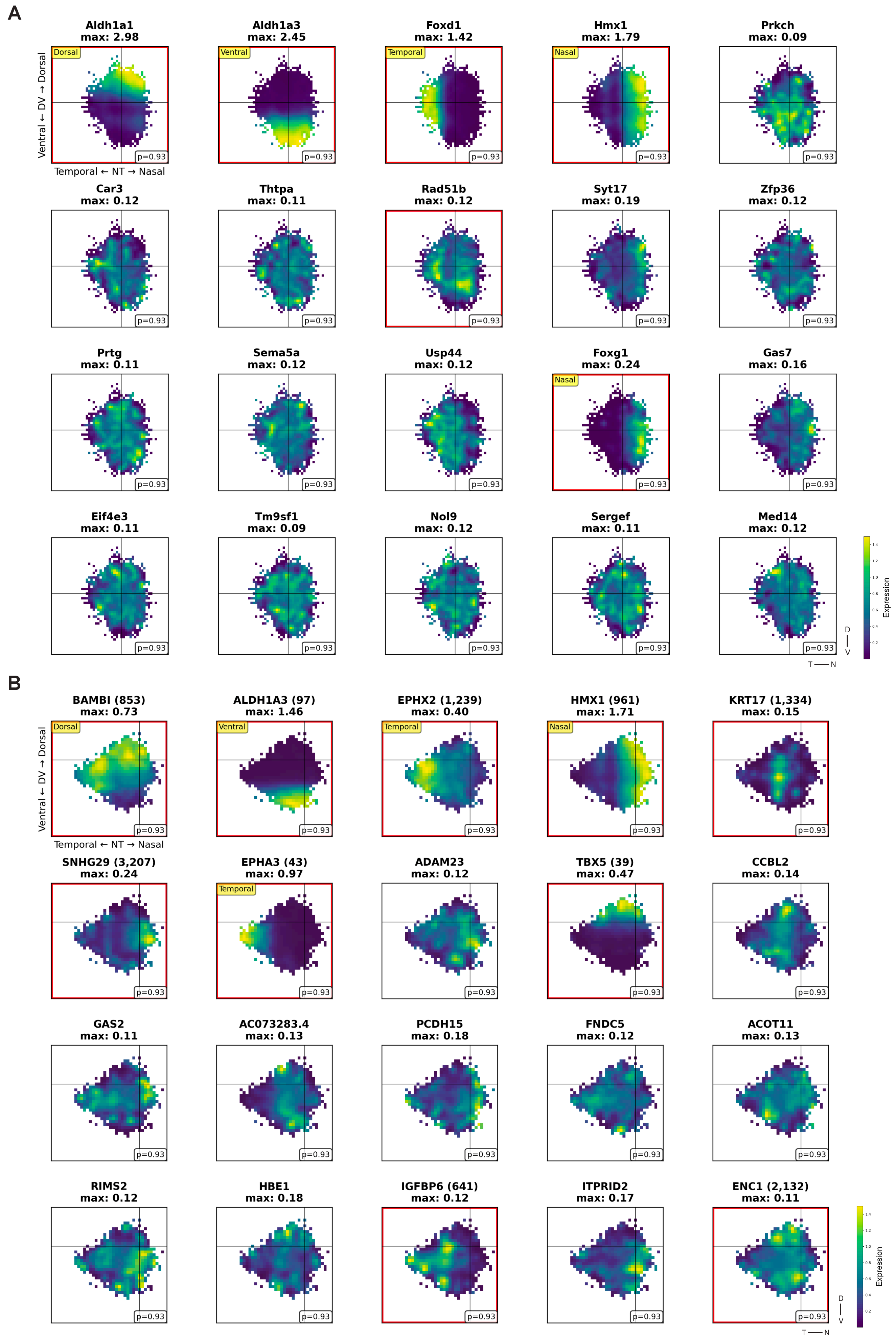
