## Supplemental Figure 21 for "Integration of in situ hybridization and scRNA-seq data provides a 2D topographical map of the developing retina across species"

### Supplementary Figure 21. 2D topographic maps of genes related to *Fgf8* signaling pathway in mouse and human retinal RPCs

**A**

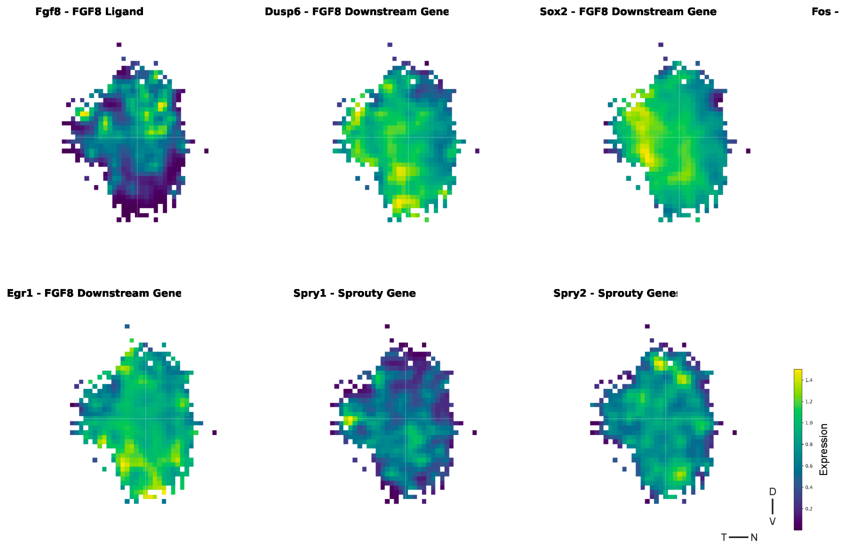


**B**

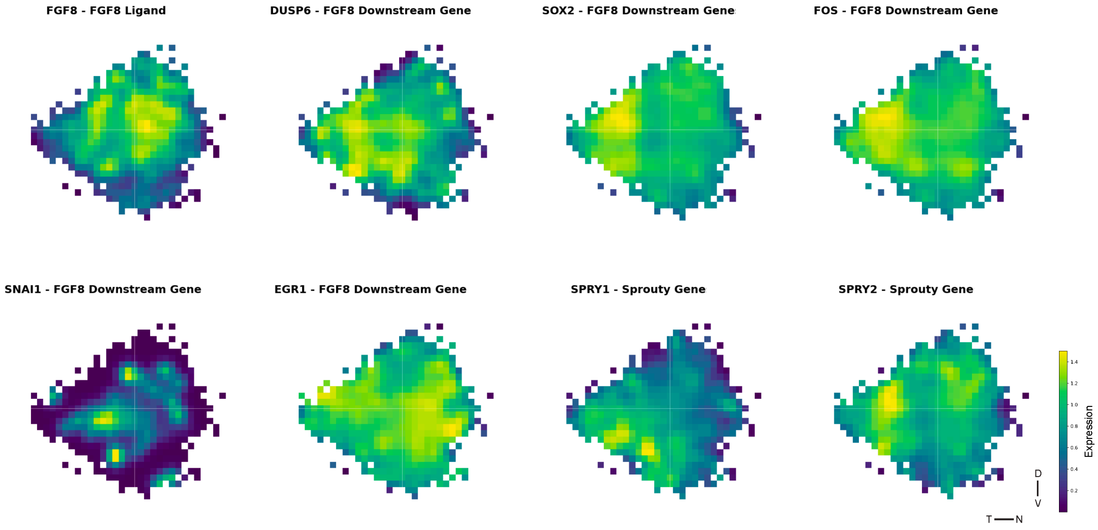
