## Supplemental Figure 22 for "Integration of in situ hybridization and scRNA-seq data provides a 2D topographical map of the developing retina across species"

**Supplementary Figure 22. 2D topographic maps of BMP signaling pathway components in developing mouse and human retinas**

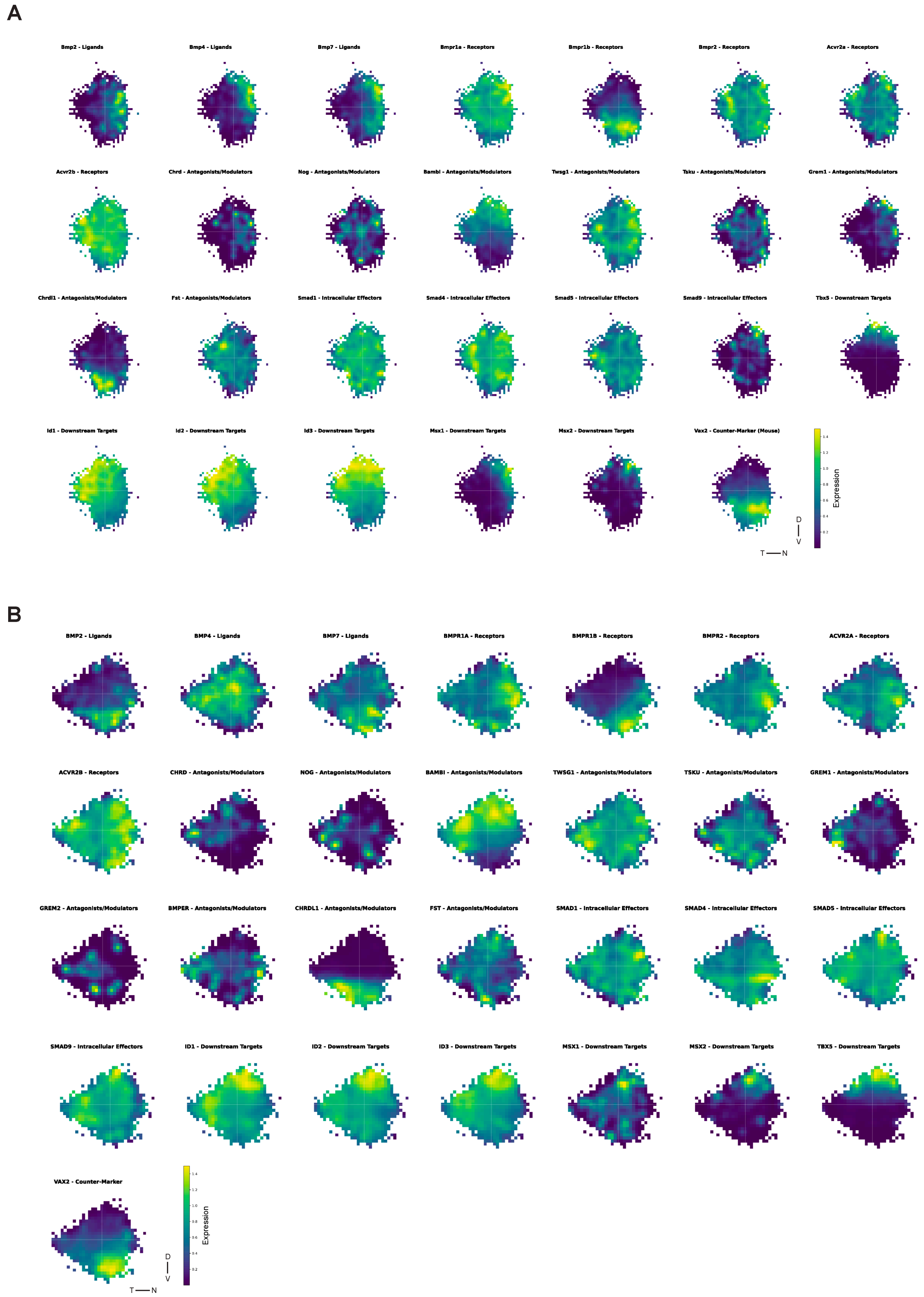
