## Supplemental Figure 23 for "Integration of in situ hybridization and scRNA-seq data provides a 2D topographical map of the developing retina across species"

### Supplementary Figure 23. Differential gene expression analysis across different regions in the human retina

## A

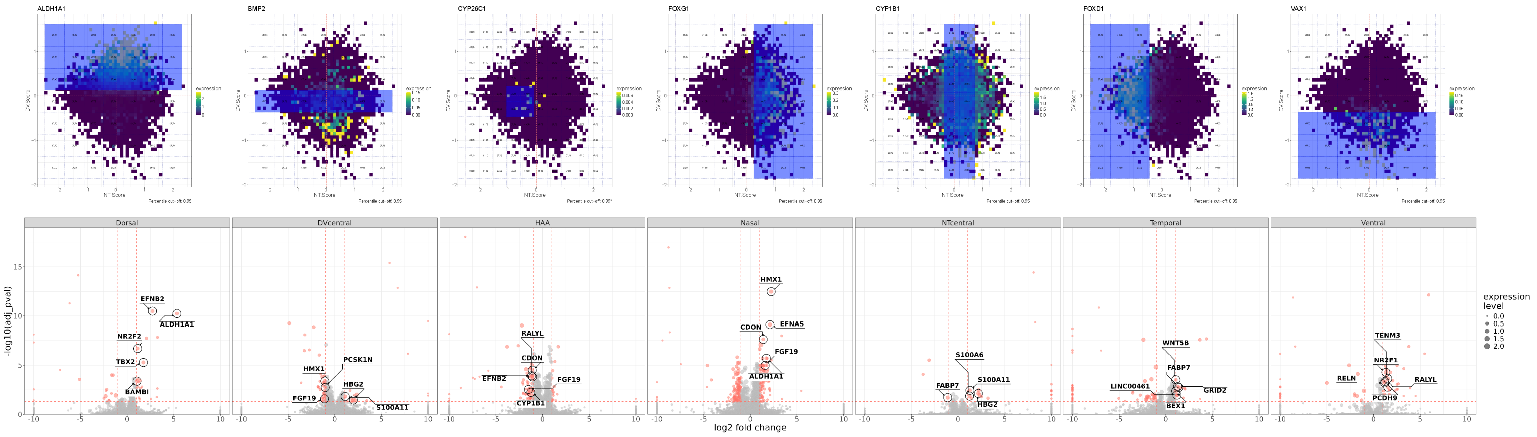
